# Blended Length Genome Sequencing (blend-seq): Combining Short Reads with Low-Coverage Long Reads to Maximize Variant Discovery

**DOI:** 10.1101/2024.11.01.621515

**Authors:** Ricky Magner, Fabio Cunial, Sumit Basu, Ron Paulsen, Scott Saponas, Megan Shand, Niall Lennon, Eric Banks

## Abstract

We introduce blend-seq, a workflow for combining data from traditional short-read sequencing pipelines with low-coverage long reads, to improve variant discovery for single samples without the full cost of high-coverage long reads. We demonstrate that with only 4x long-read coverage augmenting 30x short reads, we can improve SNP discovery across the genome, exceeding performance beyond even high-coverage short reads (60x). For genotype-agnostic discovery of structural variants, we see a threefold improvement in recall while maintaining precision by using the low-coverage long reads on their own, and show how we can improve genotyping accuracy by adding in the short-read data. In addition, we demonstrate how the long reads can better phase these variants, incorporating long-context information in the genome to substantially outperform phasing with short reads alone. Our experiments highlight the complementary nature of short- and long-read technologies: the former contributing higher depth for genotyping and the latter better resolution of larger events or those in difficult regions.

## Introduction

While market-leading short-read sequencing technologies (such as Illumina’s Novaseq X) are excellent at capturing small variations (SNPs and INDELs) in most of the human genome, they are far less effective both for hard-to-map regions as well as for structural variants (SVs, ≥ 50bp). Longread technologies, such as those from Pacific Biosciences (PacBio), with reads of 15-20kb, and Oxford Nanopore Technologies (ONT), with reads of up to 2Mb, on the other hand, have shown remarkable success in both of these areas; however, the significantly higher cost has left them out of reach for many applications. In this paper, we introduce *Blended Length Genome Sequencing* (**blend-seq**), a workflow for combining traditional short-read pipelines with lowcoverage long reads, and demonstrate how we can make substantial gains in both short and structural variant calling for single samples as well as phasing without the full cost of high-coverage long-read sequencing. Although still 2.3x the cost of short-read sequencing alone, blend-seq is around half the cost of long-read genome sequencing making it a natural economic middle ground with strong performance. Details of the cost estimate analysis are provided in an appendix (Section A).

Blend-seq is flexible with respect to choices and coverage levels of each sequencing technology. A pragmatic setting given existing practices is to augment the typical 30x coverage of Illumina short reads with 4x coverage of PacBio long reads. For short variants, we show that with this setting we can combine read types to improve SNP recall (while maintaining precision) not only in hard-to-map regions but across the entire genome, even compared to very high coverage short-read performance, which levels off well before 60x. For SVs, we show that we can improve recall by a factor of three across the genome, and by even more for variants overlapping with the exome, simply by using the lowcoverage long reads on their own. For settings where the genotype of structural variants matters, we show even greater gains by combining short and long reads using graph-based approaches. For phasing, we show performance rivaling full coverage (30x) PacBio pipelines.

The implications for the SNP results are immediate: the vast majority of known genetic diseases are identified by these single nucleotide polymorphisms, so our improvements to SNP recall have the potential to impact clinical diagnosis. Such gains apply to the discovery of new variant associations as well: greater recall implies a greater ability to associate SNPs with phenotypic profiles in biobank or cohort data. As such this improvement in SNP performance has the potential of both improving existing diagnostic procedures and the discovery of new diagnostics.

At the same time, SNPs tell only part of the story. From population genomics (1) to understanding disease mechanics (2, 3), there is a need to move beyond short variants and consider a wider set of causes for genetic disease (4). To date, the best way to address these larger variants has been through long-read technologies. These far longer reads can span much larger variants, while simultaneously mapping more accurately to the correct locations in the human genome, particularly in low-complexity regions.

Beyond the problem of finding variants, for diploid organisms there is also the need for phasing – determining which variants co-occur on the same chromosome. Concrete applications for phasing have been well-documented, from understanding variant contribution to phenotypic expression and disease states (5), to accurate HLA typing, which is associated with improved outcomes for transplant recipients (6). Short reads alone are also known to be limited in their ability to phase variants to the quality required for pharmacogenetic profiling (see e.g. (7)). Accurate phasing relies both on high precision in calling variants and on having overlapping spans of reads covering multiple variants. While short reads excel at the former problem, the latter makes phasing particularly challenging for short reads, whereas long reads have much greater capability to span multiple variants. By balancing these two advantages, blend-seq provides an affordable option for high-quality phased variants.

We demonstrate these gains with three approaches, varying in complexity and degree of integration between short- and long-read pipelines. For improvements in SNP discovery, we leverage the mapping capabilities of low-coverage long reads, then break these reads up into “virtual” short reads that can be combined with true short reads in the state-of-the-art DRAGEN calling pipeline. For SV discovery, we simply use the low-coverage long reads to call structural variants, then use the short reads to better genotype them in some settings. For phasing, we leverage the benefits of blend-seq for calling variants, then use the long reads alone to bridge the variants together, combining the strengths of both read types to substantially exceed the performance of short reads alone. Our benchmarks focus on the well-known sample HG002, from the Genome in a Bottle (GIAB) standard at NIST (8), as it is the only GIAB sample with a Q100 assembly available (9) that can accurately benchmark variants (short and structural) as well as phasing.

Our work is unique in focusing on combining *standard coverage* short reads with *low coverage* long reads with only *in silico* methods. Previous studies have either required high-coverage from both, like a recent offering from Variantyx (10, 11) using 12-15x long reads, or special library preparations to create biochemical correspondences between the data types, like in read clouds, synethetic long reads, and linked reads (12–18). As prices for both types of reads continue to drop, we expect our work to remain relevant in demonstrating the capabilities of cost-effective combinations.

## Results

### Short Variant Calling Performance

In this first subsection, we show that by breaking mapped long reads into synthetic short reads, blend-seq can improve SNP performance statistics, especially in hard-to-map regions, and end with some comments on INDELs.

#### Approach and Data

While short-read pipelines perform well where their read lengths are sufficient to map unambiguously to the genome, their performance suffers when the mapping is no longer unique; conversely, long reads are well-known to be better at mapping uniquely to these more difficult regions. Our approach was to map the long reads on their own, then break each alignment into segments of short-read size (an idea that already appeared e.g. in (19)). These “virtual” short-read alignments were then passed to the short-read calling pipeline along with the alignments of the actual short reads, to create a hybrid BAM which was then used for variant calling.

We performed our experiments with varying levels of shortread coverage, from 10x to 60x, simulated by downsampling a single set of reads (Illumina NovaSeq X) for a single sample (human sample HG002). The Illumina reads were processed using DRAGEN, an Illumina short-read pipeline (20); in particular, we used DRAGMAP to map the reads.

For the long reads, we used four layers (4x) of PacBio Revio, with its coverage selected based on the tradeoffs between coverage and discovery performance for structural variants described in the next section. The analysis involved first mapping the long reads using minimap2 (21), then partitioning these alignments into disjoint chunks of length 151 to match the expectations of DRAGEN for short reads. The alignments preserved their original positions and CIGAR strings. These “virtual” short-read alignments were then added to the BAM constructed for the real short reads. We then used DRAGEN to call variants from this hybrid BAM.

We compared the results of the short reads alone (with DRAGEN), the 4x long reads alone (with DeepVariant), and the hybrid regime described above, against the NIST truth dataset for HG002 over their high-confidence interval regions (22). In addition to calculating whole-genome precision and recall, we also collected these over some subregions of interest as determined by NIST (23).

#### SNP Performance

Using our hybrid method, we found that SNP recall with blend-seq showed significant improvements over short reads alone (at any coverage), while maintaining precision, for combinations with both PacBio and ONT (Fig. 1a). The peak performance of blend-seq is with 25-30x shortread coverage, where it significantly outperforms short reads alone in terms of recall while matching precision, even when compared to high levels of short-read coverage where shortread performance appears to level off (Table 1 for detailed values). Even at relatively low short-read coverage (15x), we saw recall exceeding this maximal short-read recall, despite a small loss in precision in ONT combinations (Table 1).

**Table 1.** SNP performance over various genomic regions. In each region and performance statistic, blend-seq outperforms the short read controls. Blend-seq with 30x Illumina (IL) + 4x PacBio (PB) exceeds asymptotic (60x) Illumina performance for all metrics. Bold values are max values in column and shaded cells are those exceeding all control values in column (first three rows).

|  | WholeGenome |  |  | LowMappability |  |  | Exome |  |  |
| --- | --- | --- | --- | --- | --- | --- | --- | --- | --- |
|  | Recall | Precision | F1 Score | Recall | Precision | F1 Score | Recall | Precision | F1 Score |
| Illumina 15x (control) | .9889 | .9944 | .9916 | .8556 | .9312 | .8918 | .9857 | .9870 | .9864 |
| Illumina 30x (control) | .9926 | .9961 | .9944 | .8813 | .9461 | .9125 | .9887 | .9902 | .9894 |
| Illumina 60x (high depth) | .9929 | .9963 | .9946 | .8856 | .9490 | .9162 | .9895 | .9901 | .9898 |
| blend-seq 15x IL + 4x ONT | .9956 | .9927 | .9942 | .9449 | .9178 | .9311 | .9927 | .9864 | .9895 |
| blend-seq 15x IL + 4x PB | .9956 | .9951 | .9954 | .9409 | .9439 | .9424 | .9934 | .9895 | .9914 |
| blend-seq 30x IL + 4x ONT | .9965 | .9952 | .9958 | .9462 | .9382 | .9422 | .9933 | .9892 | .9912 |
| blend-seq 30x IL + 4x PB | .9963 | .9964 | .9963 | .9413 | .9521 | .9467 | .9935 | .9910 | .9922 |
| 30x PB (high depth) | **.9989** | **.9993** | **.9991** | **.9861** | **.9944** | **.9902** | **.9983** | **.9991** | **.9987** |

**Fig. 1:**
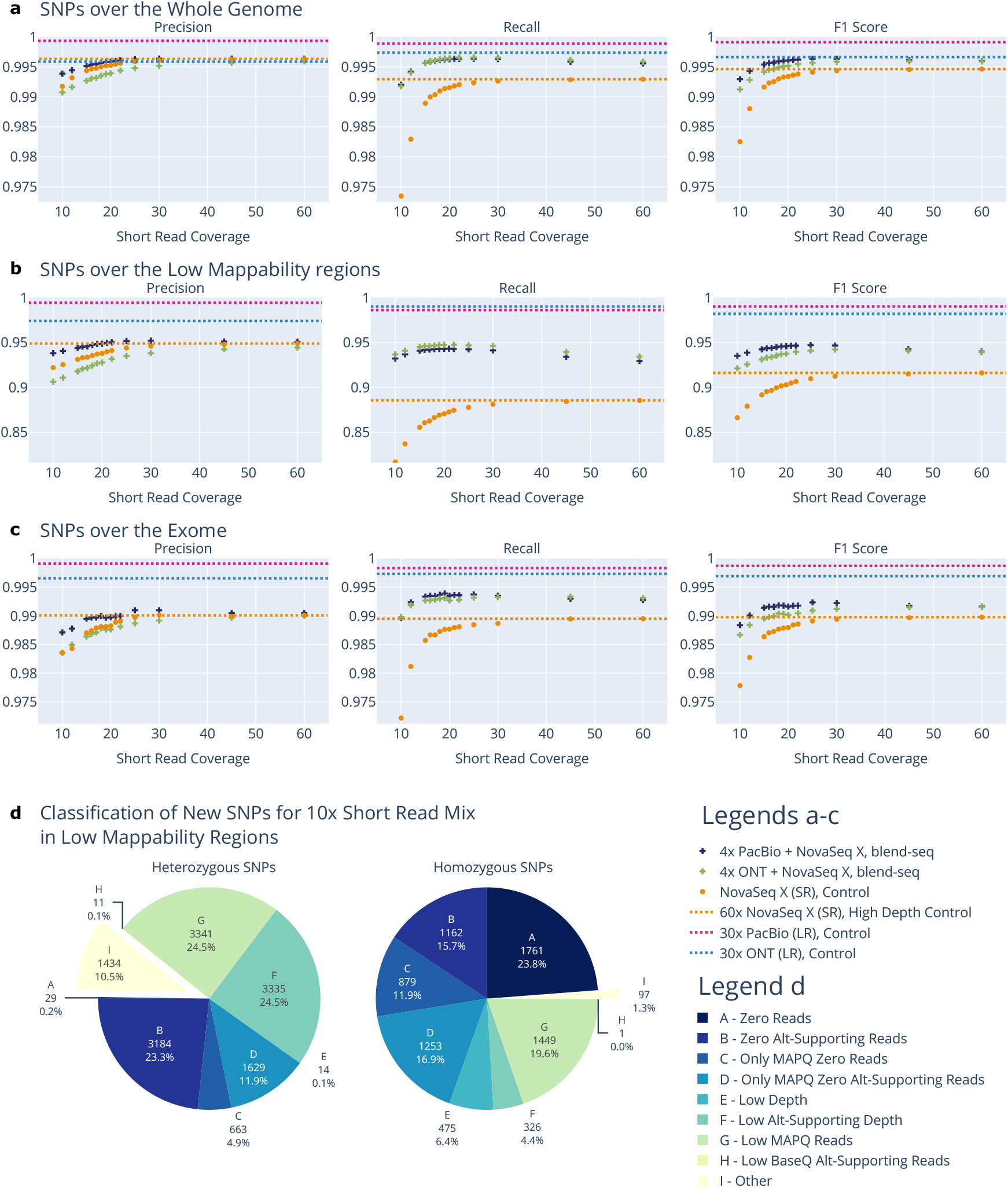
SNP performance for blend-seq in different regions. **a** The PacBio-Illumina hybrid at 4x+30x shows greater recall and precision of SNPs compared to short reads alone, even at 60x coverage; the ONT-Illumina hybrid shows greater recall but slightly lower precision. Note that even at the 4x+12x setting, both hybrids show recall exceeding short reads alone at 60x though with lower precision. **b** When restricting to low-mappability regions, the boost in hybrid performance is more pronounced. Even 4x+10x long+short read mixtures outperform the 60x short reads in recall, leading to a net F1 increase for both the PacBio and ONT hybrids with 10x short reads. **c** When restricting to the exome, blend-seq shows similar improvements in SNP performance as for the entire genome, with the PacBio-Illumina hybrid at 4x+30x outperforming recall and precision of short reads alone, even at 60x coverage. **d** A breakdown of the additional SNPs found by blend-seq in low-mappability regions. For the 30+4x Illumina-PacBio mix, we take the SNPs in low-mappability regions that were missed by short reads alone but discovered by blend-seq, then categorize them according to the nature of evidence supporting them. In the first four categories, there was no short-read evidence for the SNPs: this accounted for around 37% of the heterozygous sites and around 3/4 of the homozygous sites. The remaining labels represent sites where the short-read evidence was non-zero but limited. Detailed descriptions of each category are in the Methods section.

We saw an even stronger improvement in performance in regions that are hard to map for short reads (see Fig. 1b), substantially improving recall compared to short reads alone while making small gains in precision for the PacBio blend (see Table 1 for detailed values). These regions are difficult due to the limitations of short reads (23); as such, we expected long reads to have less difficulty mapping unambiguously. Even with only four layers of long-read coverage, we can see the complementary nature of the short and long reads boosting performance.

While improvements across the genome and in difficult regions were promising, we were concerned that these gains might only appear in regions of the genome where variants are less interpretable. To investigate whether this was the case, we ran the analysis above restricted to the exome (Fig. 1c). The result is that we see a similar gain in recall as we saw for the whole genome even in these more interpretable regions (see Table 1 for details).

#### Sources of SNP Improvement

To explain the boost in SNP performance, we characterized the new SNPs discovered using the hybrid readset. We focused on the low-mappability regions, since these experienced the largest gains. In this region, we restricted to SNPs which were previously labeled false negatives with short reads alone, but became true positives with the hybrid method. After this, we inspected a subset of sites with the Integrative Genomics Viewer (IGV) (24), and found examples where the variant caller had either no usable evidence (Supplementary Fig. 1, (**?**)), or very little (Supplementary Fig. 3). This motivated labels for the additional SNPs found in the low-mappability regions to quantify how many fell into each category. The labels, based on mapping depth and quality, articulate the different reasons why a SNP was missed in the pure short-read data but captured in the hybrid regime (details are provided in the Methods section). The breakdown of these labels for the SNPs in question is given by the pie charts in Fig. 1d for the 30x short-read and 4x long-read mix blend-seq experiment.

The first four categories (A – D) represent the case where the pure short-read variant caller has no usable evidence to make the correct call: these are easy to interpret, as the long reads were able to add valuable supporting haplotypes – enough to salvage the site and make the right call. In the 30x mix experiment, about 37% of heterozygous sites and about 78% of homozygous sites among those “corrected” in the long-reads mix fall into these categories. The remaining categories (E – I) are mostly sites where the pure shortread evidence is weak but not absent. Equivalent charts for 20x (Supplementary Fig. 5) and 10x (Supplementary Fig. 4) short-read coverage are provided as supplementary data, with similar trends observed.

#### INDEL Performance

INDELs are defined as insertion or deletion events of length less than 50bp. Applying the hybrid method above, we found that INDEL performance has greater recall for the PacBio blend compared to short reads alone, but precision is marginally lower (Supplementary Fig. 6). There was a greater boost in recall over low-mappability regions, but the overall effect was diluted by lower precision. This is most likely due to a higher INDEL error rate in long-read alignments and the fact that current tools are tuned to Illumina error rates, driving false positives for this variant type. See Table 4 for details.

### Structural Variant Discovery

In this subsection, we show how blend-seq achieves better SV performance compared to short reads, by using the long reads for discovery, and demonstrate how the short reads can be used to improve their genotypes in some situations.

#### Approach and Data

We investigated the performance of SV detection at different levels of long-read coverage, motivated by the well-established advantage of long reads for structural variant detection as compared to short reads (see e.g. (25– 32)); we do not make use of the short reads for SV discovery. We focused on structural variants (insertions and deletions with length ≥50bp) in the HG002 sample discovered using Sniffles (33) and allowing even a single read of evidence to support a call (results from other state-of-the-art SV callers, e.g. (34–36), displayed similar trends and are omitted for brevity).

For our long reads, we used PacBio Revio reads and ONT R10.4 reads from a public experiment (37), downsampled to 1–10x, as our experimental groups, and also included the full 30x setting to examine high-coverage long-read performance. These were compared against what we refer to as the “GATK-SV callset,” the HG002-specific calls from a recent Illumina 30x cohort-level callset (38). This callset is the result of performing multi-sample SV calling with GATK-SV (39) and subsequently filtering by several criteria (including restricting results to HG002). GATK-SV is a state-of-theart ensemble short-read SV discovery pipeline that integrates the output of several specialized callers, including Manta (developed by Illumina (40)), cn.MOPS and GATK-gCNV for copy-number variants (41, 42), and MELT (43) for mobile element insertions. Note that unlike our method, which operates on individual samples in isolation, GATK-SV can only be run on cohorts, as it leverages information from multiple samples to improve its precision.

Genotyping SVs is not critical for all applications, but in diagnostic scenarios, for instance, knowing the difference between a heterozygous vs. a homozygous variant could have substantial implications on a potential loss of function or other disruptions in the transcriptome. This motivated us to explore improving *genotype-aware* SV-calling performance by integrating short-read information. To do this, for each long-read technology we applied two short-read genotypers which employ a personalized (sample-specific) graph reference to try and improve the genotypes of the discovered variants. The first of these is the vg toolkit (44), which we used to create an augmented graph reference from the sample-specific SV calls produced by Sniffles as described above. Afterwards, the short reads were mapped to this graph, and genotypes were updated based on the pileup information. Note that adding SNPs to the vg graph would make graph alignment significantly slower in practice, while the short reads would still map well to easy regions, so we did not add them to the graph. The second genotyper is Paragraph (45), an SV-only tool by Illumina that maps the short reads to a linear reference, builds a local directed acyclic graph for each SV call in isolation, maps to the graph only the reads that mapped to a flanking region in the linear reference, and genotypes each SV breakpoint. In the above analysis, we found 4x long reads to be a reasonable tradeoff between coverage and SV discovery performance, and thus chose this as a fixed setting of long-read coverage for these genotyping experiments. We then varied the short-read coverage from 10 to 60x to explore the potential benefit from additional layers of short reads.

The outputs for both processes were cleaned and benchmarked with Truvari (46) against the NIST-Q100 SV V1.1 truth (9, 47), a set of calls derived from the highest-quality long-read assembly of HG002 available to date (48). Default values were used for all parameters of Truvari other than sequence comparison, which was turned off. This latter setting was used so that we could fairly compare the benchmarking results with the pure short-read pipeline, which does not report inserted bases. Note that with default settings Truvari labels variants as true positive, false positive, etc. based solely on matching alleles (up to some ambiguity in position, length, etc.), and does not consider the genotype by default. These are used to generate the usual precision, recall, and F1 score statistics reported for SV discovery. The tool also reports labels for variants which have a genotype match in addition to an allele-match, and using this criterion for defining true positives leads to our notion of “genotype-aware” versions of these statistics, denoted e.g. via “GT-Precision” instead of “Precision” in the section on SV genotyping.

#### SV Discovery Performance by Genomic Region

We observed that even at just one layer of coverage, the PacBio long reads outperformed the short reads in terms of precision and recall for the genome as a whole (Fig. 2a). With four layers of coverage, long reads substantially outperformed short reads in all regions. We observed diminishing returns on long-read depth beyond a small number of layers, with results approaching the 30x long-read performance at 10x coverage.

**Fig. 2:**
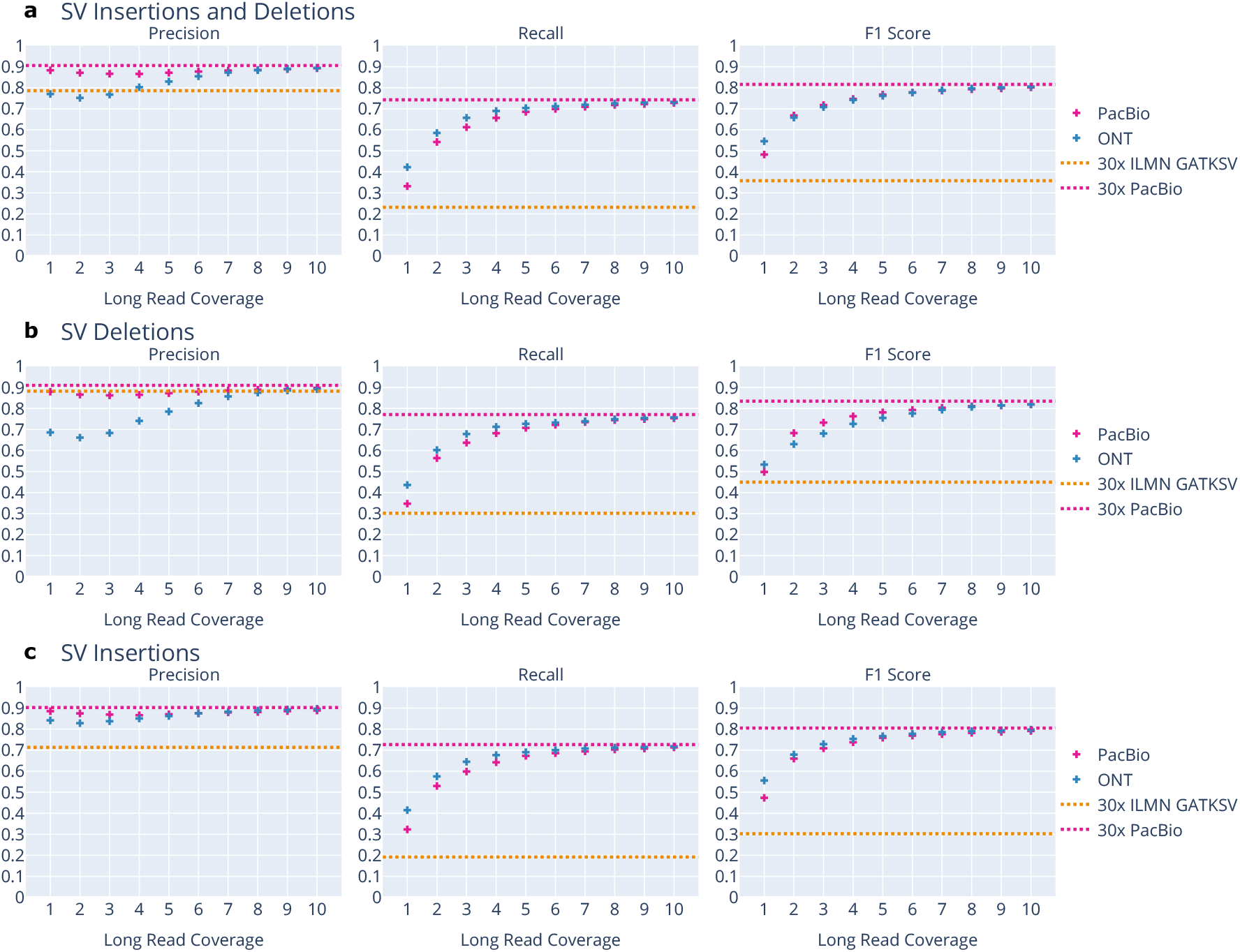
Structural variant discovery using long reads by SV type. **a** Long reads on their own, even at very low coverage, substantially outperform state-of-the-art short reads in both precision and recall. The 30x ILMN GATKSV line represents the performance of short-read SV pipelines with Novaseq X reads; PacBio-30x represents full-coverage long-read performance. **b** Performance for only SV deletions. While PacBio long reads at 4x outperform short reads substantially in terms of recall while matching precision, we note that short-read precision rivals that of long reads. This is because deletions are easier for short reads to resolve than insertions, as there is no novel sequence that cannot be mapped to the reference. **c** Performance for only SV insertions. Long reads outperform short reads even at very low coverage in this setting; this is because large insertions are difficult for short reads to resolve as they cannot be mapped to the reference. Note that these comparisons do not attempt to match the inserted sequence, as this data is not available for the short-read callset.

At 4x coverage for both PacBio and ONT, we close much of the gap to the full 30x using this pipeline; as such this serves as a compelling coverage/performance tradeoff. We broke down the results across regions of interest, such as the exome, difficult regions (NIST’s “AllDifficult” regions, deemed to be difficult for sequencing by short-read technologies (23)), and easy regions (the complement of difficult regions). Table 2 shows details for all of these regions for short reads alone, 1x and 4x coverage for both PacBio and ONT, and 30x PacBio (a proxy for the asymptotic limit of this technology). We see similar trends in all these regions, with 4x long reads substantially outperforming the short reads even in the easier regions where we expect short reads to perform the best.

**Table 2.** SV discovery by region. Long reads (both PacBio and ONT) outperform short reads in in recall and precision, even at 1x coverage when evaluated over the whole genome (except ONT precision), and at 4x for all individual regions (except ONT precision in the Easy Regions). Bold values are max values in column and shaded cells are those exceeding the control value in column (first row).

|  | Whole Genome |  |  | Easy Regions |  |  | Hard Regions |  |  | Exome |  |  |
| --- | --- | --- | --- | --- | --- | --- | --- | --- | --- | --- | --- | --- |
|  | Recall | Precision | F1 Score | Recall | Precision | F1 Score | Recall | Precision | F1 Score | Recall | Precision | F1 Score |
| Illumina 30x (control) | .2316 | .7860 | .3578 | .7205 | .8633 | .7855 | .1749 | .7611 | .2844 | .2168 | .5349 | .3085 |
| ONT 1x | .4225 | .7702 | .5457 | .5319 | .7255 | .6138 | .4087 | .8240 | .5464 | .4126 | .7740 | .5382 |
| PacBio 1x | .3318 | .8830 | .4824 | .4218 | .9332 | .5810 | .3201 | .8789 | .4693 | .2517 | .8395 | .3873 |
| ONT 4x | .6897 | .8026 | .7419 | .8787 | .7808 | .8268 | .6639 | .8381 | .7409 | **.7692** | .7714 | **.7703** |
| PacBio 4x | .6566 | .8658 | .7468 | .8332 | .9211 | .8750 | .6330 | .8612 | .7296 | .5315 | .8596 | .6568 |
| PacBio 30x (high depth) | **.7426** | **.9055** | **.8160** | **.9553** | **.9464** | **.9508** | **.7140** | **.8983** | **.7956** | .6643 | **.8819** | .7578 |

Additional figures in the style of Fig. 2a for these regions are in the supplement (exome: Supplementary Fig. 11; easy regions: Supplementary Fig. 12, Supplementary Fig. 15; table: Table 5).

#### Discovery Performance by SV Type

When we break down our results by SV type, we see a difference in performance improvement for deletions vs. insertions (Fig. 2b, Fig. 2c). Blend-seq results in large gains in recall for both categories, but for deletions there is a drop in precision for the ONT reads, and short reads already achieve relatively high precision in this category. This is not surprising, given that short reads can gather accurate depth evidence for large deletions, but struggle to assemble large inserted sequences.

We broke down our results by SV length, to see whether performance varies between relatively short (100-250bp) events and larger (2.5k-10k) events. We expected to see a difference between PacBio and ONT performance: while ONT reads are longer (mean 50.7kb with standard deviation 66.9kb), PacBio reads (mean 15.6kb with standard deviation 3.9kb) have higher per-base accuracy, so we expected to see ONT do better in larger events and PacBio in shorter events. This is indeed the case, as shown in Fig. 3a, Fig. 3b, and Table 3. The PacBio reads yield better precision for smaller events even at extremely low coverage, and the ONT recall outperforms the PacBio reads for larger events, surpassing the PacBio 30x recall even with just 3x of ONT reads. In both cases, long reads outperform the short read control.

**Table 3.** SV discovery by event size. While long reads outperformed short reads in most categories, the two long read technologies showed different strengths. For shorter events, we found PacBio performed best, likely due to its higher per-base accuracy; on larger events, ONT had better recall, leveraging its longer read lengths. This was true even when 4x ONT was compared against 30x PacBio reads. Bold values are max values in column and shaded cells are those exceeding the control value in column (first row).

|  | 100-250 |  |  | 2.5k-10.0k |  |  |
| --- | --- | --- | --- | --- | --- | --- |
|  | Recall | Precision | F1 Score | Recall | Precision | F1 Score |
| Illumina 30x (control) | .1733 | .8120 | .2857 | .2371 | .7493 | .3602 |
| ONT 1x | .3987 | .7640 | .5240 | .5278 | .8745 | .6583 |
| PacBio 1x | .3193 | .8854 | .4694 | .2418 | .8735 | .3787 |
| ONT 4x | .6498 | .7992 | .7168 | **.8805** | .8304 | .8547 |
| PacBio 4x | .6317 | .8681 | .7313 | .6322 | **.8864** | .7380 |
| PacBio 30x (high depth) | **.6997** | **.9101** | **.7912** | .8307 | .8848 | **.8569** |

**Table 4.**
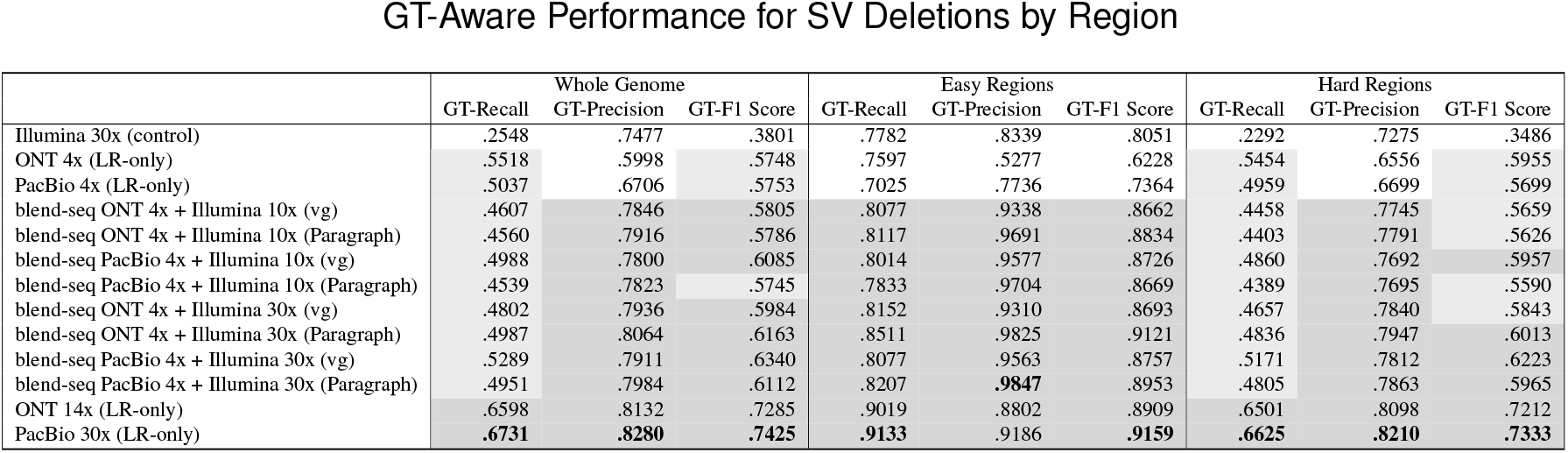
GT-Aware performance for SV deletions after using short-read genotypers. The blend-seq short-read genotyped SVs have better GT-Precision over the whole genome compared to the short-read 30x control and the 4x long-read controls, with recall still exceeding the 30x short reads alone. Over the easy regions, the blend-seq hybrids beat out these controls in all GT-aware metrics. Even over hard regions, we see the GT-Precision for the blend-seq mixtures doing better than these controls, which causes GT-F1 Scores to outperform these controls when using 30x short reads in addition to the 4x long reads. Bold values are max values in column, lightly shaded cells are those exceeding the short read control (first row), and deeply shaded cells are those where the hybrid method exceeds both short reads and long reads on their own (first three rows).

|  | Whole Genome |  |  | Easy Regions |  |  | Hard Regions |  |  |
| --- | --- | --- | --- | --- | --- | --- | --- | --- | --- |
|  | GT-Recall | GT-Precision | GT-F1 Score | GT-Recall | GT-Precision | GT-F1 Score | GT-Recall | GT-Precision | GT-F1 Score |
| Illumina 30x (control) | .2548 | .7477 | .3801 | .7782 | .8339 | .8051 | .2292 | .7275 | .3486 |
| ONT 4x (LR-only) | .5518 | .5998 | .5748 | .7597 | .5277 | .6228 | .5454 | .6556 | .5955 |
| PacBio 4x (LR-only) | .5037 | .6706 | .5753 | .7025 | .7736 | .7364 | .4959 | .6699 | .5699 |
| blend-seq ONT 4x + Illumina 10x (vg) | .4607 | .7846 | .5805 | .8077 | .9338 | .8662 | .4458 | .7745 | .5659 |
| blend-seq ONT 4x + Illumina 10x (Paragraph) | .4560 | .7916 | .5786 | .8117 | .9691 | .8834 | .4403 | .7791 | .5626 |
| blend-seq PacBio 4x + Illumina 10x (vg) | .4988 | .7800 | .6085 | .8014 | .9577 | .8726 | .4860 | .7692 | .5957 |
| blend-seq PacBio 4x + Illumina 10x (Paragraph) | .4539 | .7823 | .5745 | .7833 | .9704 | .8669 | .4389 | .7695 | .5590 |
| blend-seq ONT 4x + Illumina 30x (vg) | .4802 | .7936 | .5984 | .8152 | .9310 | .8693 | .4657 | .7840 | .5843 |
| blend-seq ONT 4x + Illumina 30x (Paragraph) | .4987 | .8064 | .6163 | .8511 | .9825 | .9121 | .4836 | .7947 | .6013 |
| blend-seq PacBio 4x + Illumina 30x (vg) | .5289 | .7911 | .6340 | .8077 | .9563 | .8757 | .5171 | .7812 | .6223 |
| blend-seq PacBio 4x + Illumina 30x (Paragraph) | .4951 | .7984 | .6112 | .8207 | **.9847** | .8953 | .4805 | .7863 | .5965 |
| ONT 14x (LR-only) | .6598 | .8132 | .7285 | .9019 | .8802 | .8909 | .6501 | .8098 | .7212 |
| PacBio 30x (LR-only) | **.6731** | **.8280** | **.7425** | **.9133** | .9186 | **.9159** | **.6625** | **.8210** | **.7333** |

**Table 5.** GT-Aware performance for SV insertions after using short-read genotypers. Using the short-reads to genotype the SV insertions from the long reads, the blend-seq experiments perform well over the whole genome in GT-Precision, beating out the short-read control at 30x and the 4x long-read controls. In easy regions, the GT-F1 Scores for the blend-seq experiments outperform these controls as well, due to a mix of improvements in both GT-Precision and GT-Recall. Even in hard regions, where short reads struggle the most to accurately map, the 4x ONT and 30x Illumina blend-seq hybrid has the best GT-Precision out of all groups, including the high-depth long reads samples. Bold values are max values in column, lightly shaded cells are those exceeding the short read control (first row), and deeply shaded cells are those where the hybrid method exceeds both short reads and long reads on their own (first three rows).

|  | Whole Genome |  |  | Easy Regions |  |  | Hard Regions |  |  |
| --- | --- | --- | --- | --- | --- | --- | --- | --- | --- |
|  | GT-Recall | GT-Precision | GT-F1 Score | GT-Recall | GT-Precision | GT-F1 Score | GT-Recall | GT-Precision | GT-F1 Score |
| Illumina 30x (control) | .1435 | .5633 | .2287 | .4758 | .6736 | .5577 | .0769 | .5012 | .1334 |
| ONT 4x (LR-only) | .3770 | .5644 | .4521 | .5387 | .5461 | .5424 | .3447 | .5577 | .4261 |
| PacBio 4x (LR-only) | .3419 | .5491 | .4214 | .4955 | .5535 | .5229 | .3109 | .5303 | .3920 |
| blend-seq ONT 4x + Illumina 10x (vg) | .2535 | .6661 | .3672 | .5223 | .7104 | .6020 | .1997 | .6473 | .3053 |
| blend-seq ONT 4x + Illumina 10x (Paragraph) | .2919 | .6145 | .3958 | .5490 | .6483 | .5945 | .2405 | .5862 | .3411 |
| blend-seq PacBio 4x + Illumina 10x (vg) | .2627 | .6589 | .3756 | .5249 | .7065 | .6023 | .2105 | .6373 | .3165 |
| blend-seq PacBio 4x + Illumina 10x (Paragraph) | .3011 | .5980 | .4005 | .5394 | .6372 | .5842 | .2534 | .5665 | .3502 |
| blend-seq ONT 4x + Illumina 30x (vg) | .2903 | **.6792** | .4067 | .5582 | **.7193** | .6286 | .2368 | **.6646** | .3492 |
| blend-seq ONT 4x + Illumina 30x (Paragraph) | .3201 | .6481 | .4285 | .5720 | .6759 | .6197 | .2699 | .6225 | .3766 |
| blend-seq PacBio 4x + Illumina 30x (vg) | .3024 | .6706 | .4169 | .5470 | .7135 | .6192 | .2536 | .6523 | .3652 |
| blend-seq PacBio 4x + Illumina 30x (Paragraph) | .3318 | .6323 | .4352 | .5612 | .6694 | .6105 | .2861 | .6030 | .3881 |
| ONT 14x (LR-only) | .4418 | .6595 | .5292 | .6535 | .6427 | .6481 | .3994 | .6370 | .4910 |
| PacBio 30x (LR-only) | **.4601** | .6741 | **.5469** | **.6719** | .6662 | **.6691** | **.4176** | .6525 | **.5093** |

**Fig. 3:**
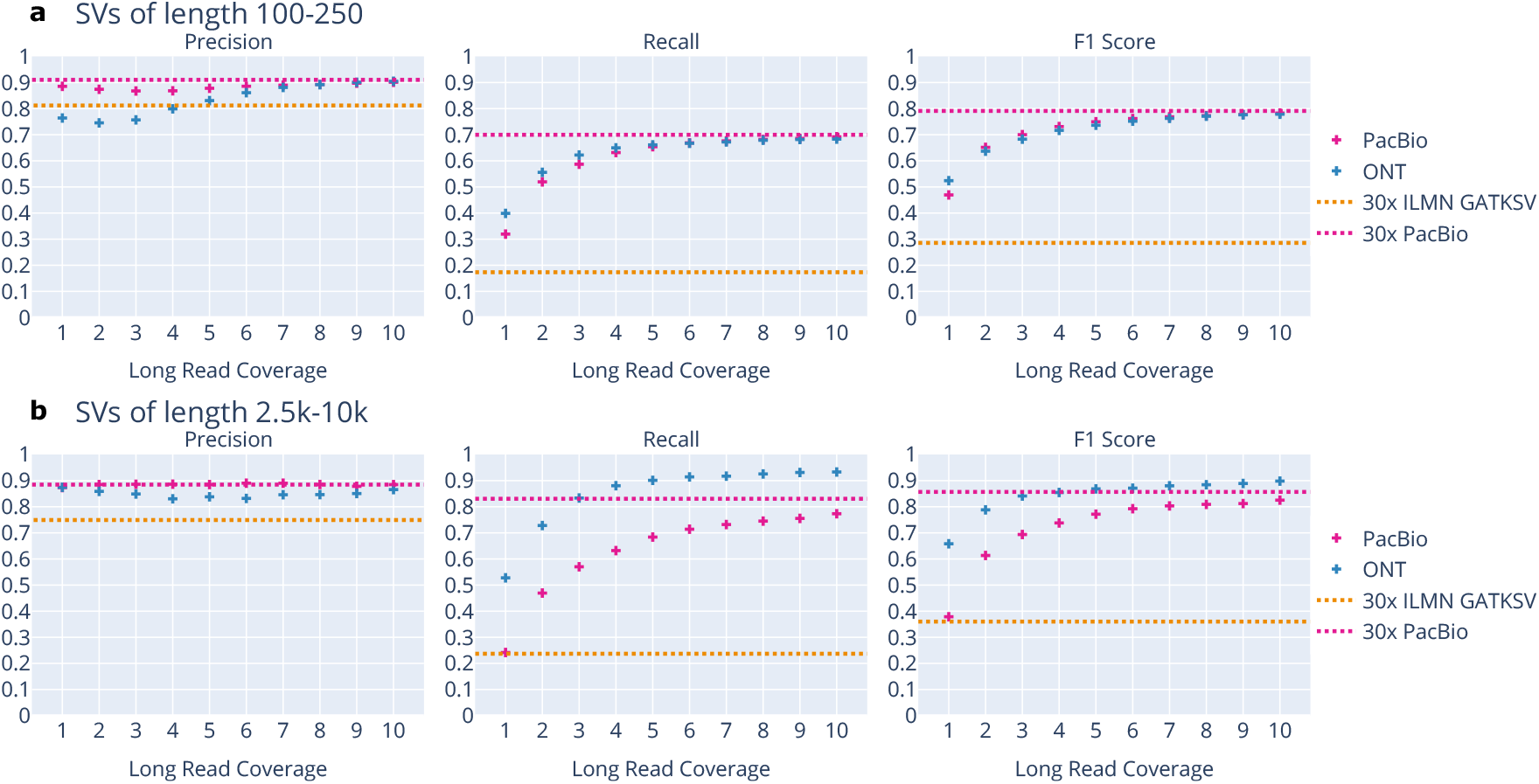
Structural variant discovery using long reads by size of event. **a** Performance on 100-250 bp structural variants for short and long reads. These are the most common type of SVs; as such, performance trends largely follow the earlier results for all SV. **b** Performance on 2.5k-10k bp structural variants for short and long reads. When restricting to these larger events, we see the ONT Ultra Long reads have better recall than PacBio.

#### Genotyping SVs with Short Reads

For the genotype-aware setting, we initially did not see any improvements overall in using the short-read genotypers on the SV calls (Supplementary Fig. 29), even when restricting to GT-aware statistics (Supplementary Fig. 30). However, we investigated performance metrics across different regions, especially where the short reads may have an advantage over the long reads with their extra depth without facing mapping difficulty. The boost in genotype-aware performance provided by the short reads is greatest where the short reads are best able to map: as in the last subsection, we refer to these as “Easy Regions” in our plots, comprised of the complement of NIST’s “AllDifficult” regions. This setting reduces the total 30k variants in our truth dataset to 11.5k overlapping or contained within this region (23). We see that the GT-aware statistics for blend-seq outperform the short-read and 4x long-read controls overall (Fig. 4a).

**Fig. 4:**
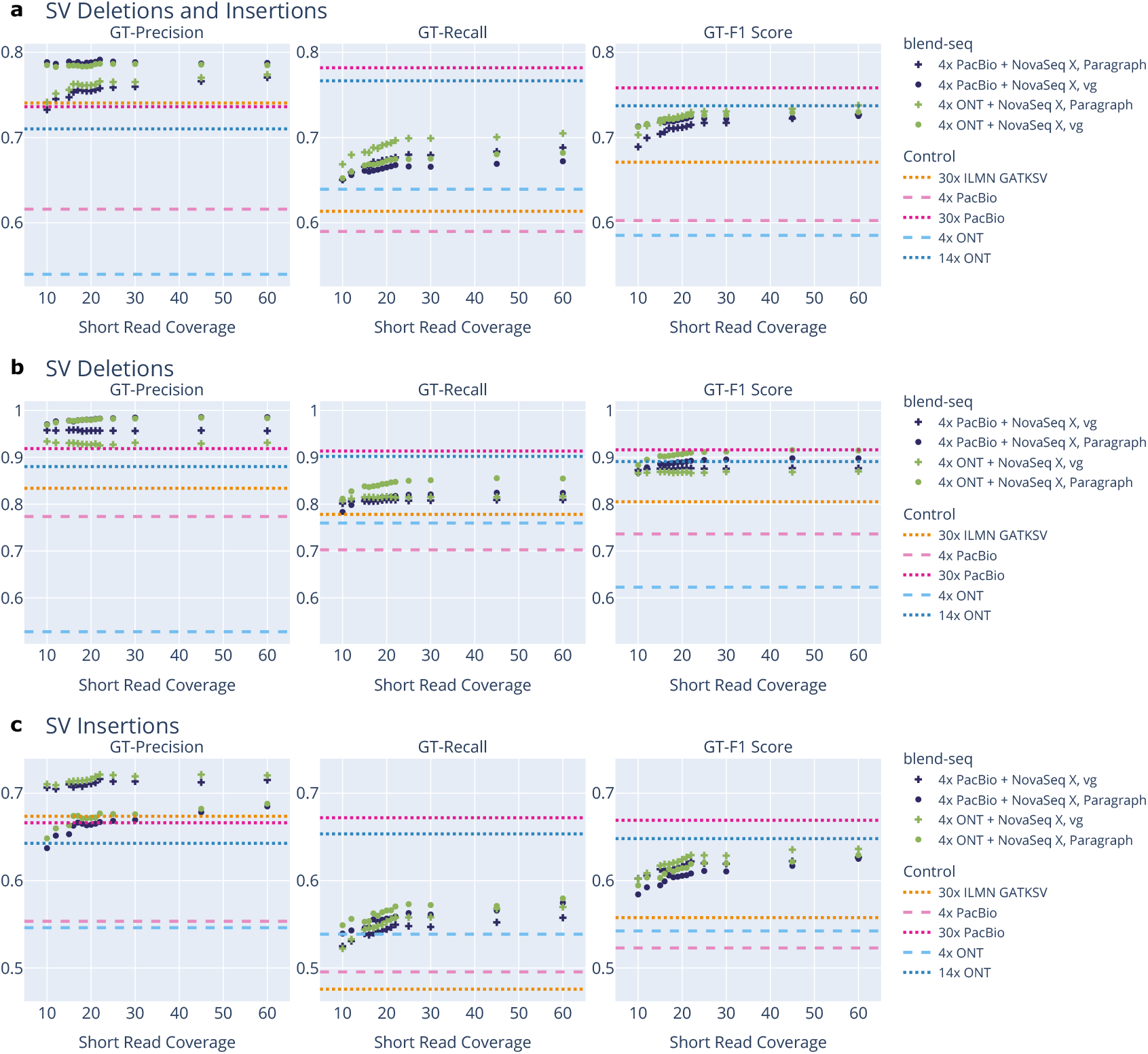
GT-Aware structural variant discovery blend-seq: performance in easier regions by variant type. **a** We see the GT-aware statistics are an improvement over the long-read and short-read controls, with GT-Precision exceeding even the high-depth long-reads’ performances. This results in the blend-seq experiments having better GT-F1 Scores compared to all controls powered also by an increase in GT-Recall. **b** On SV deletions, we see that Paragraph acheives near perfect GT-Precision, while also increasing GT-Recall. This puts the GT-F1 Score on par even with the 30x PacBio performance. **c** On SV insertions, there is a bit of precision-recall trade-off happening between the genotypers, where Paragraph improves GT-Precision further at a greater cost to GT-Recall compared to vg. In both cases, the GT-F1 Scores exceed the short-read and long-read controls.

Furthermore, this trend holds even when breaking down by SV deletions and SV insertions. In the SV deletion case (Fig. 4b), the blend-seq mixtures achieve near perfect GT-precision when using Paragraph, beating even the high-depth long-read controls, and resulting in a GT-F1 Score that is almost on par with 30x PacBio. For SV insertions (Fig. 4c), the short reads help both with GT-Precision and GT-Recall, outperforming all short-read and lower-depth long-read controls.

We remark that even outside the GT-aware setting, the short reads are useful in filtering out false positive SV calls made by the low-depth long reads, as evidenced in Supplementary Fig. 31 for deletions and Supplementary Fig. 32 for insertions, where precision increases even above the 30x PacBio control. For SV deletions, this comes with similar recall when compared to the low-depth long-read controls, making this method a net boost to SV deletion performance in easy regions. For SV insertions, the increased precision comes with lower recall, making this method only more useful where precision is more important than recall, such as some diagnostic settings.

We also investigated using long-read genotypers to improve the base set of calls, but found these to have no significant difference comapred to Sniffles (see Supplementary Fig. 21 and Supplementary Fig. 22).

### Phasing

In this subsection, we use blend-seq for shortvariant discovery via the methods previously described, and then phase the variants using the long reads, greatly outperforming the short-read controls based on standard phasing performance metrics.

#### Approach and Data

A set of heterozygous variants is said to be *phased* if for each pair in the set it can be determined whether they appear on the same copy of a chromosome or not. These are organized into *phase blocks*, i.e. subsets of variants in the VCF that are phased, whose lengths are computed as the distance from the first to last variant in the block. Larger phase blocks provide information about relative positioning of an individual’s variants, hence one desires larger phase blocks without accumulating phasing errors.

To test our ability to produce accurate phase blocks, we produced variants with the methods previously described. For pure short-read control groups, we took the set of small variants in the short-read VCF and used the short reads to phase them using WhatsHap (49), a phasing tool designed for short reads. This produced longer, better phase blocks than the default phase blocks produced by DRAGEN (Supplementary Fig. 36). Note we did not use the SVs called by GATK-SV for short-read phasing, since that output does not provide insertion sequences as required by the phasing tools. For the long-read control groups, we used the full set of SNPs, INDELs, and SVs called by the long-read pipeline. We then used the long-read phaser LongPhase (50), with comparisons to HiPhase (51) provided in the supplement (Supplementary Fig. 37 and Supplementary Fig. 38). For the blend-seq groups, we phased using the long-read data alone.

For the blend-seq scenarios, we began with the SNP and INDEL VCFs produced following the procedure described in the first section. Structural variants were called using the process of the second section, using the low-coverage parameters of Sniffles on the long reads alone, but without applying any further regenotyping. Although LongPhase allows for multiple BAMs to use for phasing, allowing us to use both short- and long-read BAMs, we found performance to be worse than using the long reads alone (Supplementary Fig. 39 and Supplementary Fig. 40). As such, all variants (SNP, INDEL, SV) from the hybrid process were combined together into one VCF and then phased together using the long reads alone. Note that while SVs were included in the phasing process, performance was only evaluated with respect to SNPs and INDELs to allow for more meaningful comparison with the short-read pipeline, where SVs are not available. The effect of including these is measured in Supplementary Fig. 41.

#### Phasing Performance

The standard phasing performance metrics can be thought of as falling into two groups: measures of how extensively the genome can be phased by a given algorithm without regard to accuracy (which does not require a truth baseline), and measures of the accuracy of the resulting phasing (relative to a phased truth baseline). Beginning with the former, we looked at metrics borrowed from genome assembly. The NG50 statistic, computed by WhatsHap, measures the minimum phase block length such that at least 50% of the genome belongs to a phase block of that length or longer. A larger number thus means that longer distance haplotype information is available for most of the genome (more than 50%). For blend-seq, we measured this statistic across a wide range of short-read coverages (10-60x) paired with either PacBio or ONT long reads at a fixed 4x coverage.

For the short-read controls, NG50 is zero because less than half of the bases are covered by phase blocks (see Fig. 5a). The NG50 for blend-seq using 4x PacBio hovers just under 100k (50% of reference bases are in phase blocks 100kb or greater in length), and the 4x ONT blend-seq NG50 is an order of magnitude greater at around 2-3M. This is due to the longer reads that allow us to better join phase blocks into larger pieces. It is also interesting to note that blend-seq does better than the 4x long-read controls alone: part of this can be explained by additional variants produced using the blend-seq short variant calling procedures from the first section, which may help bridge otherwise distant heterozygous variants. For example, the 4x PacBio control has NG50 equal to zero as well, failing to phase over half of the genome.

**Fig. 5:**
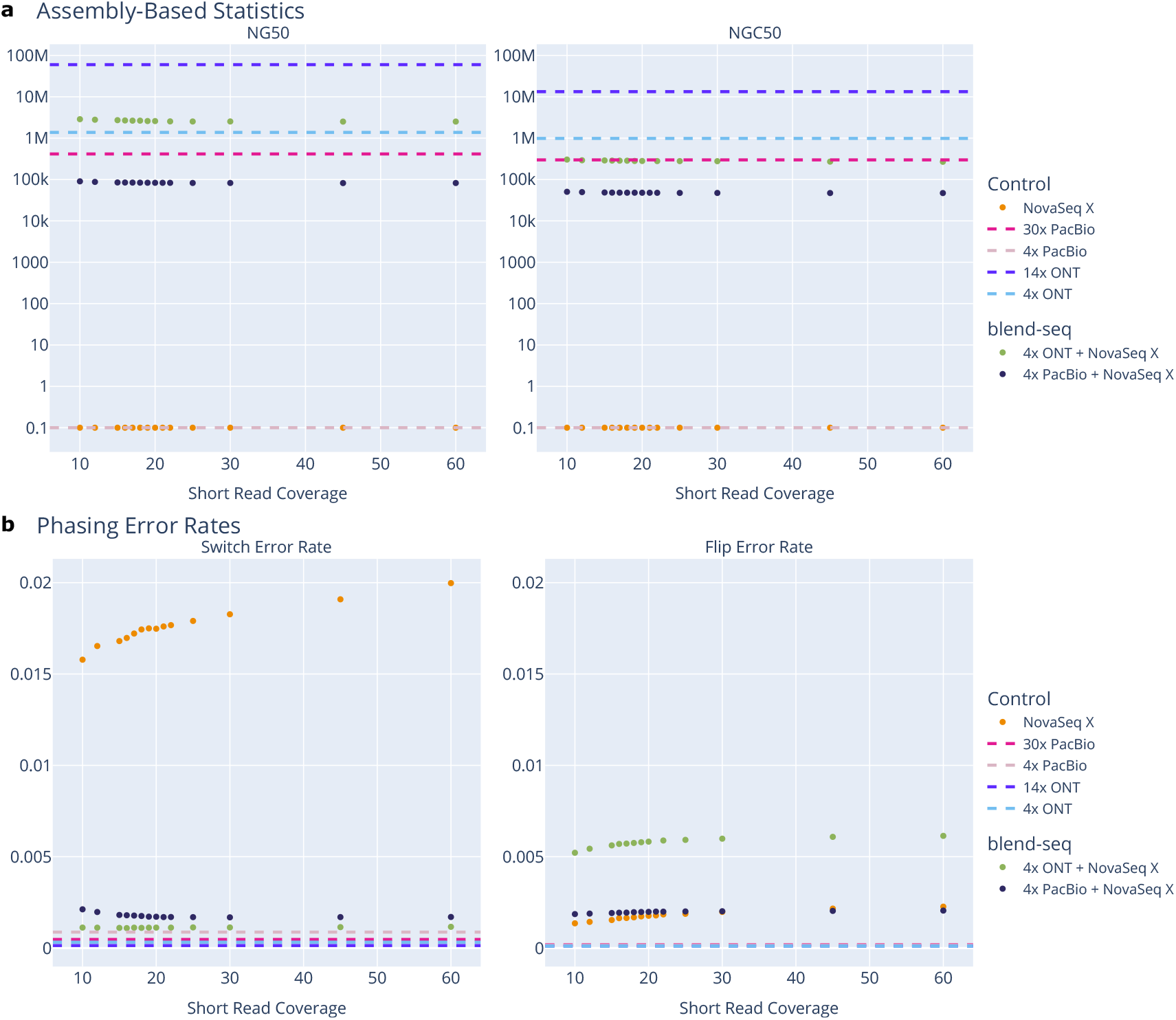
Phasing performance for blend-seq. **a** NG50 metric (50% of the genome is within a phase block of this length) and NGC50 metric (NG50 but accounting for phasing truth data) for different settings of blend-seq and pure short and long read controls. Because the y-axis is log-scaled, NG50 values of zero (for NovaSeq X and the 4x PacBio conditions) are rescaled to 0.1. We see three distinct clusters of performance correlating with average read length. Short reads struggle to bridge distant variants and achieve an NG50 of zero, and the same is true for 4x PacBio due to missing variants at low coverage. Combining these two, blend-seq PacBio mixtures have an NG50 starting around 100k, beating out both of these controls. In the third cluster, blend-seq with ultralong ONT reads have approximately 20 times larger NG50 than the corresponding PacBio groups. For NG50, the blend-seq mixtures provide better phasing than their 4x long-read controls, likely due to better variant discovery with the additional short-read coverage. When accounting for phasing truth data in the NGC50 metric, this blend-seq improvement persists for the PacBio data, but with ONT the performance of long reads alone exceeds that of blend-seq. Note that this comparison is confounded by the fewer number of variants the 4x ONT can call (see the first section on SNP calling), hence giving the long-read control fewer opportunities to make a phasing error and thus potentially inflating the NGC50 score. **b** Switch error rates and flip error rates for different settings of blend-seq and pure short and long read controls. In the blend-seq regimes, the switch error rate decreases with extra layers of short reads, and is significantly smaller than the short-read controls. The PacBio blend-seq switch error rate is nearly as low as the corresponding 4x long-read control. The flip error rate for blend-seq conditions stays mostly constant across short-read coverages for PacBio combinations, and increases slightly with coverage for ONT combinations. The flip error rates for PacBio blend-seq are very similar to the short-read controls; both of these curves sit slightly above the long-read controls.

In diploid organisms, when comparing with a truth baseline, there are two possible matches between the true pair of haplotypes and the inferred pair of haplotypes, and tools like WhatsHap choose the best match. One can expand the notion of a correct match by allowing a switch at some point within the block: these are called switch errors, and two consecutive switch errors are labeled as one flip error. While NG50 does not account for errors within phase blocks, we compute the NGC50 (or “NG50 corrected”) as computed by vcfdist (52), which first splits phase blocks at switch errors (based on the truth data) and then computes NG50 on this refined partition; the truth data used here is the fully-phased NIST-Q100 dataset from the second section. These results are shown in Fig. 5a for the different experimental groups. We see the blend-seq mixtures with PacBio maintain their superior phasing capabilities over the short reads when accounting for phasing errors in this way, but the ONT hybrids’ performance drops below their corresponding long-read control. This is potentially due to the blend-seq conditions calling substantially more variants but at lower precision, opening up the opportunity for a greater number of switch errors.

The frequency of switch errors is characterized by the *switch error rate*, defined as the number of switch errors divided by the number of pairs where a switch can occur (53). As opposed to NGC50, this metric is normalized by the number of called variants, so calling a greater number of variants does not inherently risk reducing the score; however, it still does not account for the relative difficulty of those variants, so it is possible to call/phase only “easy” variants and not be penalized for missing variants that are harder to call/genotype. This error rate for the blend-seq mixtures was significantly lower (better) than the short-read control, but it was slightly higher (worse) than the 4x long-read controls (Fig. 5b). For flip errors, the rate was about 3x higher for the ONT blend-seq experiments than the short-read control.

## Discussion

We have demonstrated how blend-seq, by combining or selecting from standard-coverage short reads and low-coverage long reads, can provide substantial gains in SNP precision and recall, in SV discovery and genotyping, and in phasing variants. However, we recognize that we have demonstrated these results with specific choices of datasets, pipelines, and settings. In this section we discuss a few limitations of these choices, with a view towards further developments.

In terms of genomic datasets, we presented all results on a single genome, HG002, because it is the genome with the highest-quality truth data available. In future work, a proxy for ground truth may be derived from high-quality assemblies like those in the Human Pangenome Reference Consortium (54) or the Human Genome Structural Variation Consortium (55), combined with assembly-based variant callers like Dipcall (56) or hapdiff (57).

Further, the tools used here represent just one set of choices in a vast and evolving landscape: for example, several longread mappers exist, as well as several SV callers and genotypers, both for long and for short reads, each with parameters that could be tuned for particular applications (see e.g. (58–65)); the field is under constant development, particularly for long reads.

For short variants, we emphasized the performance of SNPs but not INDELs due to reduced precision in the latter, likely due to greater false positives in the long-read alignments. Despite this, we observed a similar boost in INDEL recall, and hypothesize that with additional tuning of available filtering models (either for the alignments or for the variants) we might retain the boost in recall without sacrificing precision, similar to the SNPs.

Also regarding short variants, the DRAGEN version used in this paper (v3.7.8) was chosen to match the version used by the All of Us Research Program (66), a large biobank funded by the U.S. government. It should be noted that recent work shows improved performance of DRAGEN in later versions (67). However, no versions past v3.7.8 have an open-source implementation, while the variant caller we used should be functionally equivalent to DRAGEN-GATK (68), which is open-source.

Prior work on combining long and short reads for variant calling has mostly focused on SNPs and (short) INDELs. Specifically, Clair3-MP (69) combined Illumina and either PacBio Sequel II reads or ONT R9 reads (both superseded by newer versions), but only went down to 10x long reads, which did not outperform the short reads alone. DeepVariant (70) includes a hybrid model, but it is intended for only 30x Illumina plus 30x PacBio reads, which was beyond the long-read depth we considered. Simultaneous to our work, a hybrid model has been retrained (71) for the Illumina and ONT regime, showing some improvements compared to pure Illumina and pure ONT in the 20x Illumina plus 30x ONT setting. The authors also observed that their model starts to saturate in the 15x Illumina plus 15x ONT regime, at a depth of long-reads beyond our consideration.

Prior work exists on SV discovery performance at various levels of long-read depth. First, long-read downsampling analyses for SV discovery have been performed in the past, but those studies either did not consider coverage levels as low as ours (72, 73), focused on just a few known SVs challenging for short reads (73), used more stringent parameters in the callers (72, 73), or compared to less accurate and complete truthsets (72). Second, although it is in principle possible to model the theoretical recall of structural variants as a function of coverage and read length (see e.g. (74)), our perspective is that practical issues like the association of SVs with repeats, the effect of sequencing errors, and of read mapping algorithm on SV calling require the systematic empirical investigation we present here. Third, a simulation framework for estimating the combination of Sanger, 454, and Illumina reads that best reconstructs any specific predetermined SV was presented in (75), but is not modular to extend to the technologies we investigated.

In addition, because we did not want phasing performance to confound SV discovery performance, we decided not use tools like Truvari refine (76) or vcfdist (52), which require phased variants, to better compare the resulting haplotypes to the truth. This resulted in our recall being lower than what was reported by (62), which did use phasing information and thus performed this processing step. To ensure this was the source of the discrepancy, we phased our 30x PacBio sample using HiPhase (51) and ran Truvari refine, which yielded comparable results (Supplementary Fig. 20) to (62).

We note that personalized graph references for short-read mapping, like those we used for genotyping SVs, are under active development in the bioinformatics literature, but they are typically built by subsetting a large-cohort pangenome to the parts that are most similar to the short reads for the sample at hand (77–81). As such they are less likely to contain rare SVs carried by the sample compared to our approach.

For phasing, in theory only one accurate read connecting variants could suffice to phase them, so one would expect performance to increase with further layers of long reads. This is explored in the supplement using long reads alone (Supplementary Fig. 42 and Supplementary Fig. 43). Hybrid phasing methods typically combine long reads with other longrange technologies, like Hi-C, Strand-seq, or linked reads, or perform trio binning with short reads from the parents. All such approaches would increase the cost significantly, so we did not consider them. Some existing methods do combine long and short reads from the same sample, but they were not tested or designed for diploid applications (82, 83).

Related to these, much work has gone into hybrid assembly techniques, but we did not pursue these as we were interested in cost-effective solutions that minimize the amount of long reads.

The sequencing technologies investigated in this paper represent some of the most popular options, but are only a subsample of those that are now commercially available. Blend-seq provides a framework for combining various technologies in a rapidly shifting landscape. Although panels of common short and structural variants are currently being built for use in imputation pipelines, blend-seq will remain applicable to rare variant discovery and the single-sample diagnostic setting. As such, we expect the blend-seq approach will remain relevant even as current and new sequencing technologies continue to evolve.

## Methods

### Short Variant Calling

Blend-seq short variant calls were generated by combining preprocessed long reads with the Illumina short reads to create a hybrid Binary Alignment Map (BAM). This BAM was used as input to DRAGEN v3.7.8, with the version selected to match what is used for the All of Us Research Program (66). (Results for v4.2.7 are also included in the supplement Supplementary Fig. 7–10.) The tool was designed for Illumina short reads alone. Therefore, several preprocessing steps on the long reads were necessary to enable DRAGEN to run. The long read BAMs were first downsampled to 4x coverage with SAMtools (84) to simulate a lower coverage long read product. Then the following processing steps were performed:

1. In order to remove large-scale deletion information which cannot be processed by DRAGEN, which is designed for short INDELs alone, we started by converting large deletions in the long read alignments to use “N” in their CIGAR strings rather than “D” via a custom pysam script.
2. GATK’s `SplitNCigarReads` (85) was used to split alignments at these large deletions to provide cleanly aligned reads as a technical intermediate to feed into the following custom tool.
3. To simulate a short read structure from the long reads, we wrote a custom `chop-reads` program to chop the alignments evenly at 151 read bases and update the CIGAR strings accordingly. This allows DRAGEN to see reads of equal length regardless of which technology generated the read, while still keeping the alignment information generated from the original long reads.
4. To generate the final hybrid BAM, SAMtools was used to sort (84) and then merge these reads with the DRAGMAP aligned short reads. The header was also updated to just have one read group via SAMtools reheader, to work around a limitation in the DRAGEN version used.
5. Chopped long reads with CIGAR string equal to a pure insertion (i.e. “151I”) were removed, since these are not useful for short INDEL calling and caused errors in the DRAGEN pipeline.
6. SAMtools was used to index these hybrid BAMs and then they were run through the full DRAGEN variant calling tool.

For comparison with the hybrid samples, control samples for just long and just short reads were processed using bestpractice SNP and short INDEL calling pipelines for long and short reads respectively. The long reads were mapped using minimap2 for ONT (86, 87) and the wrapper pbmm2 written by PacBio for the PacBio reads. The short reads were mapped using DRAGMAP as a part of the DRAGEN (88) variant calling process, using the GRCh38 human reference in both cases. The long reads (PacBio and ONT) were run through DeepVariant (70) (using v1.3 for PacBio and v1.6 for ONT), while the short reads (Illumina) were run through DRAGEN v3.7.8.

After obtaining the hybrid DRAGEN Variant Call Format files (VCFs) at each desired mixture of coverages, we benchmarked against the NIST HG002 v4.2.1 standard Genomes in a Bottle (GIAB) truth set (8, 89). We used a pipeline wrapping RTG’s `vcfeval` tool (90, 91) which resolves haplotypes underlying variant representations to ensure accurate comparisons. From this, we obtain true positive (TP), false positive (FP), and false negative (FN) labels from which precision, recall, and F1 statistics are derived, including over different subsets of the genome. These BED file regions (“Low Mappability”, and others used in this paper) were also curated by NIST for their GIAB release (92).

To create the pie charts demonstrating the breakdown of types of short read evidence (or lacking) among the variants uniquely picked up by the hybrid method, we took that set of SNPs labeled with FN from `vcfeval` comparing the pure short read VCF against the NIST truth, and intersected with those labeled TP from `vcfeval` comparing the hybrid VCF against the NIST truth. Then we restricted to those contained in NIST’s low mappability regions, where the largest boost in performance was observed. Each SNP was given a unique label (using the `AnnotateNewSNPs.wdl` workflow) in the following list with higher labels taking precedence, depending on how the evidence in the pure short read BAM looked:

A. Zero Reads: zero depth (DP) at the site;
B. Zero Alt-Supporting Reads: zero short reads supporting the SNP in their alignment;
C. Only MAPQ Zero Reads: all reads above site have mapping quality (MQ) 0;
D. Only MAPQ Zero Alt-Supporting Reads: all reads supporting SNP in alignment have MQ 0;
E. Low Depth: total DP is less than 2;
F. Low Alt-Supporting Depth: total DP among reads supporting SNP is less than 2;
G. Low MAPQ Reads: the average of the mean MQ reference-supporting reads and the mean MQ of the SNP-supporting reads is less than or equal to 20;
H. Low BaseQ Alt-Supporting Reads: the average base qualities of reads supporting the SNP is less than 20;
I. Other: all other SNPs.

This provides a somewhat sorted categorization of SNPs from harder to call with short reads alone, to easier to call with short reads.

### Structural Variant Discovery with Long Reads

In order to measure structural variant (SV) discovery at various levels of coverage, each set of long reads (PacBio and ONT) was downsampled from 1-10x, and then run through Sniffles v2.2 (33) to call SVs (`LowCoverageSniffles.wdl`). Parameters were chosen to be more sensitive at lower coverage (see Supplementary Fig. 18 for a comparison of low-coverage flags vs. the defaults). In particular, it was run with settings: `--minsupport` `1 --qc-output-all --qc-coverage 1` `--long-dup-coverage 1 --detect-large-ins True`. The ONT 14x control reads were run with the same settings, while the PacBio 30x control reads were run with just the flag `--detect-large-ins True`. In all cases, the `--tandem-repeats` flag was used with the tandem repeats file for hg38 from https://github.com/PacificBiosciences/pbsv/blob/master/annotations/.

A custom script was then used to clean the data (`CleanSVs.wdl` from (93)) which splits multiallelic sites, recomputes the SVLEN annotation (SV Length) to standardized positive values and fills in missing values, strips any filters to maximize recall, and removes events smaller than 50bp which fall under the short INDEL category (see above). Following this step, the cleaned VCF was run through a pipeline wrapping Truvari v.4.3.1 setting `dup-to-ins` and `passonly` to true, and using `pctseq` equal to 0 in order to have a more equal comparison with the short read control which do not report basepair resolution of differences in large insertions. We show the “fuzziness” of Truvari’s comparisons does not given us an unfair advantage at lower coverage: we observed the distributions of start and end positions, along with inserted sequence similarity, match those at high depth (Supplementary Fig. 23–28). The SV VCF was also restricted to events with AC at least 1 using `bcftools view --min-ac 1`, so reference calls were removed before running Truvari. These were compared to the NIST Q100 v1.1 SV truth set for HG002 (47), which is part of the T2T-Q100 project (9), an initiative attempting to create perfect telomere-to-telomere assemblies (94) and polishing them with new tools like in the Telomere-to-Telomere project (95). The truth VCF was also restricted to events of size at least 50bp, and multiallelic sites were split, in the same way that the comparison VCFs were cleaned, except filters were kept in the truth.

We considered only events with at least 90% overlap with NIST’s truth high confidence BED file (labeled `stvar.benchmark.bed`) for our benchmarking statistics. After obtaining TP, FP, and FN labels in this way, we derived precision, recall, and F1 statistics overall, and also when restricting to events with positive percent overlap in particular regions of interest. When considering overlaps with specific regions, we followed the convention that insertions should have reference length given by the length of the inserted sequence, to avoid nearby misses due to ambiguity in the exact starting base, particularly when considering events discovered by short reads. As a control, we compared the performance to a callset curated from GATK-SV (39) calls on an Illumina 30x BAM from (96), with the HG002 sample extracted with `bcftools view -s HG002` and processed with the same cleaning and benchmarking pipeline.

### Hybrid Genotyping of SVs

Given SV calls made from the long reads, we regenotyped them using the short reads in two ways. The first, using vg v1.61.0 (97), started with 4x long reads (PacBio and ONT) mapped to the GRCh38 human reference, and created a sample-specific graph reference to map the short reads against. The full WDL pipeline steps are included in the repository described in the Code Availability section under the corresponding names. The overview of the pipeline implementation is as follows:

1. `GiraffeIndex.wdl` – The Sniffles SV calls were used to augment the linear GRCh38 human reference to create a graph reference, using `vg construct` in SV mode, `vg index`, `vg gbwt`, and `vg minimizer`. See the source code for the specific flags used in each command.
2. `GiraffeMap.wdl` – The short reads FASTQs were mapped to this sample-specific graph reference using `vg giraffe` (98).
3. `VgCall.wdl` – The SV calls were regenotyped via the short-read graph alignments using `vg pack` `--min-mapq 5 --trim-ends 5` and `vg call` `--ploidy 2 --vcf` (99).

The second genotying process used Paragraph, implemented in the `Paragraph.wdl` pipeline using the following steps:

1. Run the `CleanVCFParagraph` script to format the SV callset in ways Paragraph can handle, e.g. converting duplications into insertions, setting the reference field of symbolic deletions, and ensuring the first letter in insertion ALT field matches the reference. Calls close to the edge of chromosomes were excluded.
2. Run the `multigrmpy.py` script provided by Paragraph on the cleaned SV VCF and the short-read BAM to produce the final regenotyped SV VCF.

The output from both processes was then run through the same cleaning and benchmarking process as described in the last section. The resulting performance statistics were plotted relative to their short read coverage, and compared to the pure short read and pure long read controls, the latter of which were obtained as in the last section.

### Phasing Variants

Phasing was performed using a collection of tools for comparison and matched appropriately to the datatype for each experimental group. The exact code used for computing this is included in the repository described in the Code Availability section. The workflows used are:

1. `PhaseVCF.wdl` – This is a wrapper around `whatshap phase` (49) after running `bcftools` `+fixploidy` to avoid a bug in the tool if any haploid variant calls are passed through (like on chromosomes X and Y), even if they are not considered in the analysis. This was run on the short-read control SNP and INDEL VCFs produced via DRAGEN to improve on the default phasing outputs from DRAGEN (see Supplementary Fig. 36 for comparison), using just the short reads alone. We did not phase these together with the GATK-SV SV calls because there were missing sequences making it impossible to phase using the reads. A comparison of the effect of including SVs or not in the phasing process for the long-read experiments is included in the supplement (Supplementary Fig. 41), where a small benefit to including the SVs is observed but not as big as the difference between the short-read controls and other experimental groups.
2. `HiPhase.wdl` – This is a wrapper around HiPhase (51), a tool developed to use long reads to phase. It was run on the blend-seq experimental group SNP and INDEL VCFs merged with the SV VCFs called from their long reads, and with the long-reads-only controls using just the associated long reads on their SNP and INDEL VCFs merged with their SV VCFs. These results are labeled as using the phasing tool Hiphase in the figures.
3. `LongPhase.wdl` – This is a wrapper around Long-Phase (50). This was run same as HiPhase on the blend-seq experimental groups using their combined SNP, INDEL, and SV calls as input and just the long reads for phasing, as well as the long-reads-only experiments. We note that LongPhase allows for multiple BAM inputs, meaning we could use both a short- and long-read BAM, but our comparisons for our experimental groups showed worse performance than just using the long reads (Supplementary Fig. 39 and Supplementary Fig. 40).

To evaluate phasing performance, the `BenchmarkPhasing.wdl` pipeline was used, provided in the repository described under Code Availability. The pipeline performs the following calculations:

1. Remove SVs if present (and used to help phase the other variants), then use `whatshap compare` with the phased VCF produced above to compare it to the fully-phased NIST Q100 V1.1 short variant VCF. This produces statistics which require a truth VCF to compute, like switch error counts. Also run `whatshap stats` to compute intrinsic statistics, like NG50.
2. A custom script using `bedtools` (100) divides the phase block bed file produced by `whatshap stats` at switch errors produced by `whatshap compare`, and then recomputes the NG50 on this refined “corrected” set of phase blocks to result in the NGC50 (“corrected NG50”) statistic, similar to the method used to produce this as in vcfdist output (52).

#### 0.1. Data Sources

For each technology, we started with a high coverage set of reads and then downsampled to the approximately desired coverage using `samtools view -s`.

The initial set of reads for each technology came from the following sources:

- Illumina: We took two replicates of HG002 sequenced at the Broad on a NovaSeq X machine, and merged into a 68x BAM. It was mapped using DRAGMAP v3.7.8. This was first downsampled to 60x, and then from there to all the desired coverages.
- PacBio: We used a 30x HG002 run on a Revio machine sequenced at the Broad for internal validation studies. It was mapped using pbmm2 (101, 102) with the flags `--preset CCS --sample NA24385_1 --strip --sort --unmapped`.
- ONT: We used the set of reads from (37) which totaled around 14x, previously mapped using minimap2, and then downsampled from there, for all experiments involving “ultralong” (UL) ONT reads. This includes all of the blend-seq hybrid experimental groups using 4x ONT, in addition to the long-read structural variant discovery results. For the SNP discovery control line for 30x ONT, we found the other UL ONT reads from (37) to have dissatisfactory performance when combined to represent a reasonable control, so we used the reads from the HG002 T2T project (ht tps://github.com/marbl/HG002) and down-sampled to 30x. In particular, we merged the “pass” BAMs from the AWS server referenced at https://github.com/marbl/HG002/blob/main/Sequencing_data.md, then downsampled keeping 47% of the reads. We then realigned them with minimap2 using the flags `-x map-ont -L -K 3G` `--MD -I 8g -a -c --eqx -t 64 -k 17`.

See the Data Availability section for accessing the Illumina and PacBio read data used.

## ACKNOWLEDGEMENTS

This project was funded in part by Microsoft Corporation. F.C. was supported by the All of Us award (OT2 OD038121) and the Gabriella Miller Kids First award (5U24HD090743). The authors would also like to thank Broad’s Data Sciences Platform (DSP) for their helpful advice and guidance for this project. In particular, the authors would like to thank Steve Huang, Ryan Lorig-Roach, and James Emery (Broad DSP) for sharing their expertise on long read data, in addition to Mark Walker (Broad DSP) for insights on the short read SV calling pipelines and on structural variants. The authors also would like to thank Kiran Garimella and Chris Kachulis (Broad DSP) for helpful conceptual discussions in early phases of the project.

## CODE AVAILABILITY

Custom scripts and code specifically written for this project can be found at: https://github.com/broadinstitute/blend_seq_paper. This includes the WDL scripts mentioned in the Methods section, as well as URLs pointing to existing pipelines used. In addition, the custom `chop-reads` utility tool can be found at: https://github.com/rickymagner/chop_reads.

## DATA AVAILABILITY

The Illumina and PacBio read data used in this study is available under BioProject ID PRJNA1220034 and BioSample accession SAMN46707599. It can be found at https://www.ncbi.nlm.nih.gov/bioproject/1220034. All other data is from public references and cited appropriately.

## A: Cost Estimate Analysis

In this appendix we provide an analysis estimating the cost difference between the different sequence data generation processes to provide an accurate picture of the cost-savings potential of using blend-seq. We emphasize that the real cost of generating sequencing data is highly variable and constantly shifting, due to factors including changing costs in sequencer machines, chemistry price changes, demand-based multiplexing options, and human labor costs. Despite this, we use publicly available cost data based on offerings from various institutions to try to estimate different aspects of the cost in a simple model described further below.

### A.1. Methods

Because our focus is primarily computational, we make some simplifying assumptions about the laboratory processes for producing the sequencing data to arrive at our estimates. These are:

1. The cost of producing data on a sample for a particular technology at a particular coverage is affine in the desired depth, given by

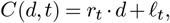

where *d* is the number of layers of coverage, *t* is a choice of sequencing technology, *r*_*t*_ is a rate of cost-per-layer for that technology, and *ℓ*_*t*_ is a constant overhead for library preparation costs. (In particular, we ignore subtleties around differences in labor costs, reagents, sample/DNA extraction, etc. beyond what is described as “library preparation” as a separate cost in some offerings.)
2. We assume the value *ℓ*_*t*_ is roughly constant percent of the total cost for a particular technology across different institutions, so that it can be estimated as a fraction of a “full cost” for institutions which do not separate library construction costs.
3. After fitting approximate values for *r*_*t*_ and *ℓ*_*t*_ across different technologies, we can approximate the cost of a blend-seq mixture between two technologies as

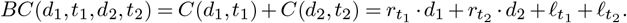

In other words, you pay the rate *r*_*t*_ for each layer of depth from technology *t*, and then add the respective technology overhead for library preparation.

With this framework, we found a list of institutions offering various products for whole-genome sequencing with the technologies used in this paper. Because we found it rare for an institute to offer all of our technologies used, some values in the table are missing as they are unavailable.

In order to normalize the cost across organizations, we need to calibrate how the product offer corresponds to coverage. In cases where the product offered is framed in terms of number of reads produced, we convert to approximate depth using the formula

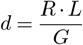

where *R* is the number of reads produced, *L* is the average read length, *d* is the depth, and *G* is the length of the reference genome (we use *G* = 3, 000, 000, 000 for our human reference).

Furthermore, a number of organizations did not separate library cost out of the total cost. In this case, we use assumption #2 above, first estimating the ratio *ℓ*_*t*_*/C*(*d, t*) across institutions where the price is split. Then, averaging those, we use this mean value as an assumption on the fraction of the library preparation costs for that institution, and fill the table values appropriate. Library preparation costs inferred in this manner are marked with an asterisk (*).

### A.2. Conclusion

Cost estimates for different technologies across different institutes offering services are provided in Supplementary Table 1. From this, using the above method, we derived estimates for blend-seq costs in Supplementary Table 2. From here, we estimate that blend-seq using PacBio reads would cost 53-58% (mean 55.8%) the cost of a 30x PacBio run while only being 1.7-2.8x (mean 2.3x) the cost of a full-depth Illumina run. For ONT, we estimate blend-seq would cost 58-62% (mean 60.4%) the cost of a 30x ONT run while only being 1.8-1.9x (mean 1.9x) the cost of a full-depth Illumina run.

References used to tabulate the original cost table can be found in Supplementary Table 3. Note we provide some extra institutes to estimate the cost of Illumina library preparation, to allow readers to derive similar estimates for other blend-seq mixtures (e.g. 15x short reads plus 4x long reads).

**Supplementary Table 1.** Sequencing platform costs per institute. Each institute lists costs separately for ILMN, PACB, and ONT platforms. Entries with an asterisk are imputed using the method described in the main text.

| Institute | Platform | Library Prep | Cost/Layer | 30x Cost |
| --- | --- | --- | --- | --- |
| BCL | ILMN | \$93.20* | \$10.23* | \$400.00 |
| BCL | PACB | \$380.38* | \$43.32* | \$1,680.00 |
| BCL | ONT | N/A | N/A | N/A |
| Harvard Bauer | ILMN | \$31.00 | \$22.59 | \$708.68 |
| Harvard Bauer | PACB | \$906.00 | \$94.93 | \$3,754.00 |
| Harvard Bauer | ONT | \$428.00 | \$58.83 | \$2,193.00 |
| NIH NCI | ILMN | \$136.00 | \$19.33 | \$715.90 |
| NIH NCI | PACB | \$294.00 | \$67.33 | \$2,314.00 |
| NIH NCI | ONT | \$370.00 | \$60.87 | \$2,196.00 |
| CGM Northwestern | ILMN | \$240.00 | \$18.48 | \$794.30 |
| CGM Northwestern | PACB | \$720.00 | \$82.00 | \$3,180.00 |
| CGM Northwestern | ONT | \$437.50 | \$72.92 | \$2,625.00 |
| Huntmans (Utah) | ILMN | \$124.99* | \$13.71* | \$536.42 |
| Huntmans (Utah) | PACB | N/A | N/A | N/A |
| Huntmans (Utah) | ONT | N/A | N/A | N/A |
| Case Western | ILMN | \$125.00 | \$10.93 | \$452.81 |
| Case Western | PACB | N/A | N/A | N/A |
| Case Western | ONT | N/A | N/A | N/A |

**Supplementary Table 2.**
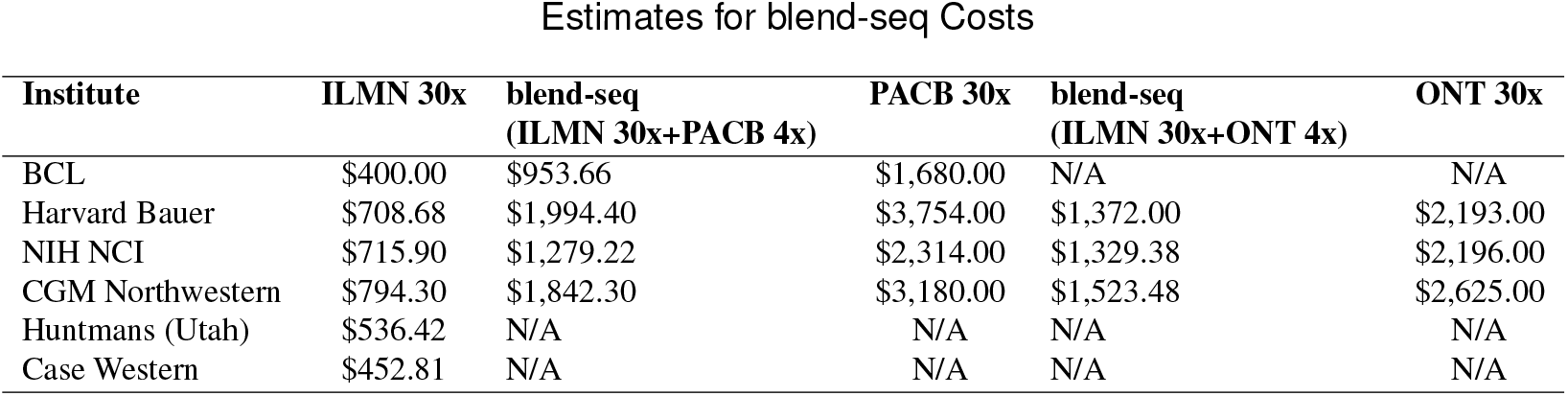
Cost estimates for 30x sequencing coverage and blend-seq pricing estimates using Illumina short reads and either PacBio or ONT for long reads.

**Supplementary Table 3.**
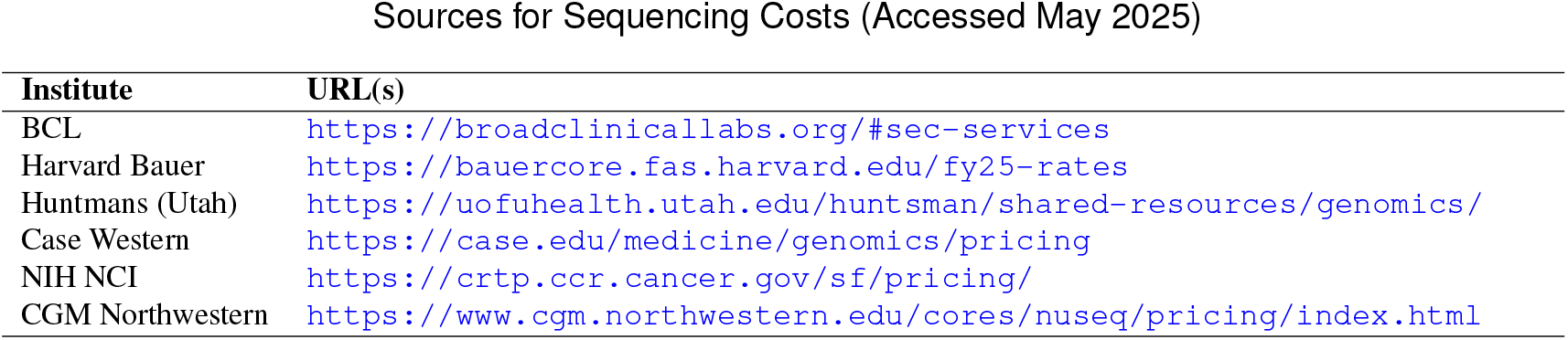
Source URLs used for retrieving sequencing cost data by institute. Accessed May 2025.

## B: Extra Figures and Tables

This section contains a collection of extra figures referenced in the main text.

### B.1. Extra Materials for Short Variant Calling Performance

**Supplementary Fig. 1.**
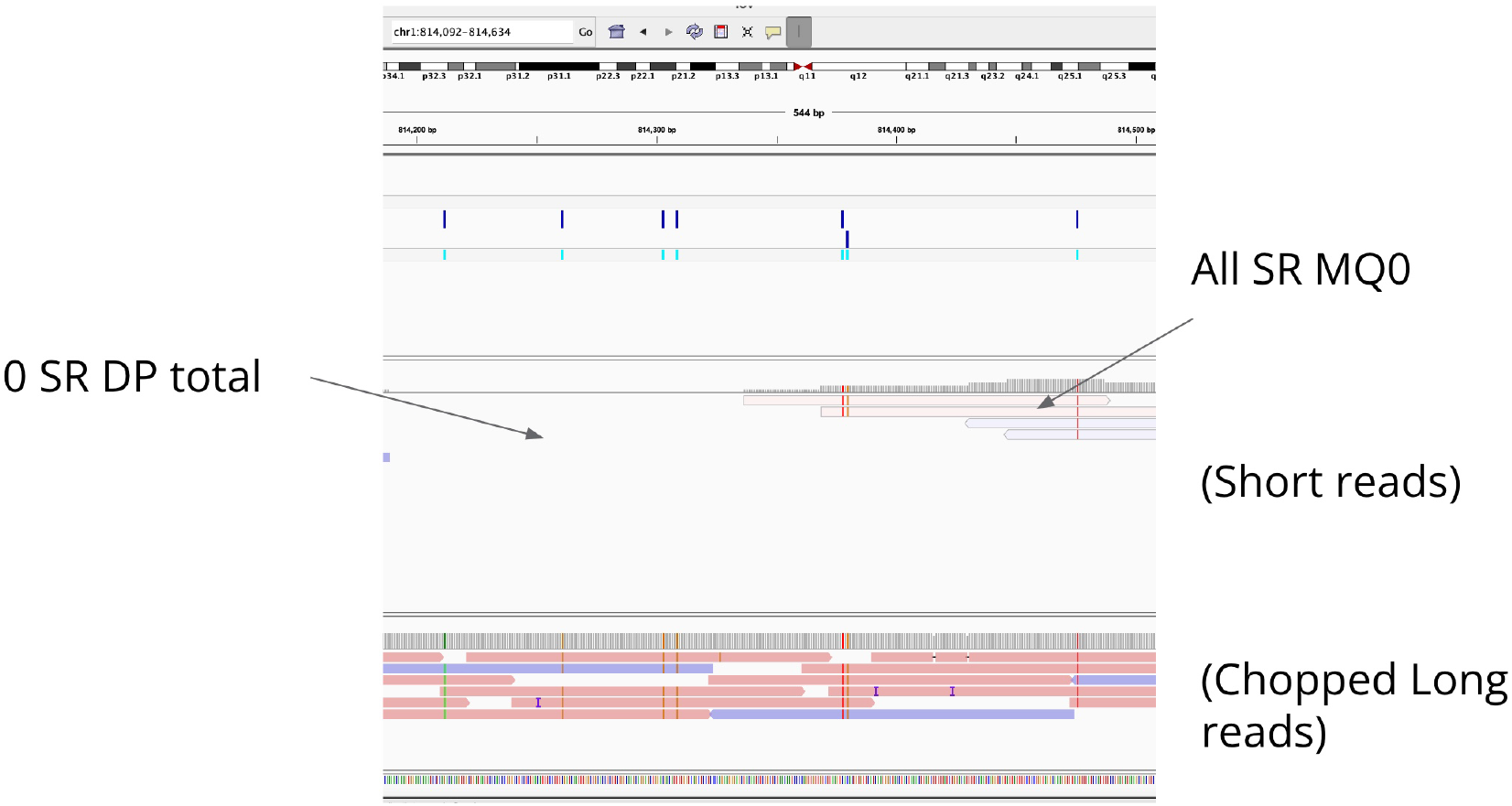
Example of variants with no short read depth or all zero mapping quality. This IGV screenshot demonstrates an example of a 0-DP-SR SNP (left) and a MQ0-SR-ALL SNP (right). The top set of reads are short reads, and the bottom are PacBio. The shading demonstrates that the long reads have good mapping quality, and we see by the mismatch colorings they can correctly pick up the actual SNPs around this locus.

**Supplementary Fig. 2.**
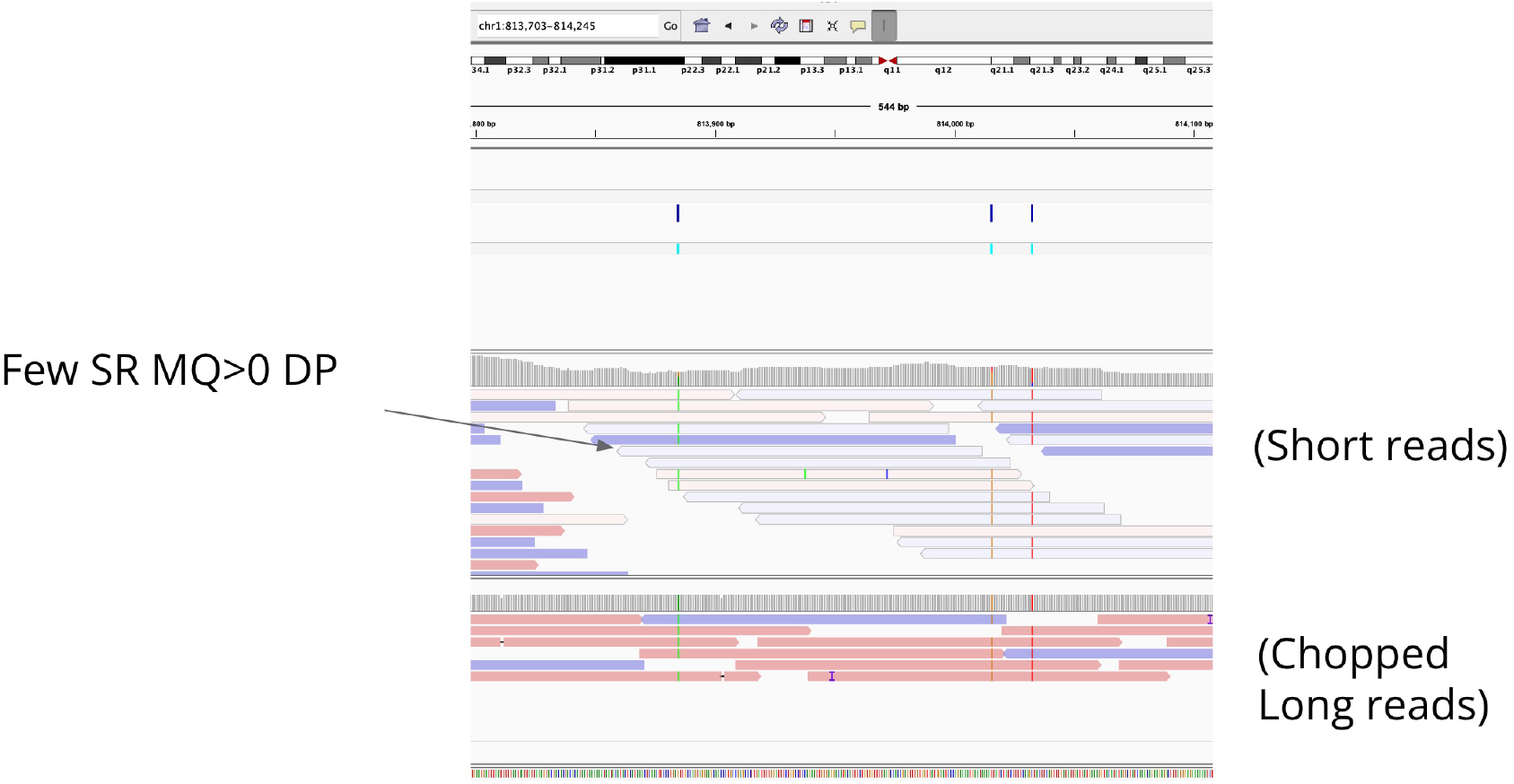
Example of variants with low mapping quality in short-read evidence. This IGV screenshot includes an example of a LOWMQ-SR-ALL SNP, as we see the short reads (top) have mostly mapping quality 0 here. The long reads (bottom) have no trouble aligning here with good mapping quality, and provide majority evidence for the SNP call in the hybrid.

**Supplementary Fig. 3.**
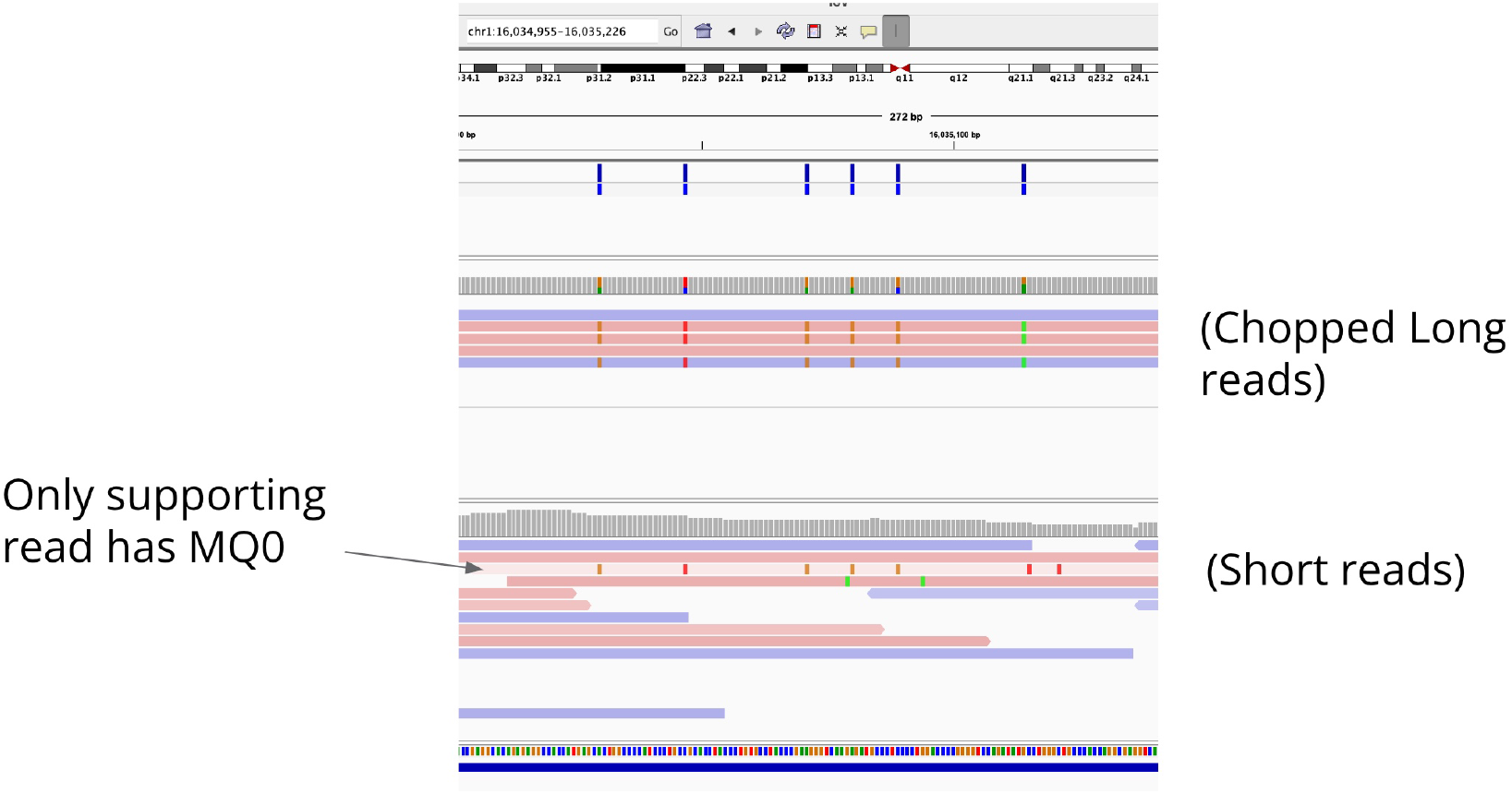
Example of variants with only supporting short reads have mapping quality zero. This IGV screenshot provides an example of a MQ0-SR-SUPP SNP, as the short reads (bottom) are split between mapping quality 0 and good mapping quality for the SNP-supporting and reference-supporting alignments respectively. We presume both the higher-than-usual concentration of SNPs in the sample plus low mappability of the region overall contribute to making the short read alignment for the SNP-supporting read zero. The long reads (top) have no trouble aligning here, even for the reads with a cluster of (real) SNPs.

**Supplementary Fig. 4.**
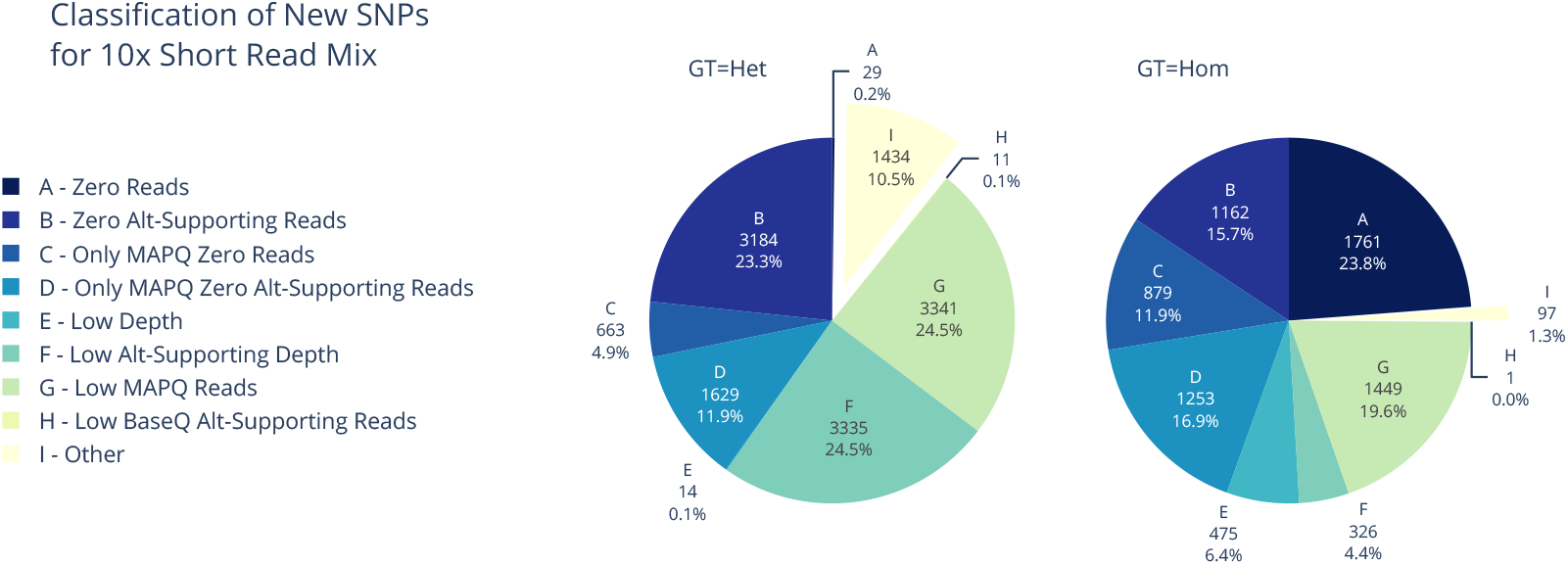
The breakdown of short-read false negative and hybrid true-positive SNPs for 10x short reads and 4x PacBio long reads in low mappability regions. We still see a large chunk of SNPs gained from the hybrid in low mappability regions are explained by lack of short read evidence (40% heterozygous and 2/3 of the homozygous sites).

**Supplementary Fig. 5.**
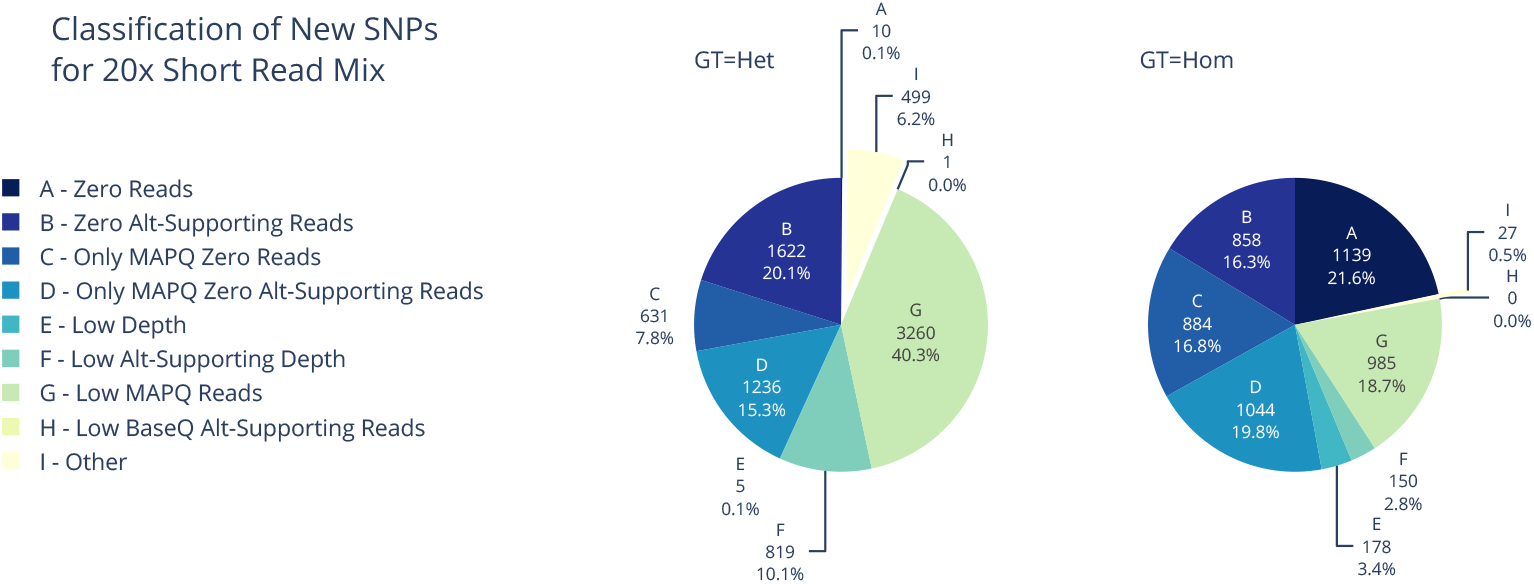
The breakdown of short-read false negative and hybrid true-positive SNPs for 20x short reads and 4x PacBio long reads in low mappability regions. We still see a large chunk of SNPs gained from the hybrid in low mappability regions are explained by lack of short read evidence (43% heterozygous and 75% homozygous).

**Supplementary Fig. 6.**
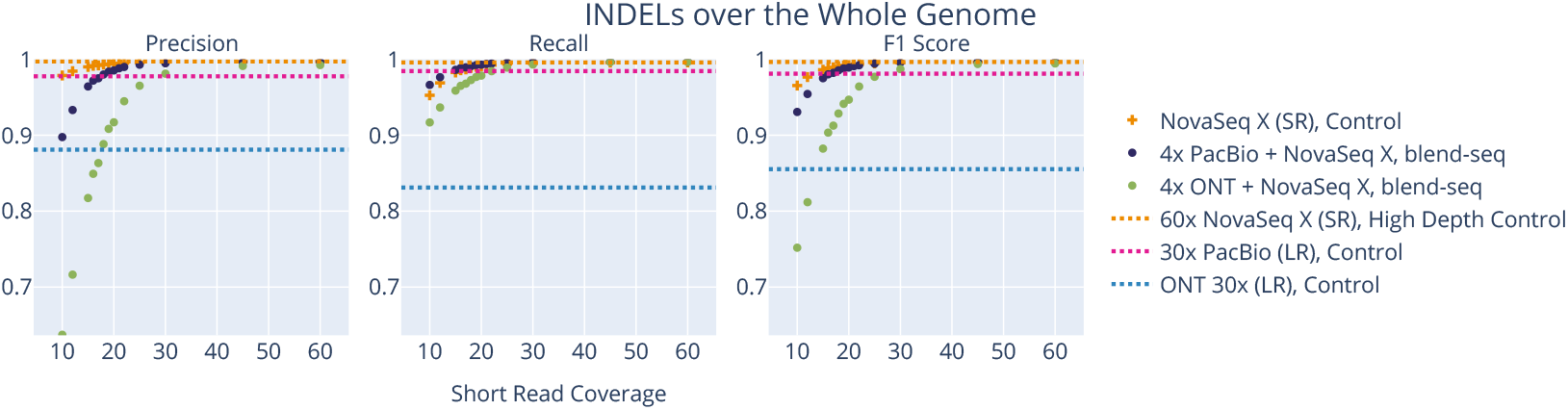
INDEL performance over the whole genome. The performance for INDELs over the whole genome looks about even for the hybrids vs the pure short reads, with ONT alone standing out with poor performance as a control.

**Supplementary Table 4.** INDEL performance by broken down by region. The blend-seq methods outperform the short read methods in recall overall, especially in hard-to-map regions. Precision lags a bit behind, possibly due to no hybrid filtering methods being applied to the results. Bold values are max values in column and shaded cells are those exceeding all control values in column (first three rows).

|  | WholeGenome |  |  | LowMappability |  |  | Exome |  |  |
| --- | --- | --- | --- | --- | --- | --- | --- | --- | --- |
|  | Recall | Precision | F1 Score | Recall | Precision | F1 Score | Recall | Precision | F1 Score |
| Illumina 15x (control) | .9826 | .9900 | .9863 | .8258 | .9168 | .8689 | .9813 | .9659 | .9735 |
| Illumina 30x (control) | .9946 | .9964 | .9955 | .8593 | .9354 | .8957 | .9938 | .9779 | .9858 |
| Illumina 60x (high depth) | .9959 | **.9972** | **.9965** | .8647 | .9364 | .8991 | .9958 | **.9780** | **.9868** |
| blend-seq 15x IL + 4x ONT | .9588 | .8173 | .8824 | .9171 | .5952 | .7218 | .9875 | .4681 | .6351 |
| blend-seq 15x IL + 4x PB | .9861 | .9643 | .9751 | .9197 | .8368 | .8763 | .9917 | .8602 | .9213 |
| blend-seq 30x IL + 4x ONT | .9932 | .9811 | .9871 | .9248 | .7253 | .8130 | .9938 | .8665 | .9258 |
| blend-seq 30x IL + 4x PB | **.9965** | .9949 | .9957 | .9241 | .8889 | .9062 | .9938 | .9625 | .9779 |
| 30x PB (high depth) | .9845 | .9776 | .9810 | **.9811** | **.9869** | **.9840** | **.9979** | .9273 | .9613 |

#### B.1.1. Updated DRAGEN Variant Calling Performance

This subsection includes small variant performance curves when using DRAGEN v4.2.7 instead of v3.7.8 as in the rest of the paper. We refer to the Methods section for details on why we chose to use the latter, but include the former latest version for completion.

**Supplementary Fig. 7.**
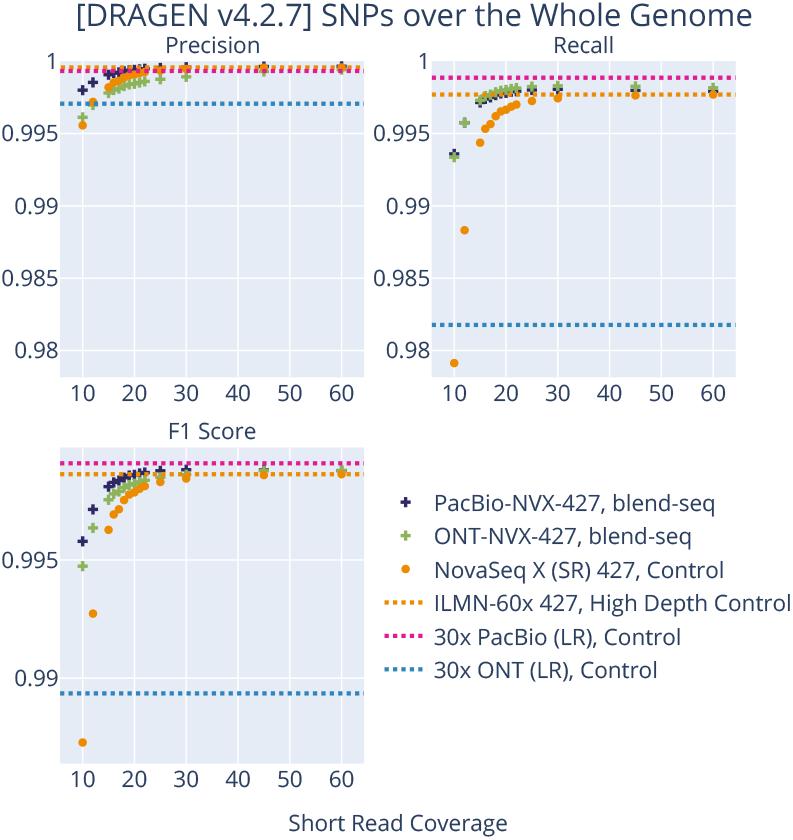
DRAGEN v4.2.7 SNP performance. When using the closed-source DRAGEN v4.2.7, we see the SNP performance on the whole genome still has higher recall for the blend-seq experiments, with higher performance over the high depth 60x short read control around 20x short reads + 4x long reads.

**Supplementary Fig. 8.**
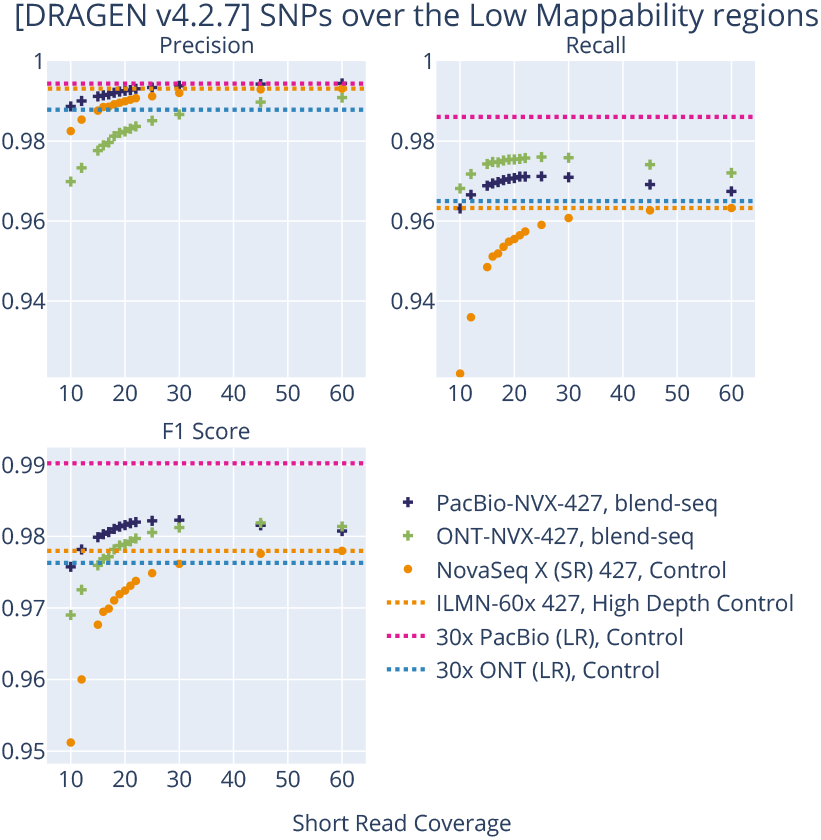
DRAGEN v4.2.7 SNP performance in low mappability regions. When using the closed-source DRAGEN v4.2.7, we see the biggest gains occur again in the low mappability regions for the blend-seq experiments, with them outperforming the short controls even at the highest depth for recall, for every combination of depths considered. The F1 scores look better than the high depth 60x short control for blend-seq starting around 15x short reads + 4x PacBio reads and 20x short reads + 4x ONT reads.

**Supplementary Fig. 9.**
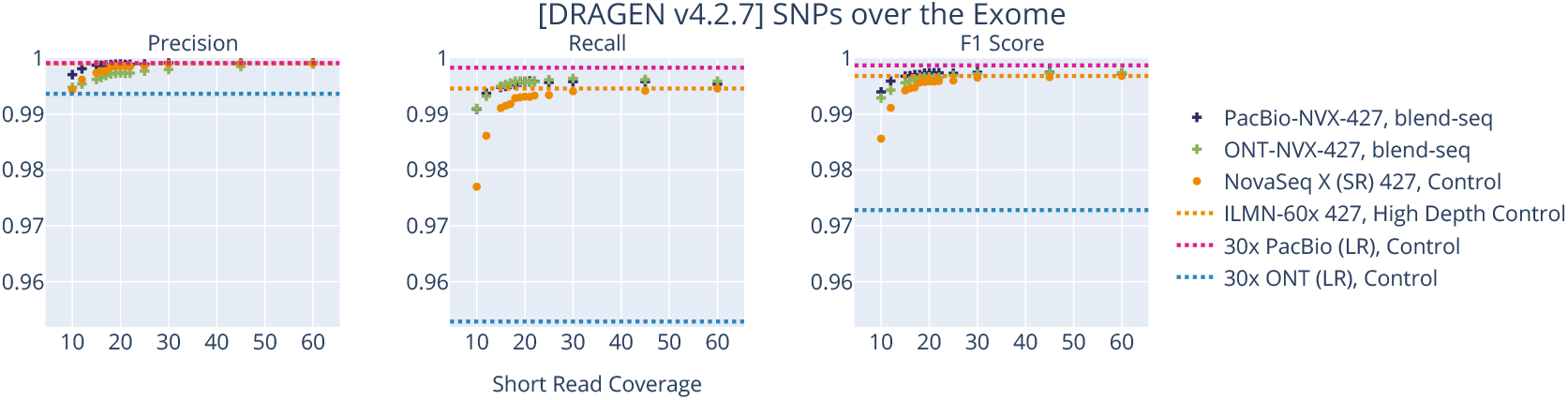
DRAGEN v4.2.7 SNP performance over the exome. When using the closed-source DRAGEN v.4.2.7, blend-seq still outperforms in recall around 15x short reads + 4x long reads.

**Supplementary Fig. 10.**
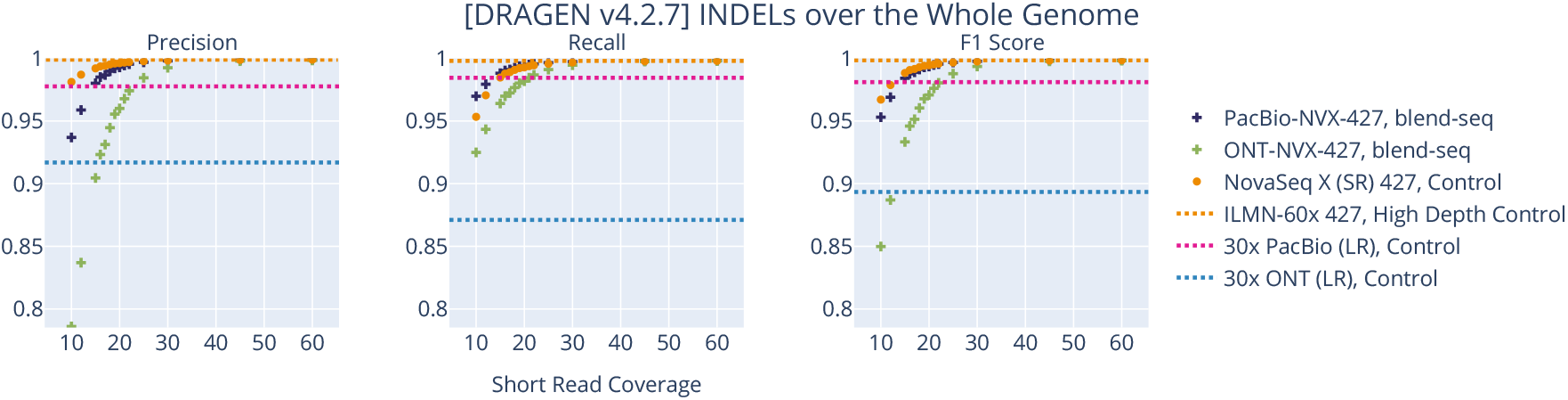
DRAGEN v4.2.7 INDEL performance. When using the closed-source DRAGEN v4.2.7, blend-seq shows a slight boost in recall over the short-read controls, but at a larger sacrifice in precision, suggesting a greater need for filtering in this setting.

### B.2. Extra Materials for Structural Variant Discovery

**Supplementary Fig. 11.**
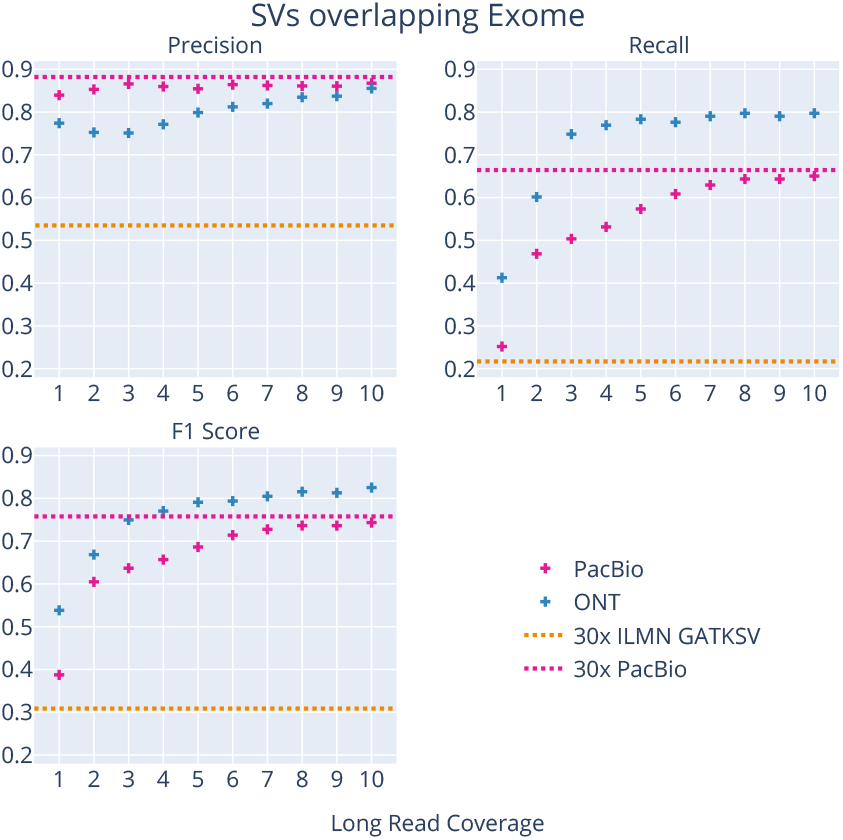
SVs overlapping the exome. The SV discovery performance boost for the pure long reads is maintained when restricting to events overlapping the exome, with the ONT reads having better recall than the PacBio reads.

**Supplementary Fig. 12.**
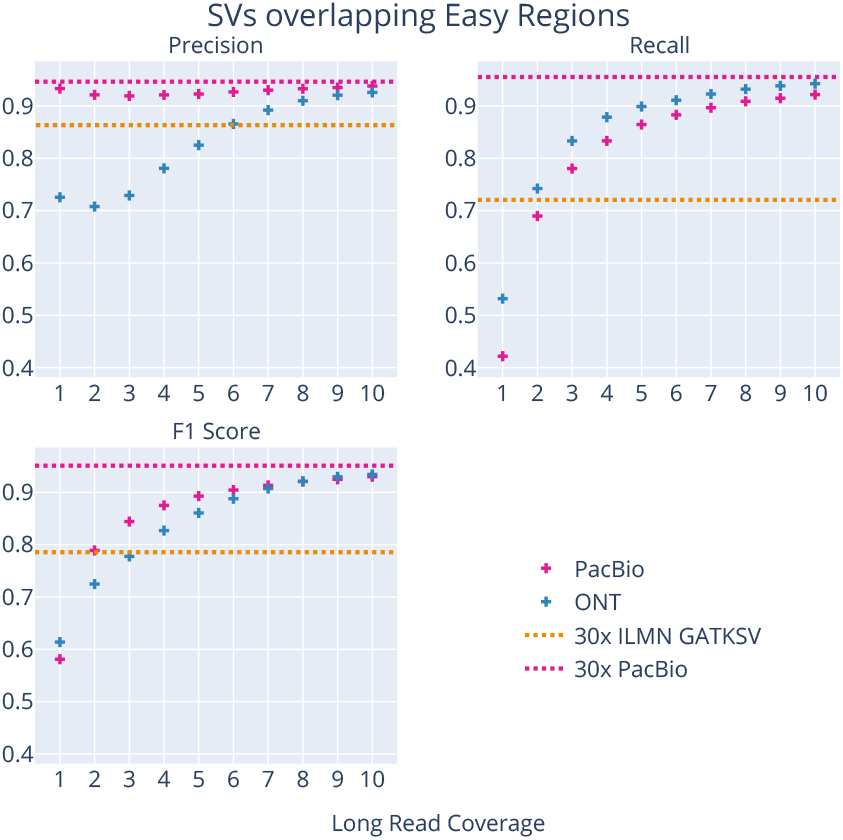
SVs overlapping easy regions. When restricting to the easy regions, we give the short reads the best region for them to perform well, and still see long reads outperforming in F1 starting at only 2-3x.

**Supplementary Table 5.** SV deletion discovery performance by region. With just a few layers, pure long reads vastly outperform short reads for SV discovery in terms of recall. This is powered by extra discovery coming from more difficult regions, although the short reads perform well in the Easy Regions. Bold values are max values in column and shaded cells are those exceeding the control value in column (first row). We note it had been previously observed that short-read perform comparably to long-read for SV deletions in easier regions (see e.g. (26)).

|  | Whole Genome |  |  | Easy Regions |  |  | Hard Regions |  |  | Exome |  |  |
| --- | --- | --- | --- | --- | --- | --- | --- | --- | --- | --- | --- | --- |
|  | Recall | Precision | F1 Score | Recall | Precision | F1 Score | Recall | Precision | F1 Score | Recall | Precision | F1 Score |
| Illumina 30x (control) | .3016 | **.8821** | .4495 | .8589 | .9160 | **.8866** | .2759 | .8721 | .4192 | .3333 | .6000 | .4286 |
| PacBio 3x | .6370 | .8621 | .7326 | .7841 | .9045 | .8400 | .6312 | .8689 | .7312 | .5067 | .7647 | .6095 |
| PacBio 4x | .6821 | .8650 | .7627 | .8377 | **.9180** | .8760 | .6766 | .8685 | .7606 | .5600 | **.7818** | .6526 |
| ONT 6x | .7325 | .8253 | .7761 | .8995 | .7608 | .8244 | .7275 | .8557 | .7864 | .7600 | .6543 | .7032 |
| ONT 7x | **.7390** | .8573 | **.7937** | **.9125** | .8247 | .8664 | **.7334** | **.8748** | **.7979** | **.7733** | .7397 | **.7562** |

**Supplementary Fig. 13.**
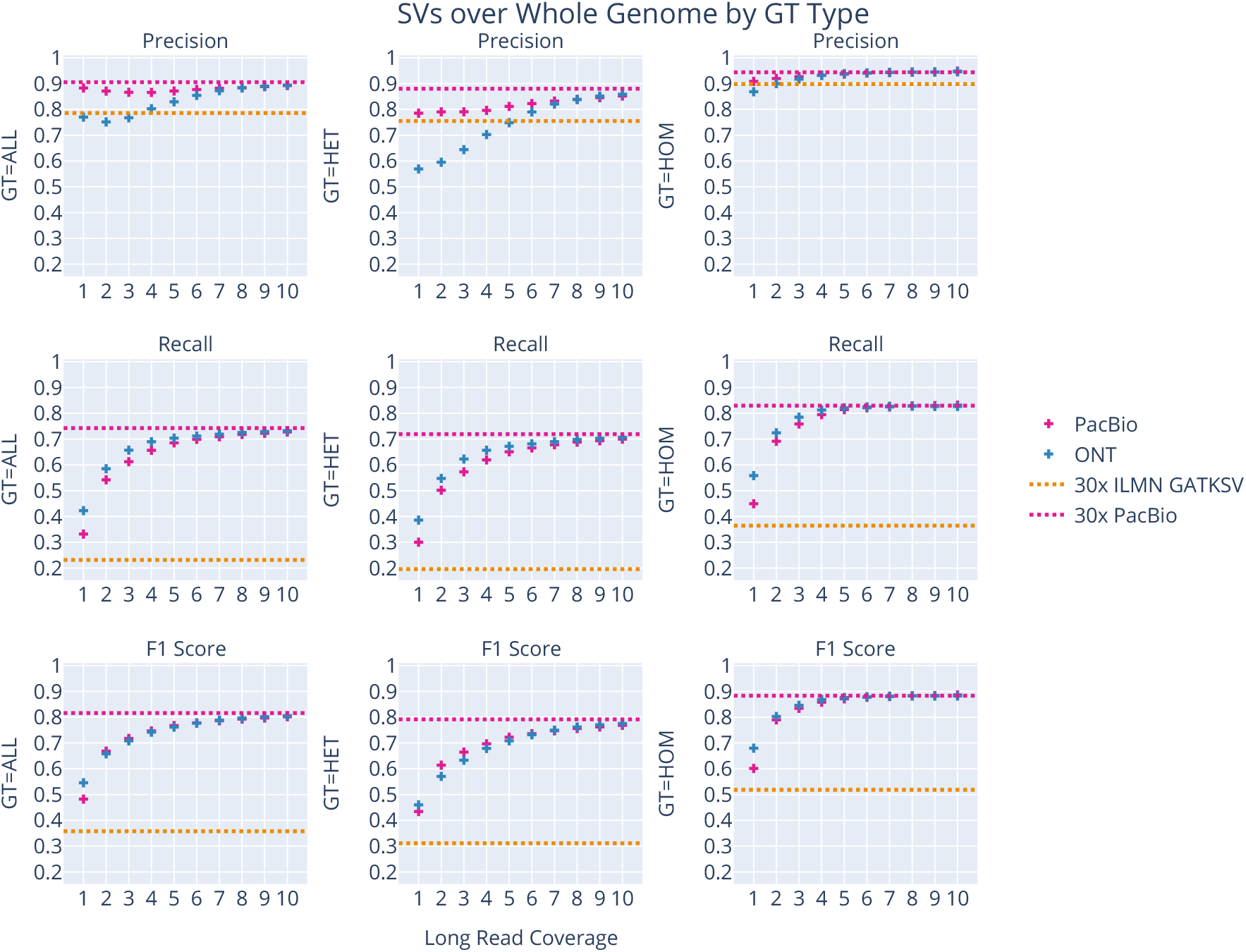
Breakdown of SV performance by GT type. When breaking down performance based on genotype category, we see performance slightly better for homozygous (HOM) variants than compared to heterozygous (HET) variants. For precision, we consider HETs in the query VCF, whereas for recall we consider HOMs in the truth VCF. It should be remarked that about 20% of variants in the confidence regions for the truth file are homozygous, so the overall trends skew slightly better than the HET categories. Nevertheless, the blend-seq curves sit above the control for recall and F1 already at 1x, and for precision the PacBio reads outperform at all depths while ONT breaks even around 5x. It is worth remarking that the HOM variant recall plateaus already around 4-5x for both technologies compared to the PacBio 30x “asymptote.”

**Supplementary Fig. 14.**
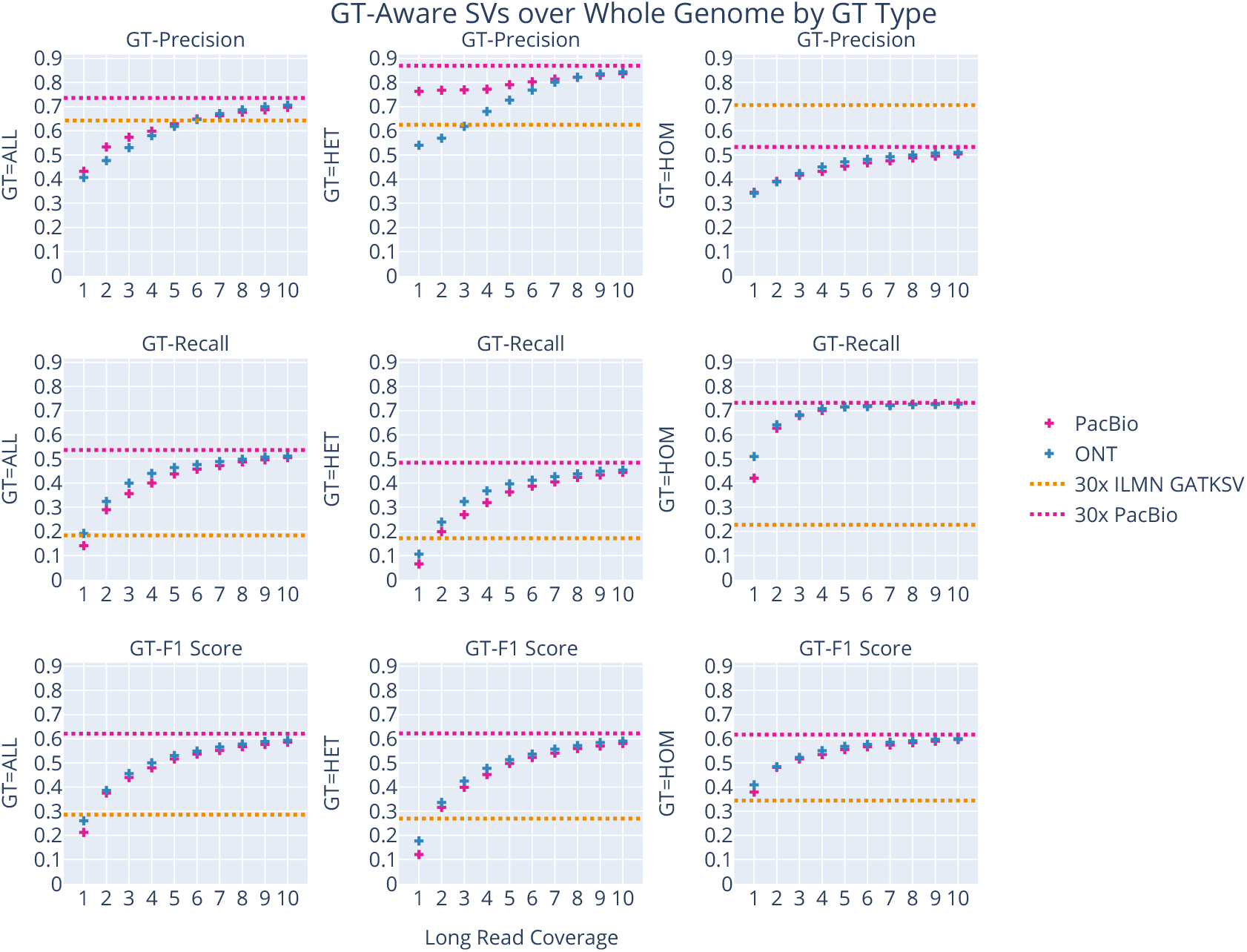
Breakdown of SV GT-aware performance by GT type. When looking at GT-aware performance by genotype category, we see a similar story as the non-GT-aware breakdown. The HOM variants yield better performance unsurprisingly, while the HETs lag a little behind. Here, the GT-Precision is always better for the short read controls for HOMs, balanced out by the much lower GT-Recall, shaking out to a better GT-F1 curve for the long reads when depth is at least 2x across all variant types.

**Supplementary Fig. 15.**
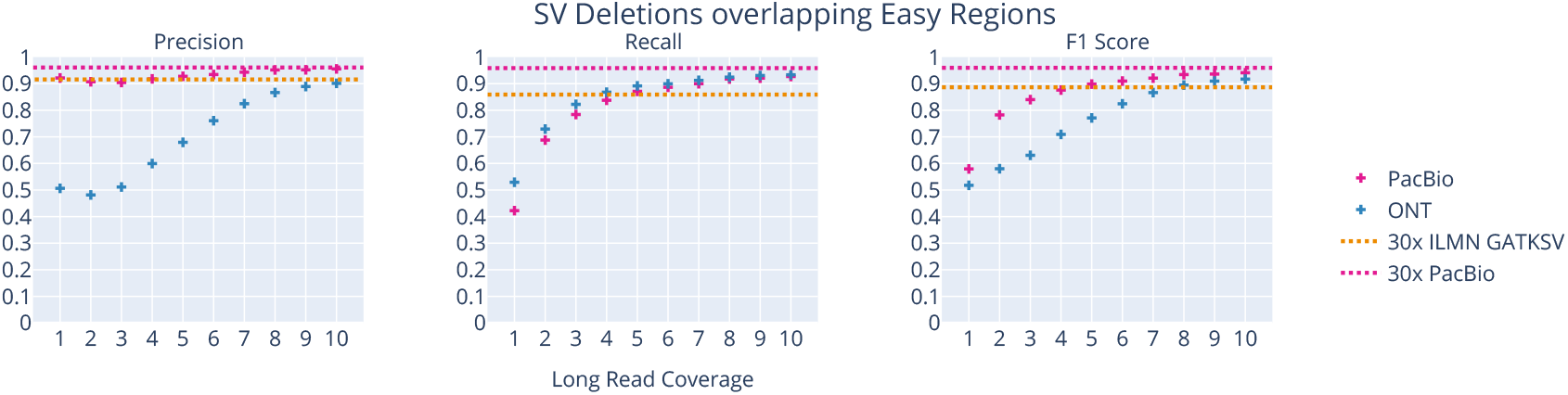
Performance of SV deletion discovery in easy regions. In the easiest possible regime for short reads (overlapping the easy regions and just considering large deletions), we see that ONT precision is the worst, but the PacBio precision still beats out short reads alone. The recall is also better for long reads at just around 4-5x depth. The small gap between short- and long-read performance for SV deletions in easy regions is consistent with previous findings (see e.g. (26)).

**Supplementary Fig. 16.**
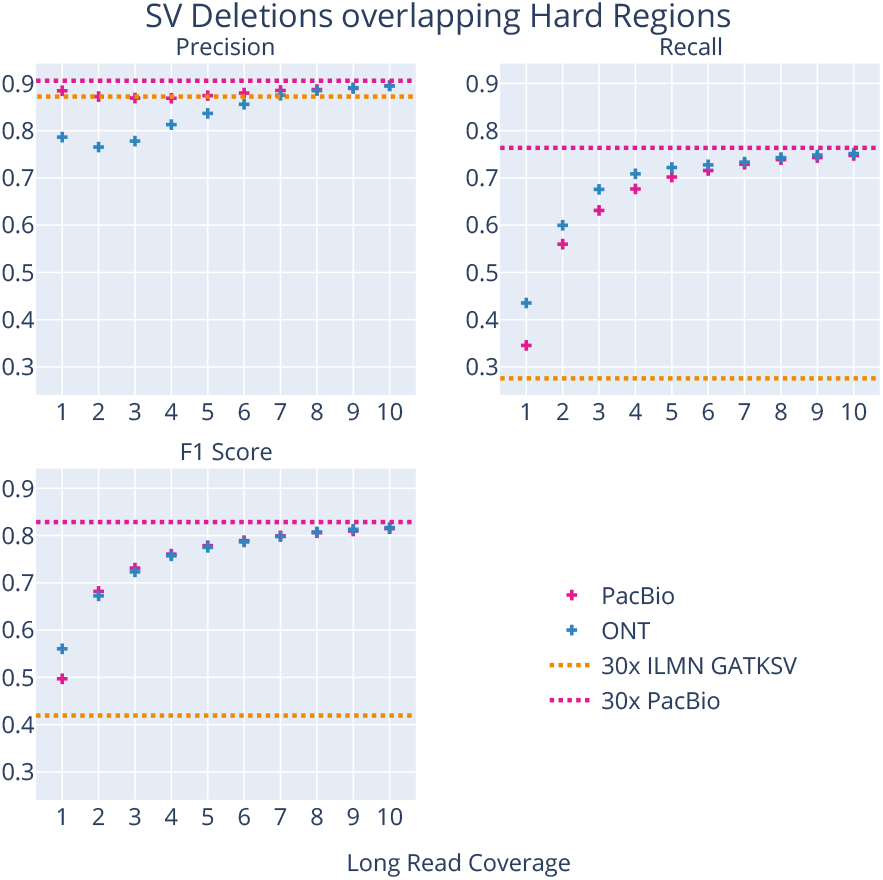
Performance of SV deletion discovery in hard regions. For events overlapping difficult regions, the long reads perform best when measured by F1 even for deletions. The precision is best with PacBio, followed by Illumina, and then ONT, but recall is much higher for long reads even at 1x coverage.

**Supplementary Fig. 17.**
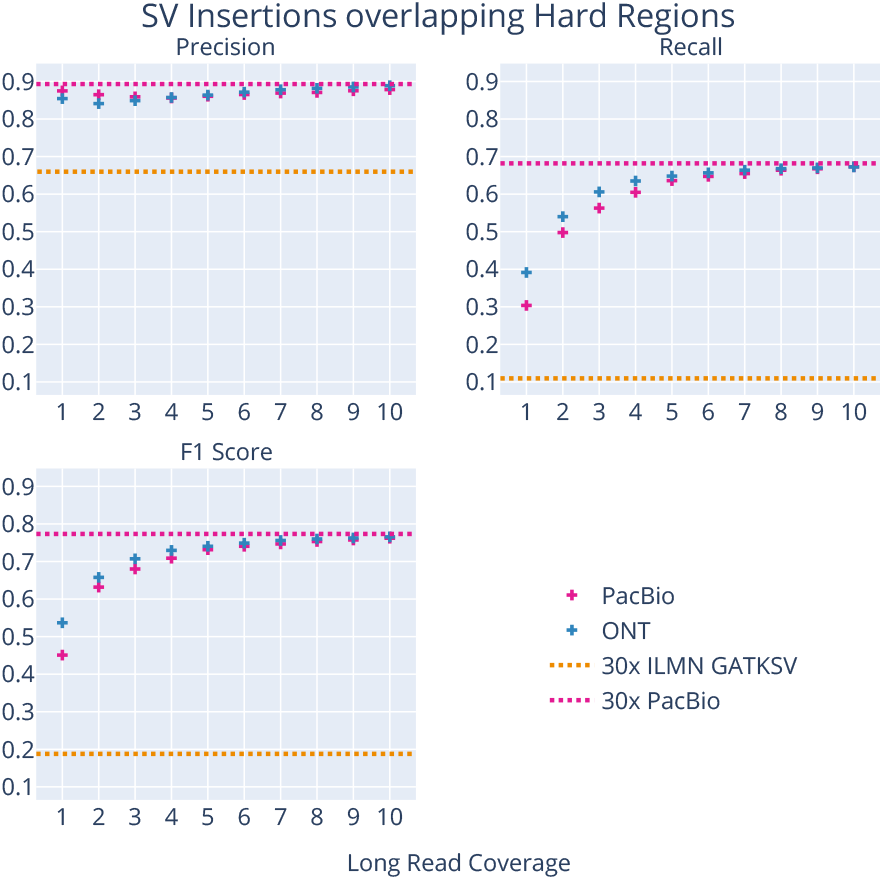
Performance of SV insertion discovery in hard regions. Insertions in difficult regions are the hardest for short reads to discover, and we observe long read discovery methods outperforming in every statistical category.

#### B.2.1. Other SV Calling Choices

In this subsection, we present some plots demonstrating other choices for processing the data for SV discovery.

**Supplementary Fig. 18.**
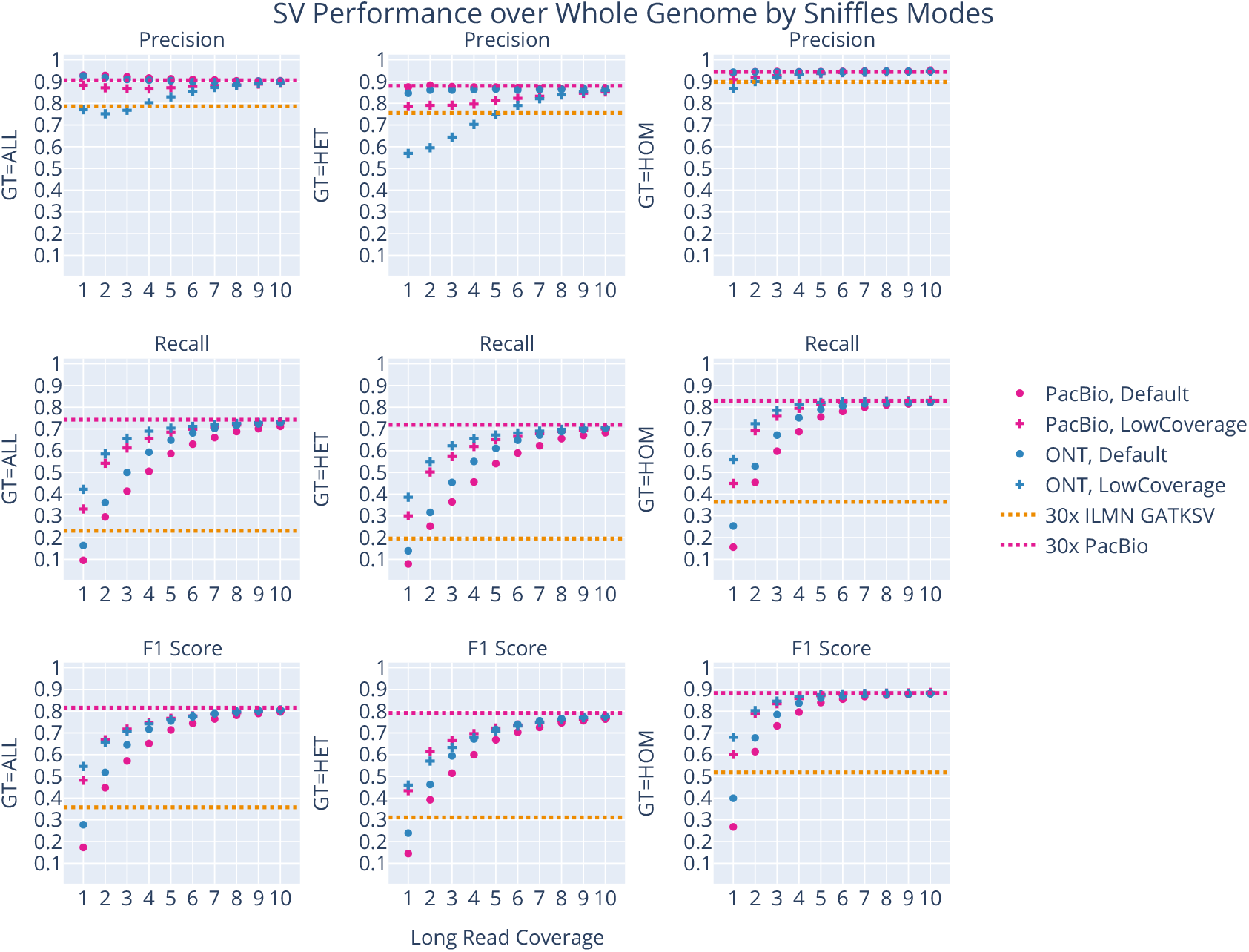
Comparison of Sniffles low-coverage mode to its default mode. We compared using Sniffles with its “low coverage” flags (cross symbols), as described in the Methods section, to running with its default parameters (circles) at the depths considered for SV discovery, with the 30x PacBio control run using the default parameters. The low-coverage flags outperform at lower depths for recall with a small loss in precision, which leads to an overall better F1. The gap between them shrinks at around 8x, and even sooner for ONT recall on HET sites. The drop in precision for using the low-coverage flags for PacBio still places it above the short read performance and is justified by the large boost in recall.

**Supplementary Fig. 19.**
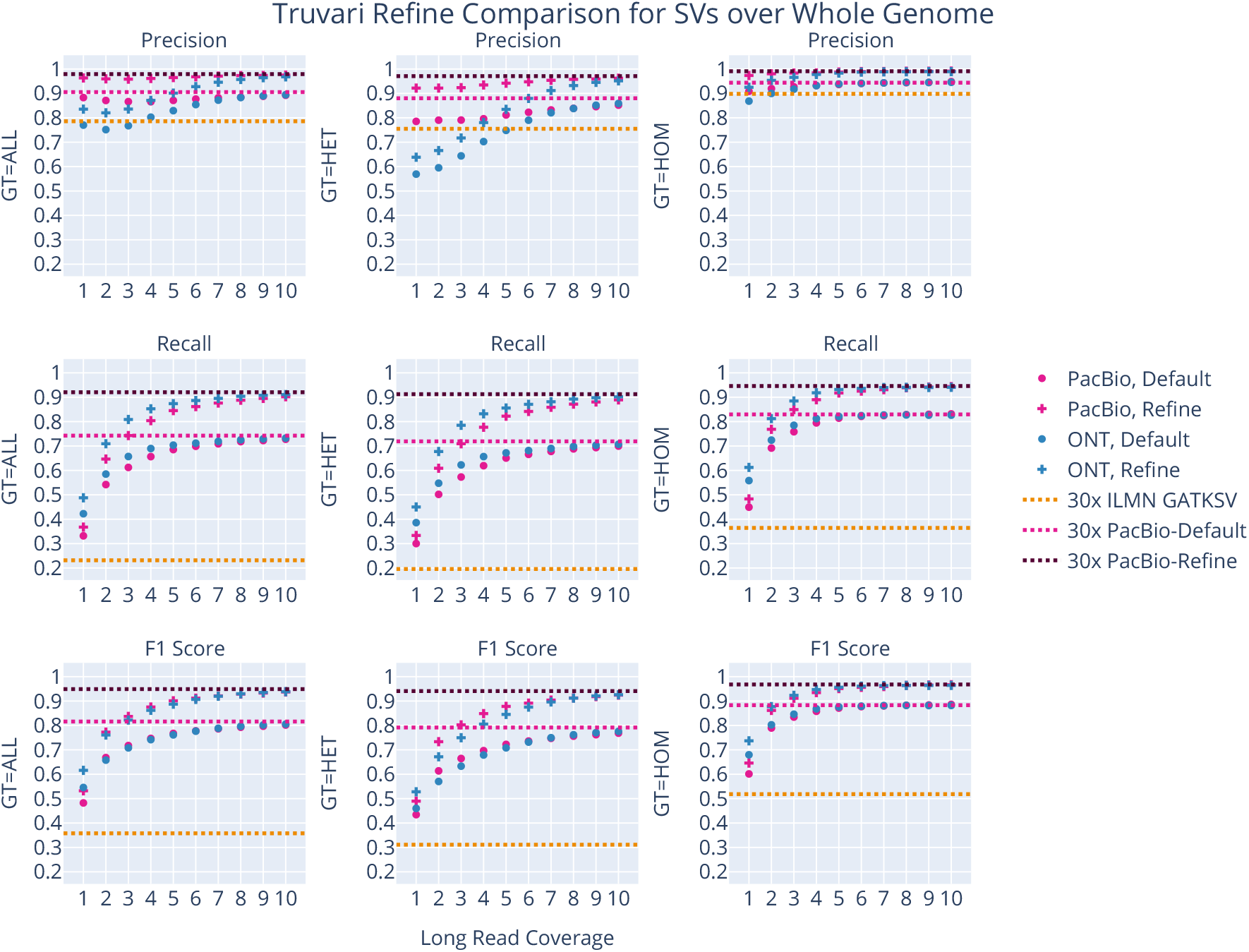
Truvari refine compared to default Truvari parameters. We compared two different methods of using Truvari to compute benchmarking statistics. The “Default” one labeled here reflects the techniques outlined in the Methods section, whereas the “Refine” one uses the Truvari refine command as an intermediate, as described in (62). We see the Refine version has better statistics, as was observed in ibid., but emphasize that this is a process which harmonizes the query and truth variants together, so is not reflective of a processing step that can be done in the absense of truth data. Despite this, we see similar trends in our benchmarking statistics using our “Default” evaluation although the exact numbers differ.

**Supplementary Fig. 20.**
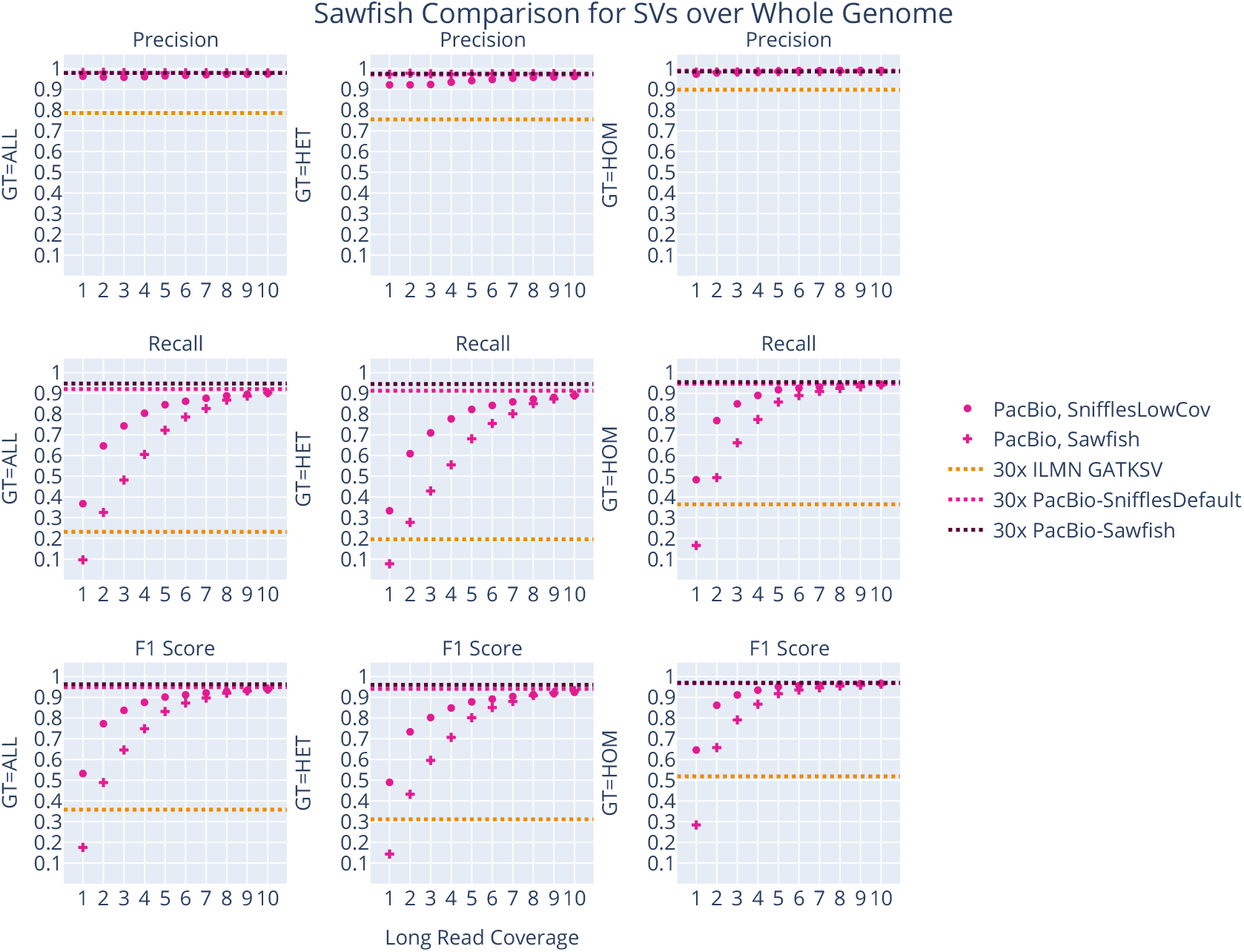
Sawfish compared to Sniffles in low coverage mode. We compared the performance of Sawfish (62), an SV caller written by PacBio for their reads, to the Sniffles low-coverage settings used in our paper. The version of Sawfish that was available at the time of our experiment did not allow changing core parameters like the number of supporting reads on the command line. Although (62) investigates the change in performance by coverage, it did not analyze the lower depths we investigated here. The two methods converge at higher depth, but at lower depth Sniffles with low-coverage parameters outperforms in recall for a small drop in precision. This becomes better F1 performance, especially at the lowest 1-7x depths.

#### B.2.2. Long-Read Genotyper Comparison

**Supplementary Fig. 21.**
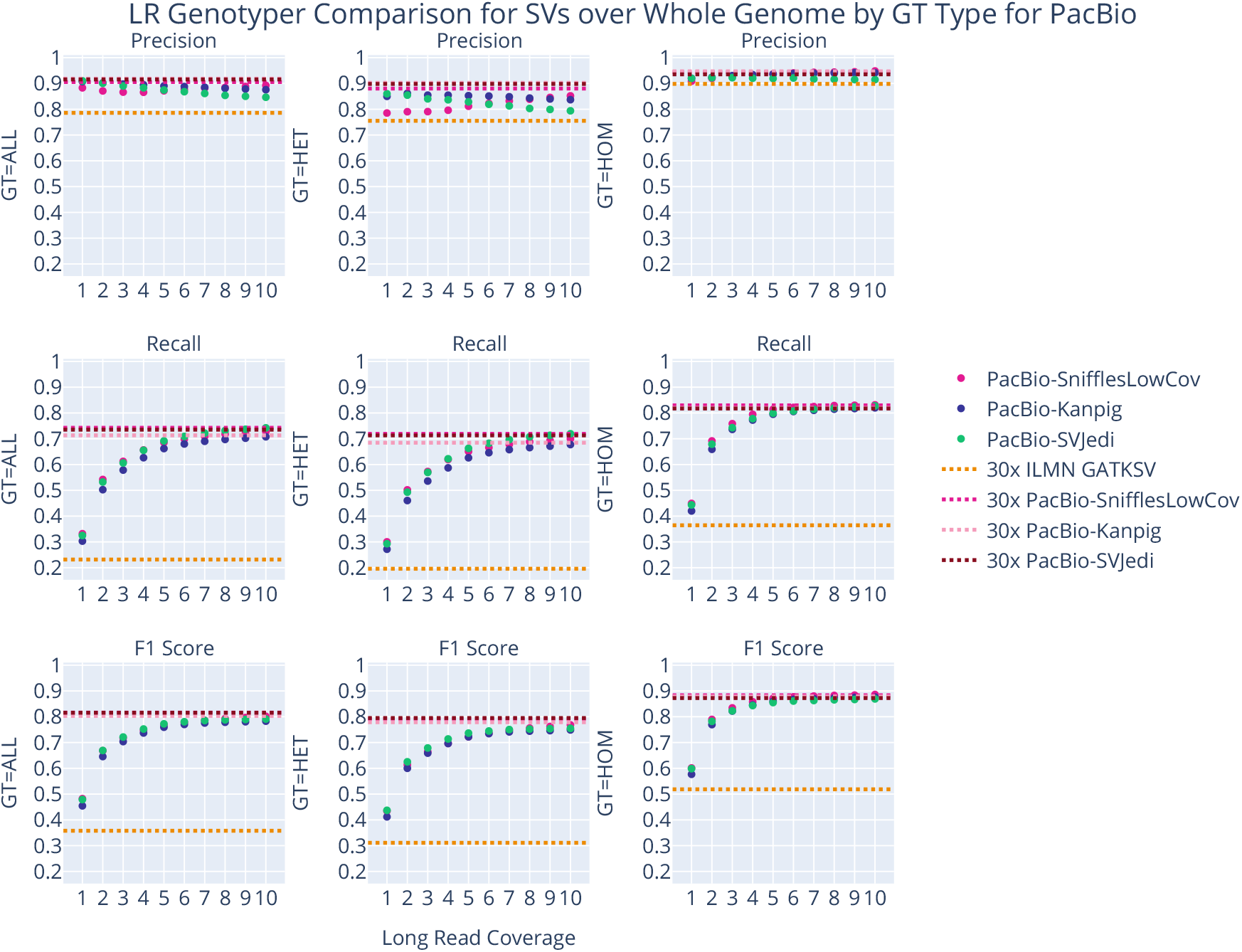
Comparison of long-read genotypers SVJedi (61) and kanpig (103) to Sniffles low-coverage outputs, for PacBio reads. Using the long-read SV genotypers SVJedi and kanpig on PacBio data yields very little difference over using the Sniffles low-coverage calls directly, except perhaps a small boost in precision for heterozygous sites. This tradeoff with recall is barely noticeable in the F1 statistics, justifying omitting these genotyper tools from the pipeline.

**Supplementary Fig. 22.**
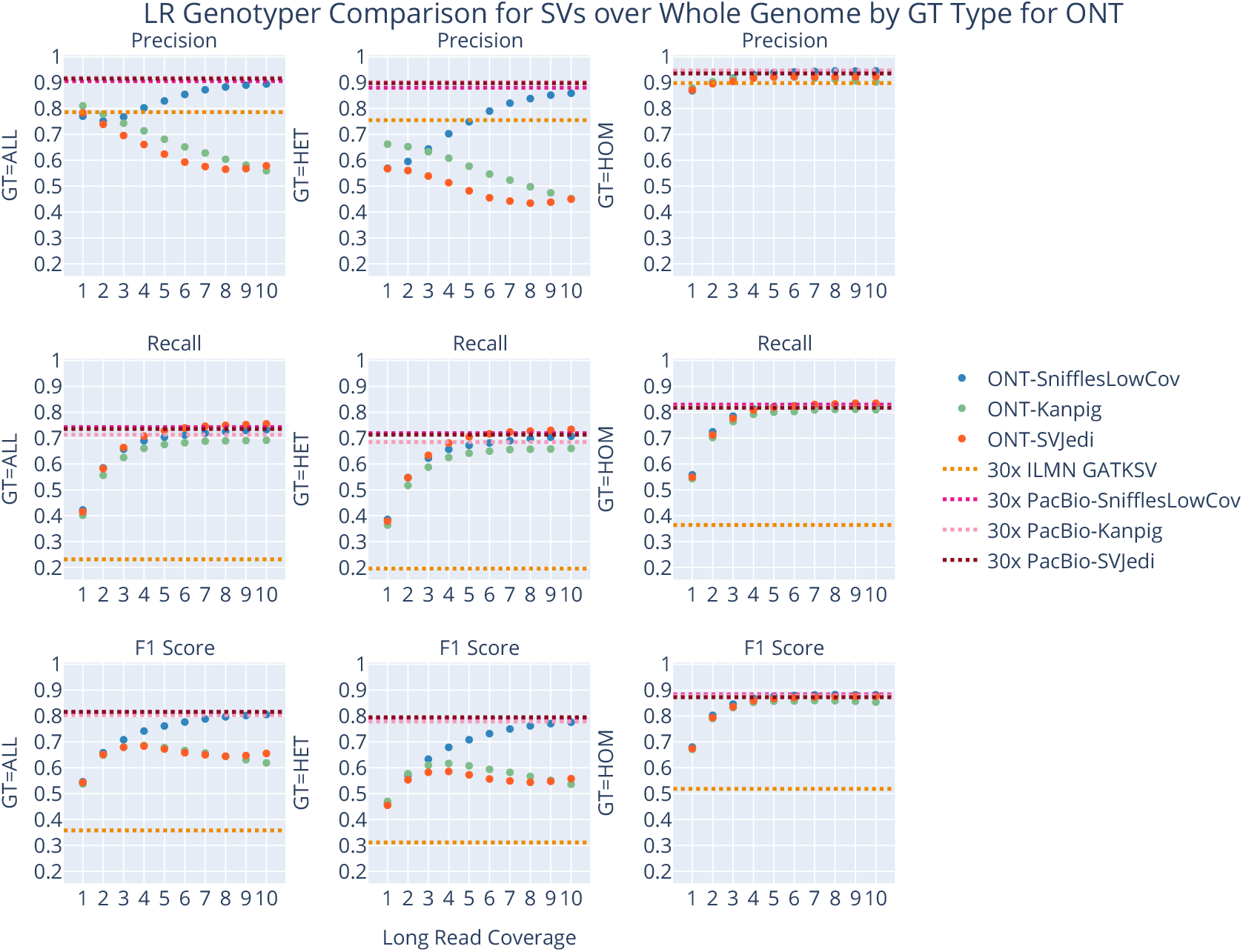
Comparison of long-read genotypers SVJedi and kanpig to Sniffles low-coverage outputs, for ONT reads. Using the long-read SV genotypers SVJedi and kanpig on ONT data reflects the fact they were not designed for this data-type, as performance drops dramatically as depth increases. This justifies using the Sniffles low-coverage calls directly rather than regenotyping them.

#### B.2.3. Truvari Statistics Distributions by Long-Read Coverage

**Supplementary Fig. 23.**
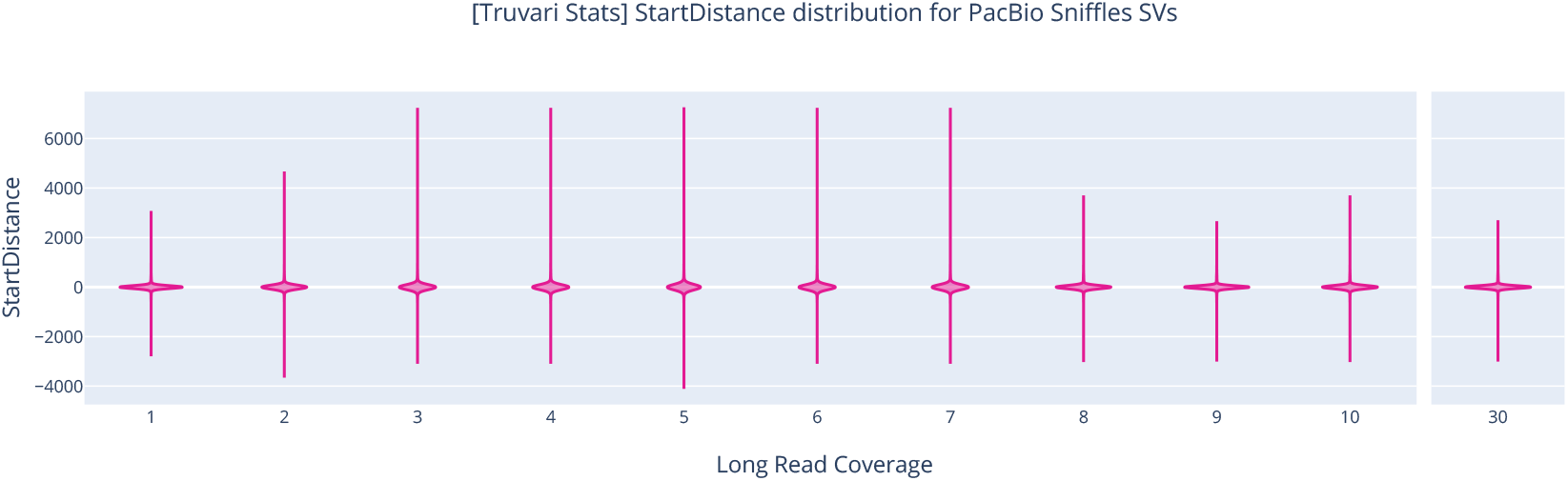
Distribution of Truvari’s “StartDistance” metric for PacBio reads. This measures how far the input VCF’s SVs’ start positions are relative to the start positions in the truth VCF for TP matches. We observe a relative consistency across long-read coverages, matching the pattern of the high-coverage control, suggesting the distribution captured at low coverage is reflective of the one at high coverage.

**Supplementary Fig. 24.**
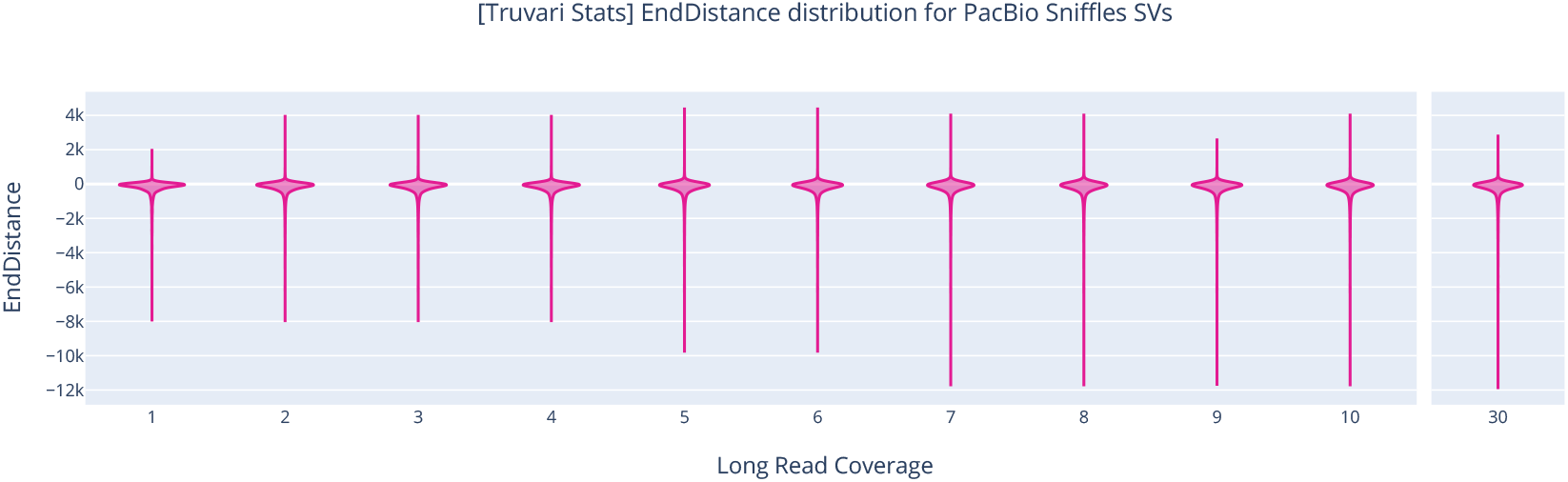
Distribution of Truvari’s “EndDistance” metric for PacBio reads. This measures how far the input VCF’s SVs’ end positions are relative to the end positions in the truth VCF for TP matches. We observe a relative consistency across long-read coverages, matching the pattern of the high-coverage control, suggesting the distribution captured at low coverage is reflective of the one at high coverage.

**Supplementary Fig. 25.**
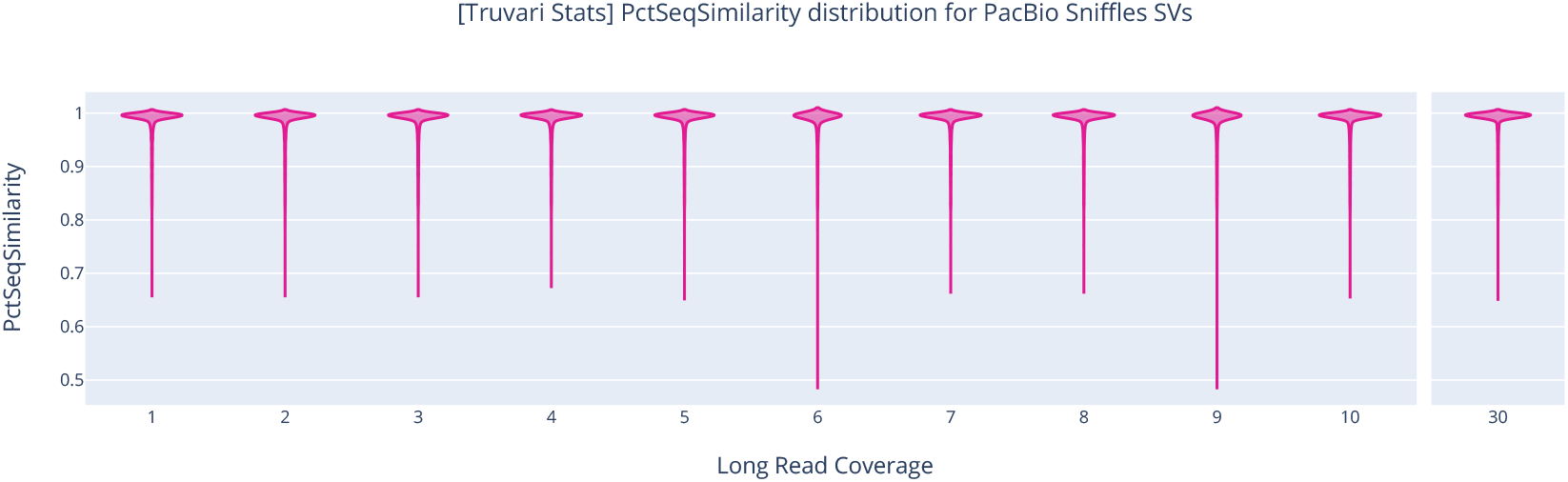
Distribution of Truvari’s “PctSeqSimilarity” metric for PacBio reads. This measures the relative similarity of allele sequences in the input VCF compared to the sequences in the truth VCF for TP matches. We observe a relative consistency across long-read coverages, matching the pattern of the high-coverage control, suggesting the distribution captured at low coverage is reflective of the one at high coverage.

**Supplementary Fig. 26.**
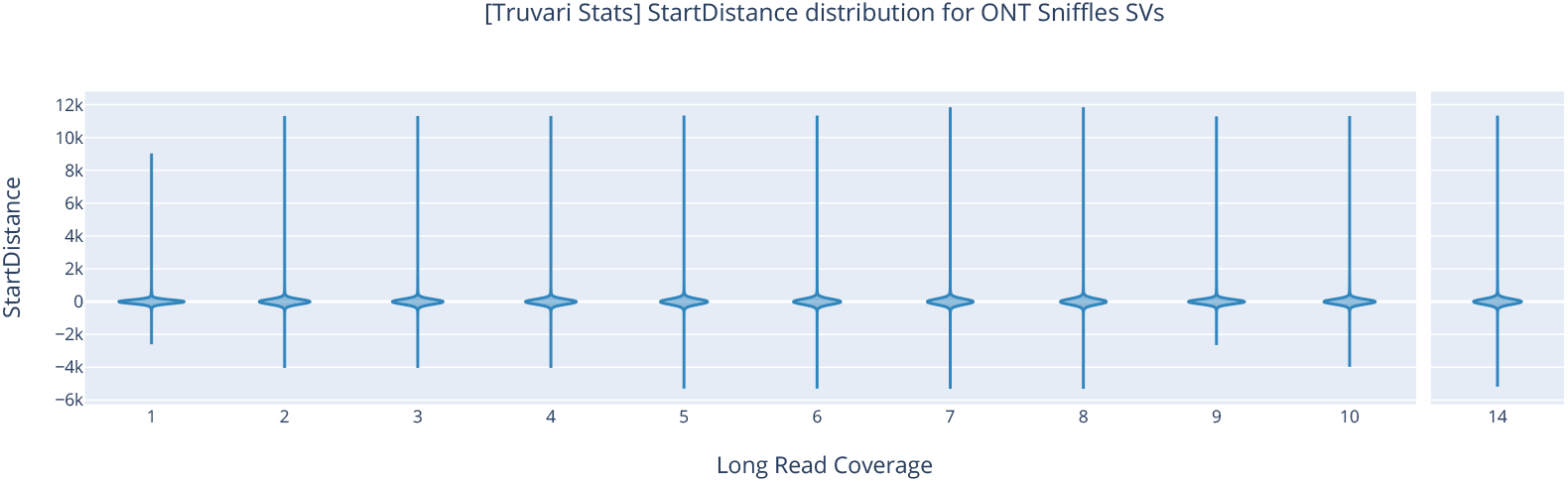
Distribution of Truvari’s “StartDistance” metric for ONT reads. This measures how far the input VCF’s SVs’ start positions are relative to the start positions in the truth VCF for TP matches. We observe a relative consistency across long-read coverages, matching the pattern of the high-coverage control, suggesting the distribution captured at low coverage is reflective of the one at high coverage.

**Supplementary Fig. 27.**
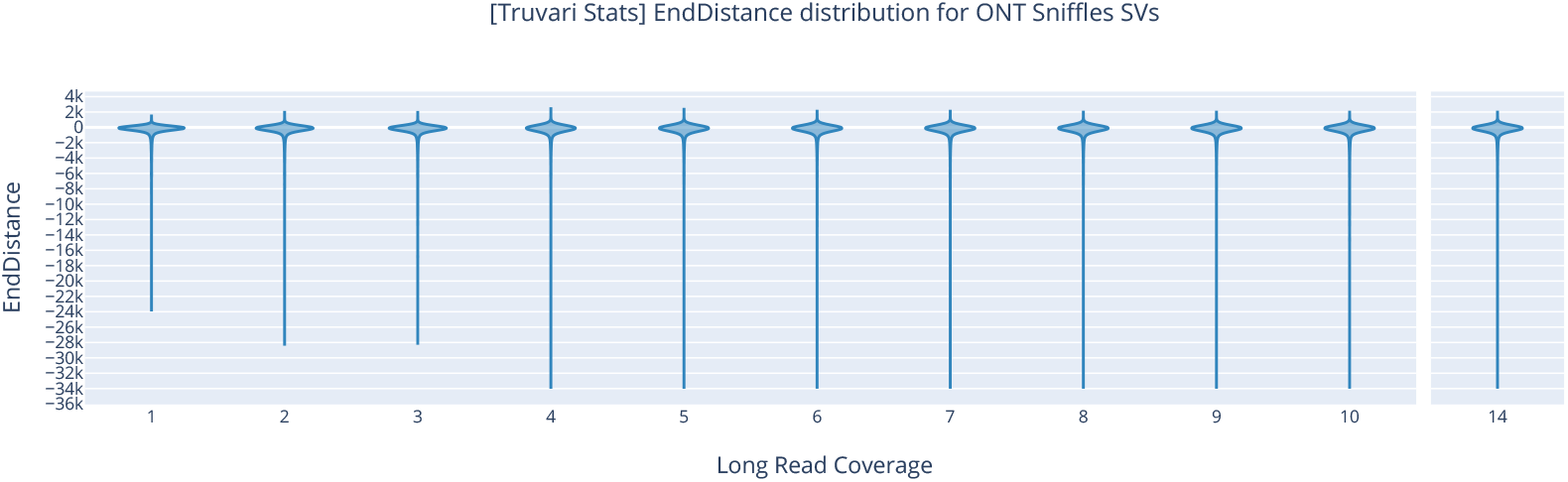
Distribution of Truvari’s “EndDistance” metric for ONT reads. This measures how far the input VCF’s SVs’ end positions are relative to the end positions in the truth VCF for TP matches. We observe a relative consistency across long-read coverages, matching the pattern of the high-coverage control, suggesting the distribution captured at low coverage is reflective of the one at high coverage.

**Supplementary Fig. 28.**
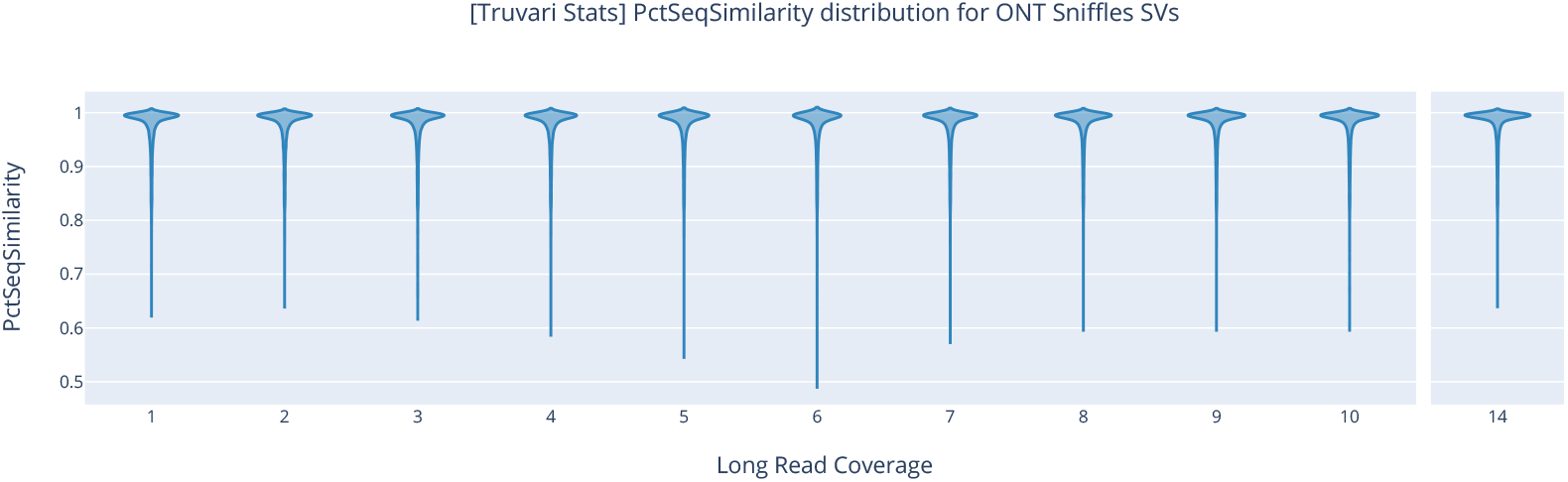
Distribution of Truvari’s “PctSeqSimilarity” metric for ONT reads. This measures the relative similarity of allele sequences in the input VCF compared to the sequences in the truth VCF for TP matches. We observe a relative consistency across long-read coverages, matching the pattern of the high-coverage control, suggesting the distribution captured at low coverage is reflective of the one at high coverage.

#### B.2.4. Short-Read Regenotyping of Structural Variants

**Supplementary Fig. 29.**
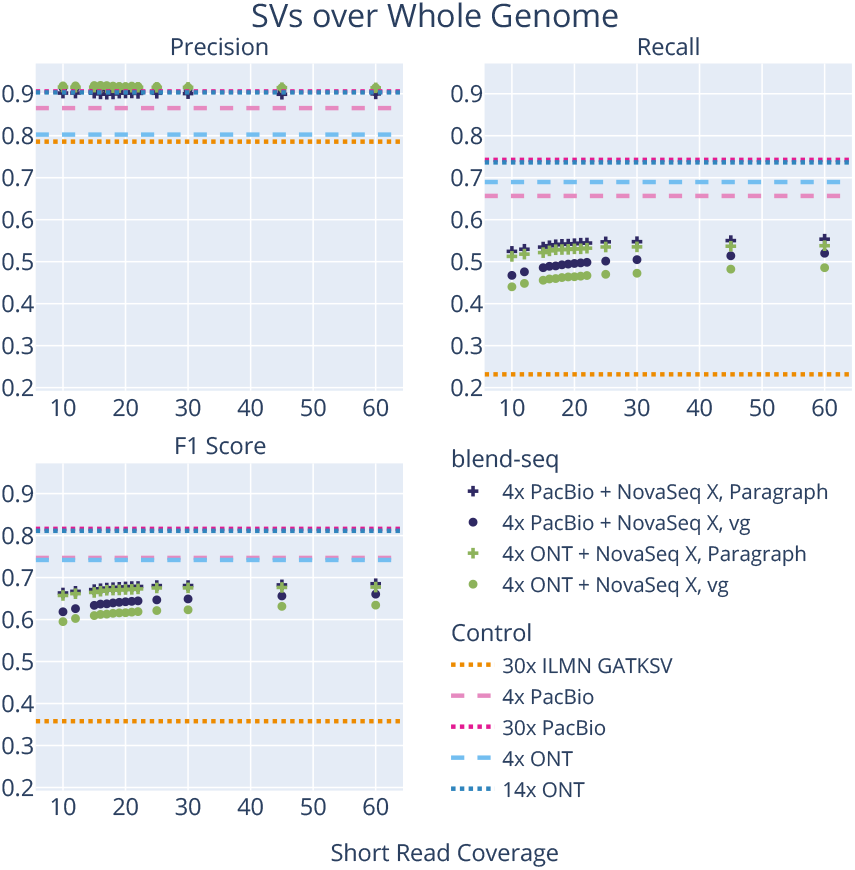
Short-read SV genotyper performance for SVs overall. Although the genotypers improve SV precision overall, it comes at a large cost in decreasing recall, leading to a net decrease in performance compared to the 4x longread controls.

**Supplementary Fig. 30.**
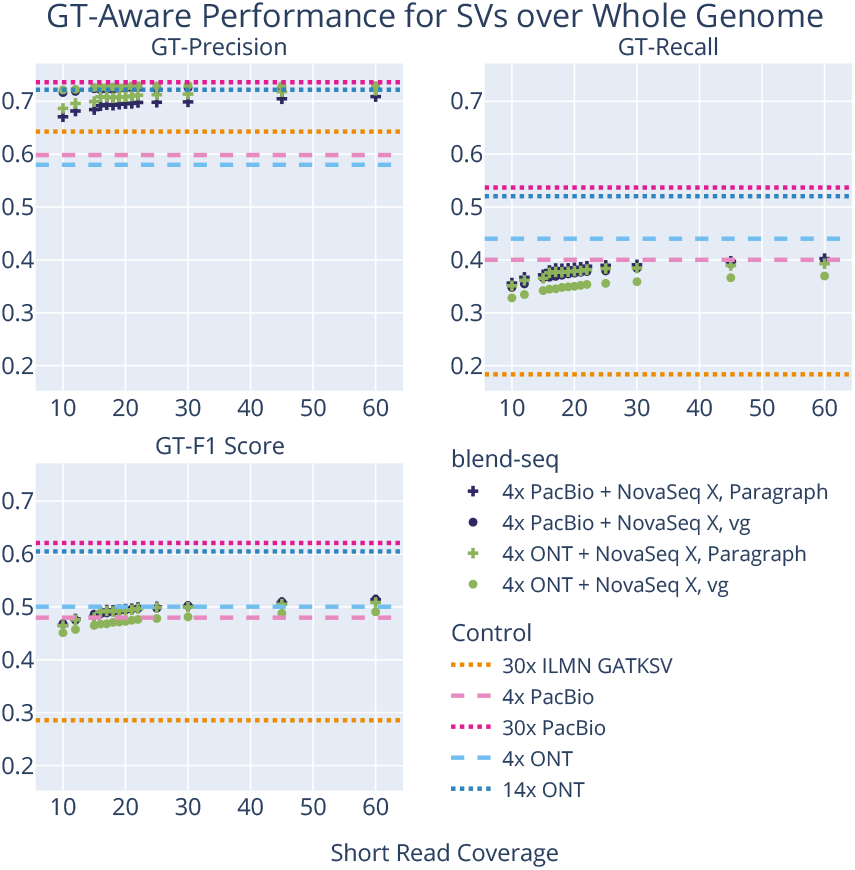
Short-read SV genotyper GT-aware performance for SVs overall. Using the short-read genotypers leads to a precision-recall trade-off overall, improving GT-Precision but decreasing GT-Recall, which leads to similar GT-F1 Scores when compared to the 4x long-read controls.

**Supplementary Fig. 31.**
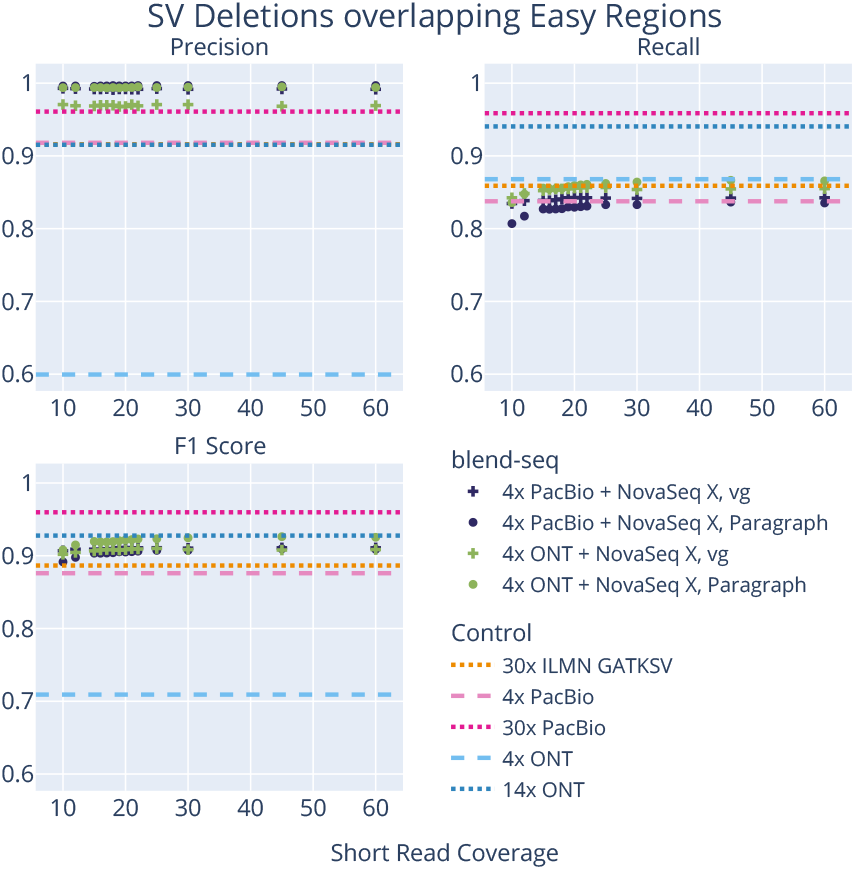
Short-read SV genotyper performance for SV deletions in easy regions. Precision for SV deletions increases over the 4x long-read controls for the blend-seq experiments when using the genotypers, without sacrificing much recall, especially when short read coverage is at least 15x. This leads to a net increase in F1 scores for SV deletions in easy regions compared to the 4x long-read controls.

**Supplementary Fig. 32.**
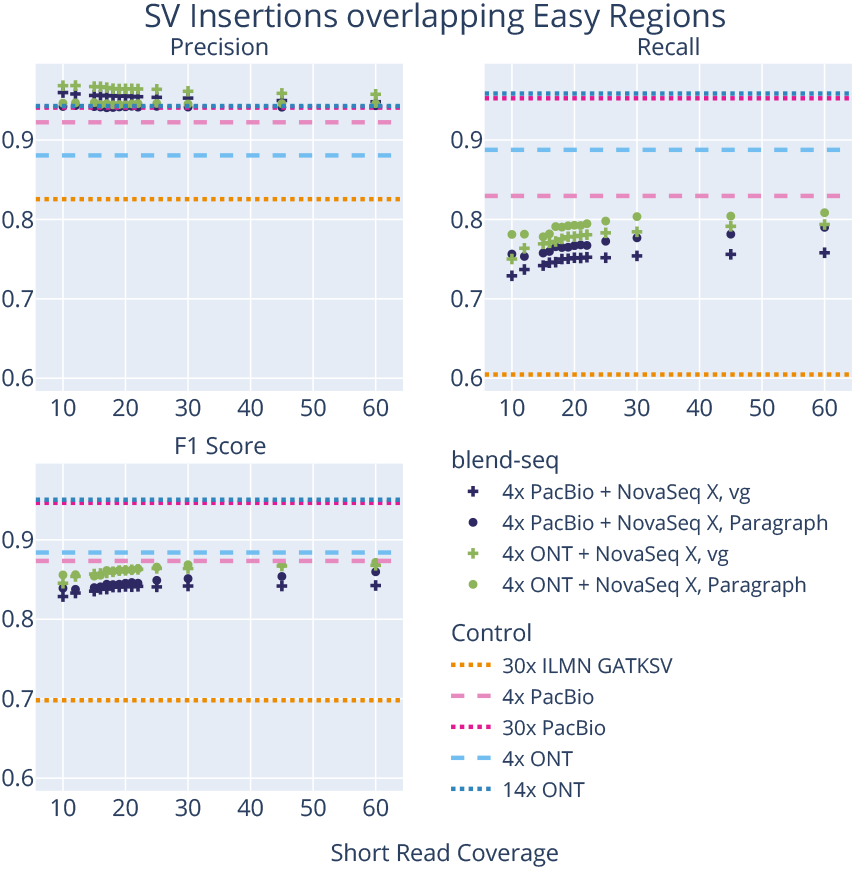
Short-read SV genotyper performance for SV insertions in easy regions. Precision for SV insertions increases over the 4x long-read controls for the blend-seq experiments when using the genotypers, but comes at the cost of losing recall. This leads to a net decrease in F1 scores for SV insertions in easy regions compared to the 4x long-read controls.

**Supplementary Fig. 33.**
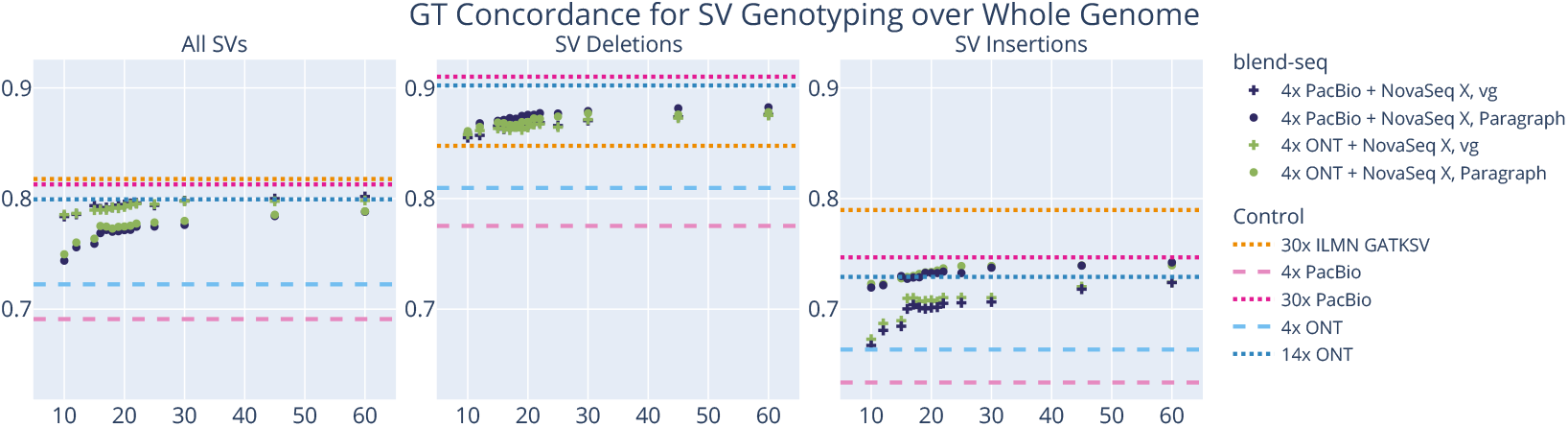
GT concordance performance by short-read coverage over the whole genome. The blend-seq experiments outperform the low-depth long-read controls and the short-read controls for SV deletions, but lag behind for SV insertions. Note that GT concordance is defined as the number of correctly genotyped true positive calls divided by the total number of true positive calls. This means the short reads may better genotype the insertions they are capable of calling, which are likely to be “easier” and occur at a much lower quantity than blend-seq.

**Supplementary Fig. 34.**
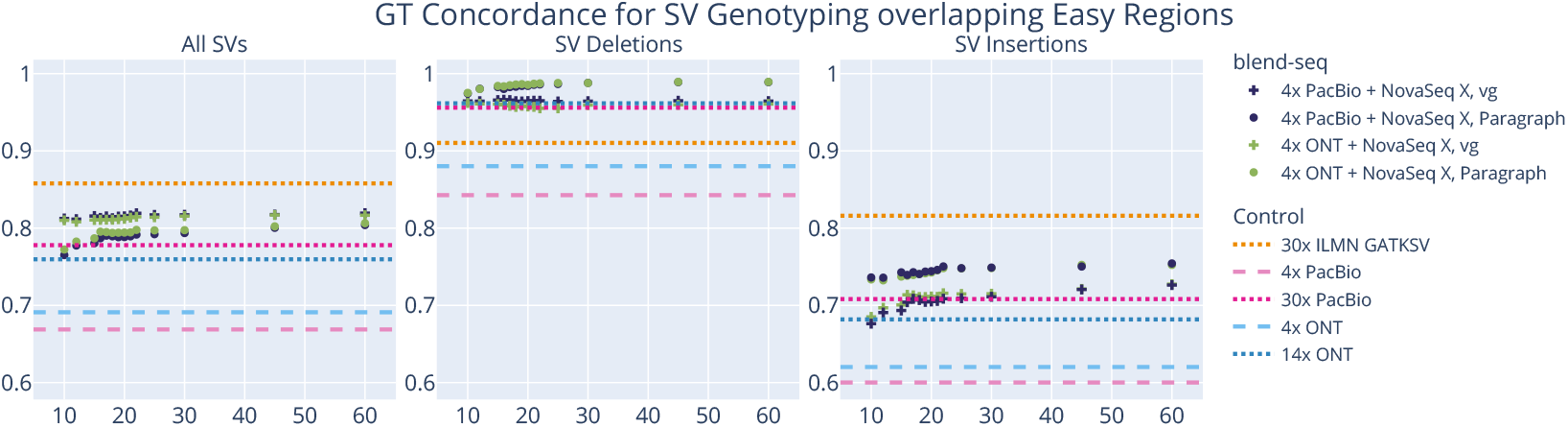
GT concordance performance by short-read coverage over easy regions. The blend-seq experiments outperform all others in SV deletions, but fall behind the short-read control for insertions. Note that GT concordance is defined as the number of correctly genotyped true positive calls divided by the total number of true positive calls. This means the short reads may better genotype the insertions they are capable of calling, which are likely to be “easier” and occur at a much lower quantity than blend-seq.

**Supplementary Fig. 35.**
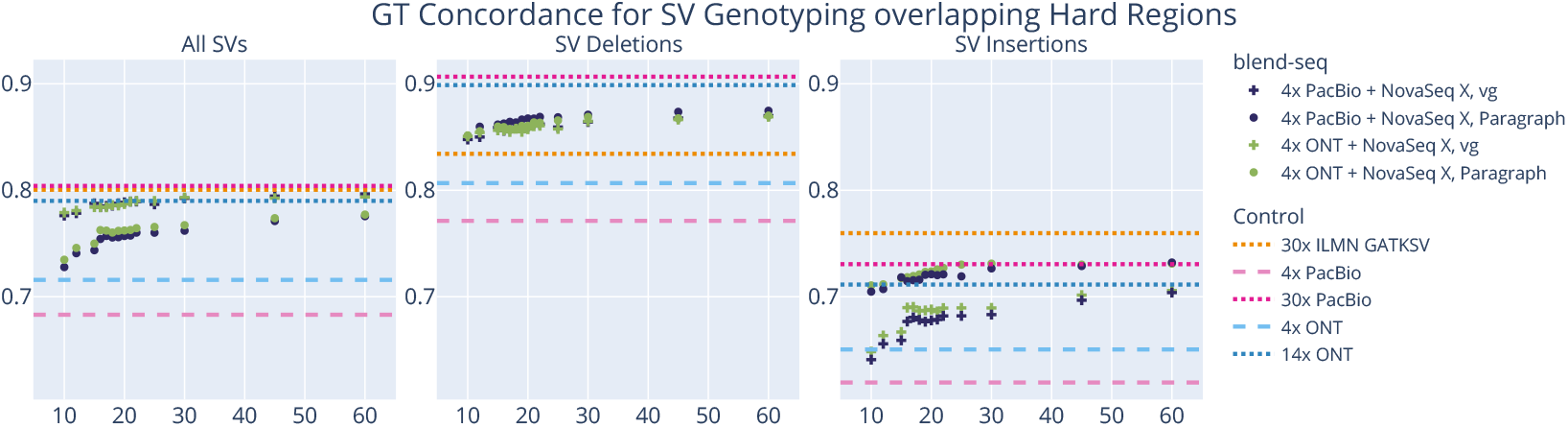
GT concordance performance by short-read coverage over hard regions. The blend-seq experiments outperform the low-depth long-read controls and the short-read controls for SV deletions, but lag behind for SV insertions. Note that GT concordance is defined as the number of correctly genotyped true positive calls divided by the total number of true positive calls. This means the short reads may better genotype the insertions they are capable of calling, which are likely to be “easier” and occur at a much lower quantity than blend-seq.

### B.3. Extra Materials for Phasing Performance

**Supplementary Fig. 36.**
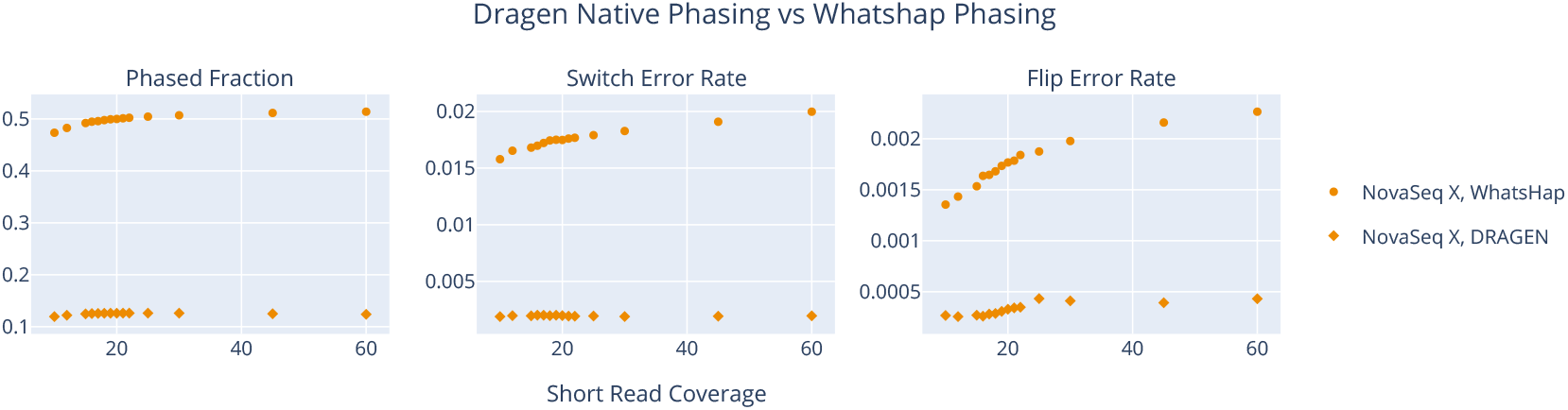
Comparing DRAGEN native phasing performance to using WhatsHap phase. We see that WhatsHap phases 50% of the heterozygous variants (compared to DRAGEN’s 12%). In addition, WhatsHap produces hundreds of fully-phased genes compared to the zero fully-phased by DRAGEN. Despite the higher rate of switch and flip errors, these are expected to occur more often with longer phase blocks, and the statistics for WhatsHap demonstrate a better attempt at phasing using the read data than the DRAGEN outputs.

**Supplementary Fig. 37.**
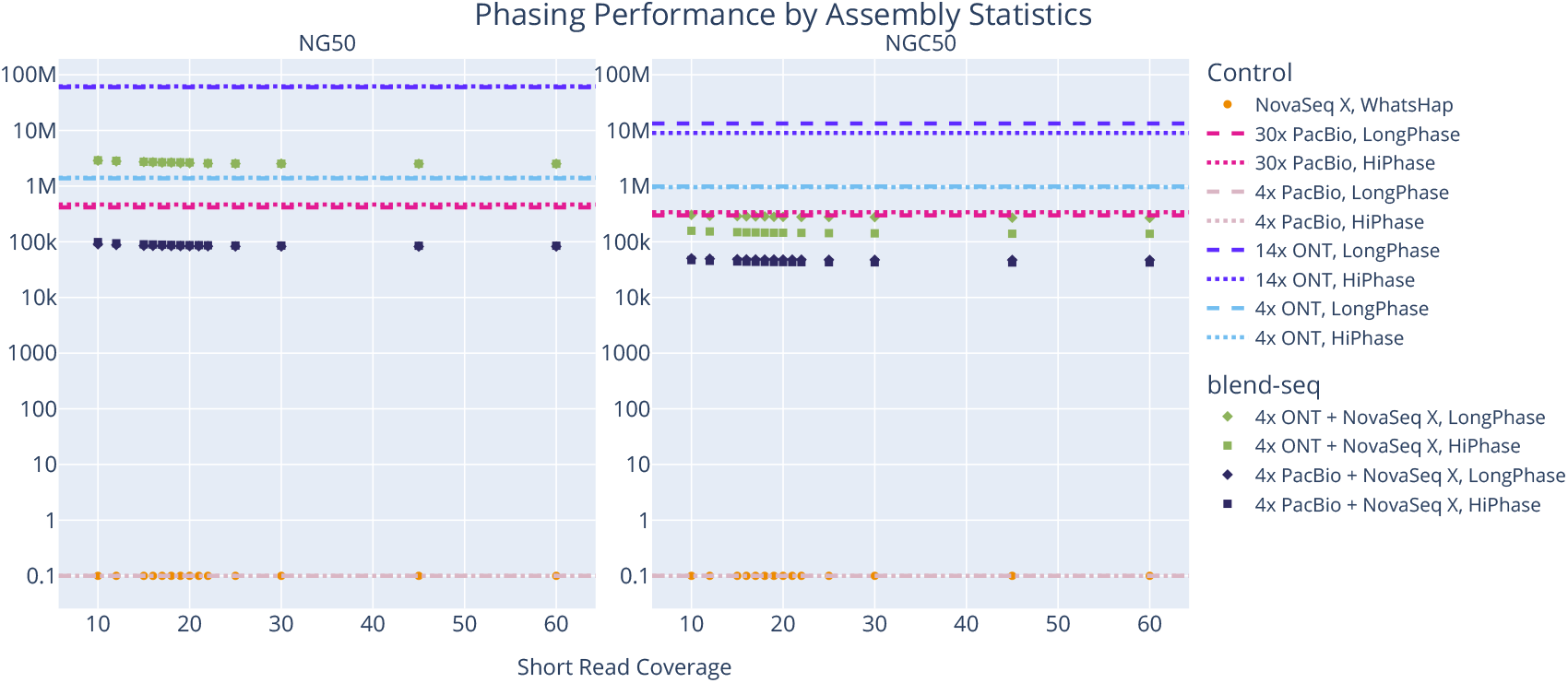
Comparing LongPhase and HiPhase for phasing performance by assembly statistics. We see very little difference between the tools overall, except for a slight advantage to LongPhase when comparing the NGC50 stats, especially when using ONT reads.

**Supplementary Fig. 38.**
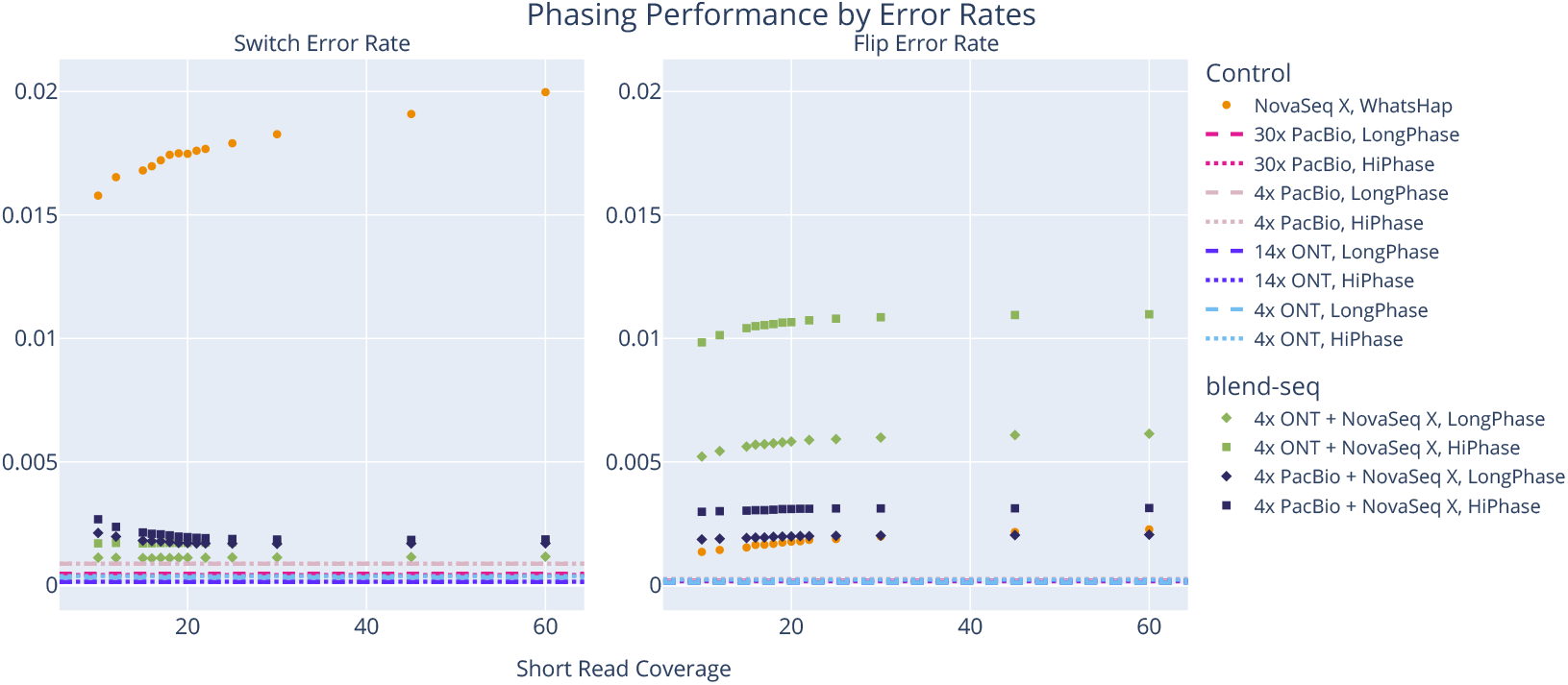
Comparing LongPhase and HiPhase for phasing performance by error rates. We see LongPhase produces fewer switch errors for the blend-seq experiments especially at lower short-read coverages. There is also a significant drop in flip errors for the blend-seq experiments, particularly for the ONT hybrids, justifying the choice of LongPhase for the pipeline.

**Supplementary Fig. 39.**
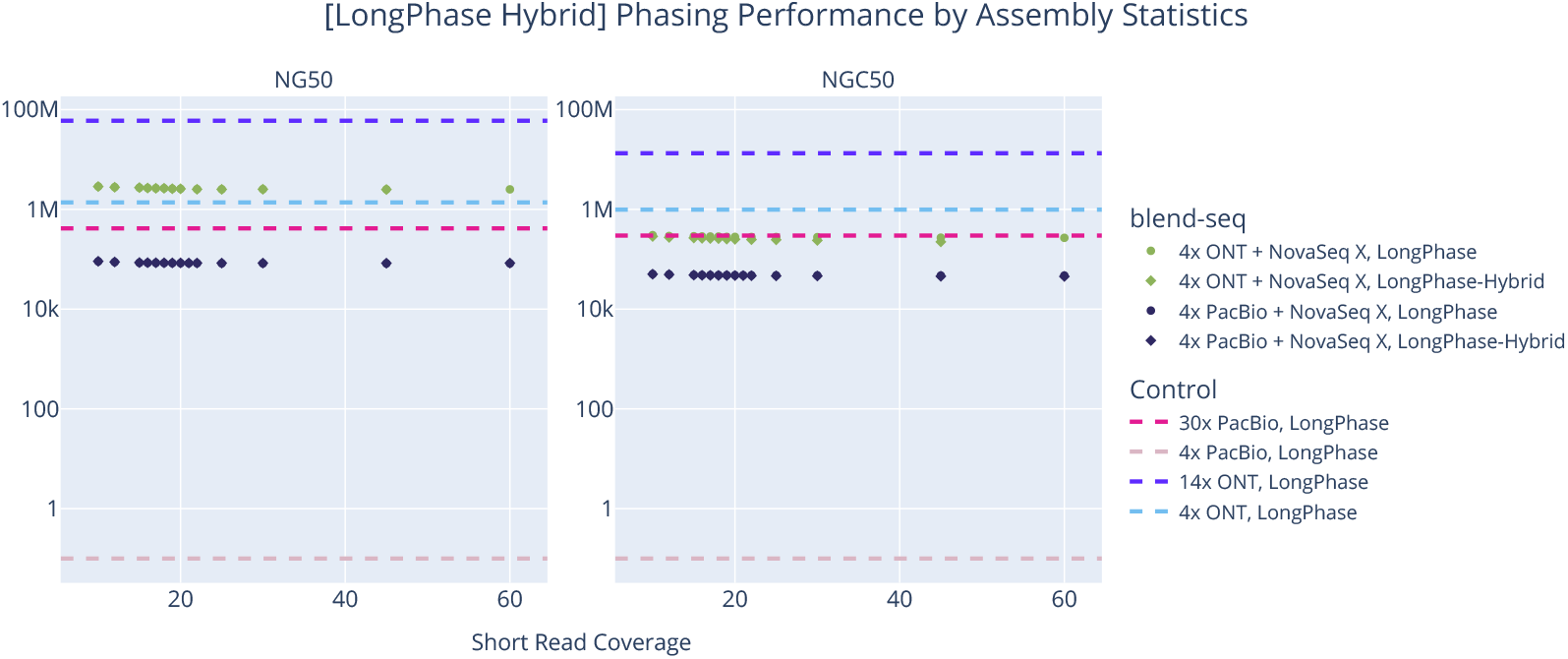
LongPhase hybrid mode assembly phasing statistics comparison. Using the LongPhase hybrid mode, we see that the NG50 is mostly unchanged. However, the additional switch errors added (see Supplementary Fig. 40) causes the NGC50 to drop slightly. Values of 0 are rescaled to 0.1 to be visible on the log-scaled y-axis.

**Supplementary Fig. 40.**
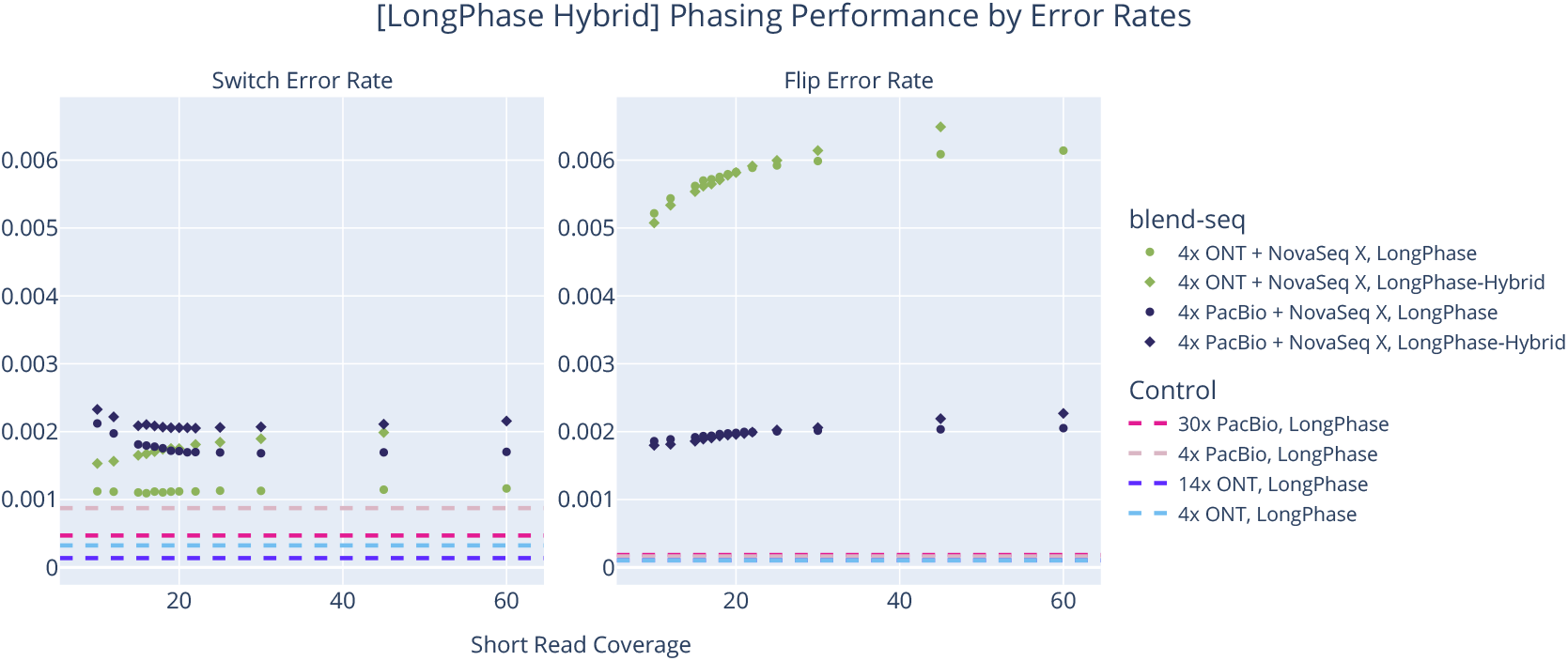
LongPhase hybrid mode phasing error rates comparison. Using the LongPhase hybrid mode, we see that it provides more phased variants, but at the cost of significantly more switch errors. This supports the fact that the short reads are adding extra shorter blocks or extending existing ones but with a higher degree of inaccuracy.

**Supplementary Fig. 41.**
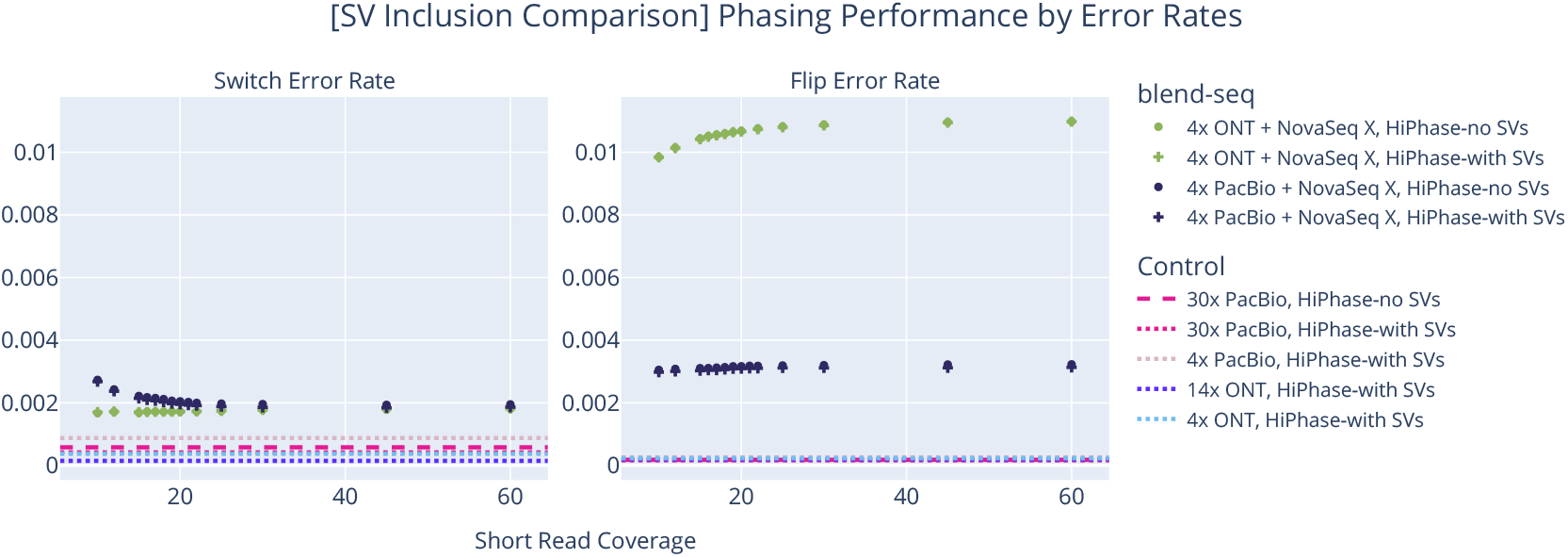
Comparison of phasing performance when including or excluding SVs from phasing process. We investigated the effects on phasing performance when including or excluding SVs from the input VCF for HiPhase (no difference was observed for LongPhase). There is very little difference in the error rates, but it is still a small improvement when including the SVs (especially when using the PacBio reads).

#### B.3.1. Long-Read-Only Phasing

This subsection contains some plots of phasing performance stratified by varying long-read coverage, with no short reads used. Variant calls were made using just the long reads with the same processes used to generate the long-read-only variant calls for controls as described in the Methods section.

**Supplementary Fig. 42.**
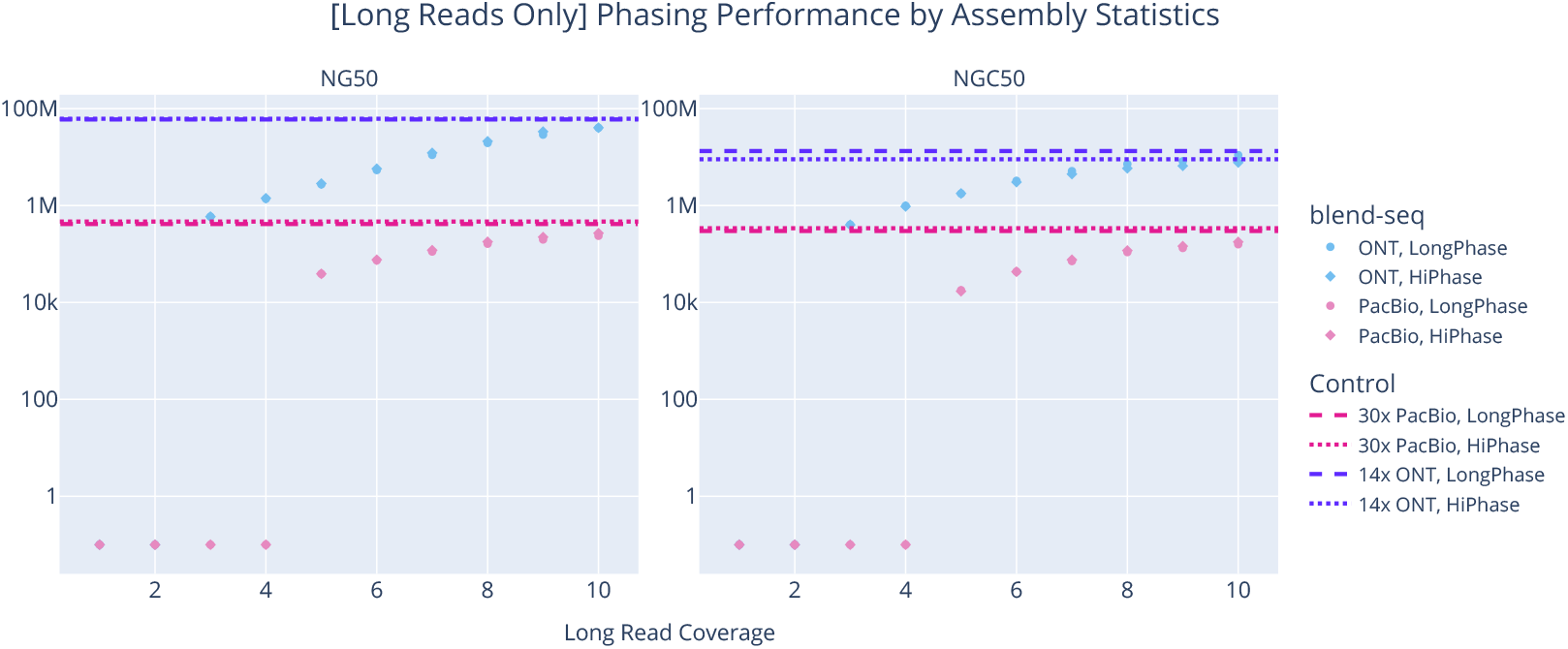
Comparison of phasing assembly statistics performance using long reads alone at varying coverages. On the log-scaled shared axis, the NG50 and NGC50 statistics generally increase linearly in each layer of PacBio reads and expontentially for each layer of ONT reads, reaching close to saturation by 10x. Values of 0 are rescaled to 0.1 to be visible on the log-scaled y-axis.

**Supplementary Fig. 43.**
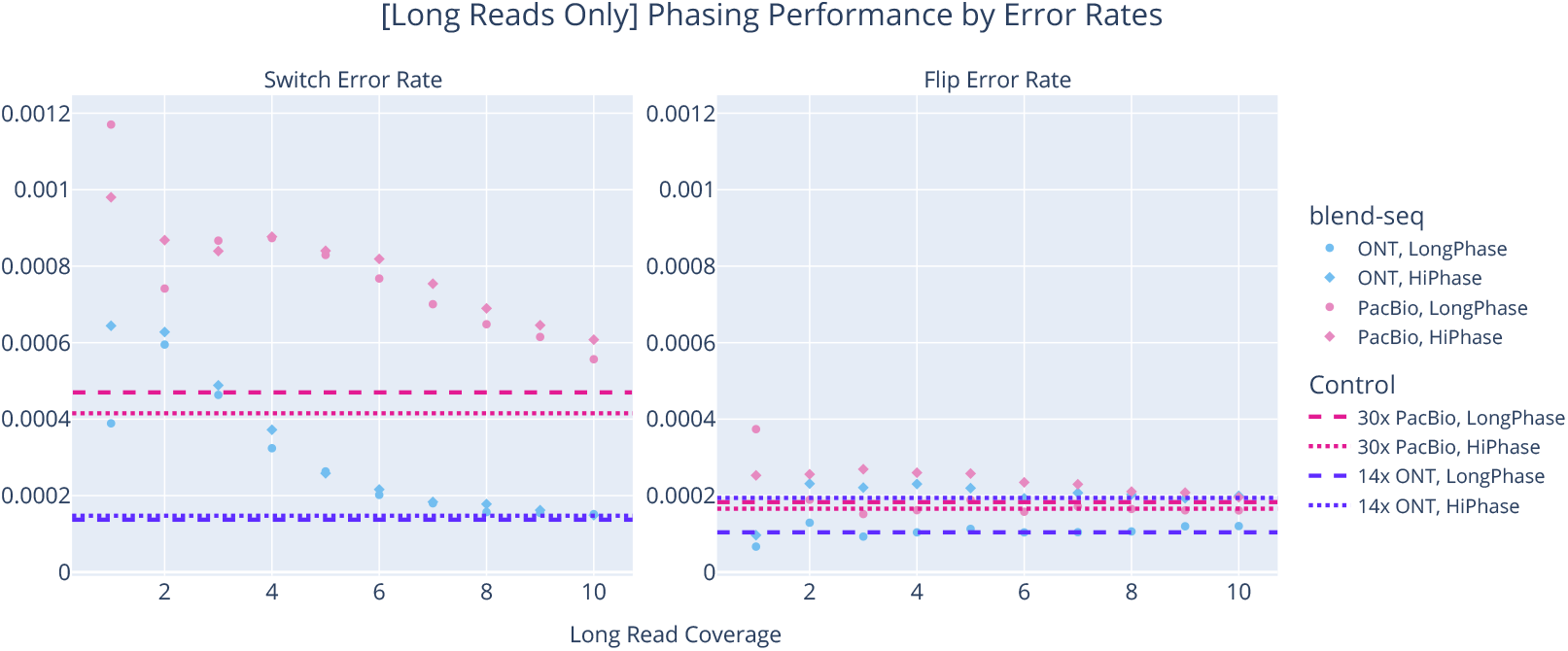
Comparison of phasing error rates using long reads alone at varying coverages. The switch error rate clearly decreases with extra layers of long reads, while the flip error rate follows a less defined trend either slightly downward (HiPhase) or roughly stable (LongPhase).

